# Genetic effect estimates in case-control studies when a continuous variable is omitted from the model

**DOI:** 10.1101/756015

**Authors:** Ying Sheng, Chiung-Yu Huang, Siarhei Lobach, Lydia Zablotska, Iryna Lobach, for the Alzheimer’s Disease Neuroimaging Initiative

## Abstract

Large-scale genome-wide analyses scans provide massive volumes of genetic variants on large number of cases and controls that can be used to estimate the genetic effects. Yet, the sets of non-genetic variables available in publicly available databases are often brief. It is known that omitting a continuous variable from a logistic regression model can result in biased estimates of odds ratios (OR) (e.g., Gail et al (1984), Neuhaus et al (1993), Hauck et al (1991), Zeger et al (1988)). We are interested to assess what information is needed to recover the bias in the OR estimate of genotype due to omitting a continuous variable in settings when the actual values of the omitted variable are not available. We derive two estimating procedures that can recover the degree of bias based on a conditional density of the omitted variable or knowing the distribution of the omitted variable. Importantly, our derivations show that omitting a continuous variable can result in either under- or over-estimation of the genetic effects. We performed extensive simulation studies to examine bias, variability, false positive rate, and power in the model that omits a continuous variable. We show the application to two genome-wide studies of Alzheimer’s disease.

**Data Availability Statement:** The data that support the findings of this study are openly available in the Database of Genotypes and Phenotypes at [https://www.ncbi.nlm.nih.gov/projects/gap/cgibin/study.cgi?study_id=phs000372.v1.p1], reference number [phs000372.v1.p1] and at the Alzheimer’s Disease Neuroimaging Initiative http://adni.loni.usc.edu/.

## INTRODUCTION

Recent advances in genotyping technology generated volumes and variety of datasets that are archived in massive publicly available databases (e.g. the Database of Genotypes and Phenotypes https://www.ncbi.nlm.nih.gov/gap/ the Cancer Genome Athlas https://portal.gdc.cancer.gov/, the UK Biobank https://www.ukbiobank.ac.uk/). These data provide valuable information that can be analyzed to improve our understanding about the genetic predisposition to complex diseases, such as cancer, diabetes, neurodegenerative disease. Such analyses of association might serve multiple purposes one of which is to identify the genetic variants and rank them according to strength of the evidence for an association with the complex diseases. As the result we might obtain valuable clues to the underlying aeteologic mechanisms of complex diseases. A commonly overseen complication is that omitting a variable from a logistic regression model can substantially bias the genetic effect estimates. We are interested to derive what types of information are needed to recover bias in settings when the actual values of the omitted variable are not available to the researcher.

From the statistical literature (Gail et al (1984), Neuhaus et al (1993), Hauck et al (1991), Zeger et al (1988)) we know that omitting variables associated with the disease can cause bias in the odds ratio (OR) estimates, because the OR estimates reflect both the effect size and variability in the error terms. Gail et al (1984), Neuhaus et al (1993), Zeger et al (1988) derive the magnitude of bias in the estimate that is a function of the OR of the omitted variable and the distribution of the omitted variable.

Because the correct OR estimates of the omitted variable, i.e. the estimates from the full model that includes both the genetic effects and the omitted variable, are rarely available in the literature, we are interested to examine what other types of information are needed for a researcher to be able to correct the bias. We are also interested to assess what determines the directionality of the bias.

The setting we consider is unique. The model with an omitted variable is misspecified for three reasons. First, the data are collected using retrospective design where the cases and controls are sampled from their populations, while the data are analyzed in a prospective logistic regression model. As pointed out in the seminal work by Prentice and Pyke (1979), we know that this aspect of misspecification does not result in bias of OR estimates because the OR can be estimated consistently from retrospective likelihood-based methods. Secondly, model is misspecified because the variable is omitted from the model, what also results in the third misspecification, namely that if the true risk function is logistic, the link between the other variables and risk of the disease with omitted variable might not be logistic.

The setting we consider is also unique in that usually the magnitude of the effect of a genetic variant is estimated to be small to moderate, i.e. the range of effect sizes somewhere between -log(1.5) and log(1.5) (Park et al, 2011). A few exceptions, however, have been noted in the literature. For example, in the context of Alzheimer’s disease, ApoE genotype is estimated to have OR of 3.1 for heterozygous *ε*4 genotype and 34.3 for homozygous *ε*4 genotype (Kukull et al, 1996).

Our paper is organized as follows. We first perform a series of simulation studies to assess the problem empirically. The simulations are described in the Assessment of the Problem section. Next, in the Estimates of the Reduced vs. Full Models section we derive the relationships between parameters of the reduced model where the variable is omitted and the parameters in the full model where the variable is included. We further conduct simulation studies described in Simulation Studies section to assess how various pieces of information can contribute to recovery of the bias. We show the application to the studies of Alzheimer’s disease. And we conclude the paper by a brief discussion.

## MATERIALS AND METHODS

### ASSESSMENT OF THE PROBLEM

We first perform a series of simulation studies to assess potential bias, variance, mean squared error (MSE), false discovery rate (FDR), and power reduction due to omitting a continuous variable that is associated with the disease status. We assume that the omitted variable *O* and the genotype *G* are distributed independently in the population.

#### Setting 1

We first examine models with one genetic variant. We simulate the genetic variant from Bernoulli(0.1) and an omitted variable *O* from Normal(0, *σ*^2^). We set *σ* = 1,2 and next simulate the disease status according to the full disease risk model

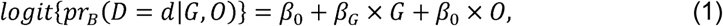

where we let *β*_0_ = −1, −5; *β*_G_ = log(1), log(1.5), log(2), log(2.5), log(3), log(5), log(8), and *β*_*O*_ = log(1), log(1.5), log(2), log(2.5), log(3), log(5), log(8) across various settings. Generate 5,000 samples of 3,000/10,000 cases and 3,000/10,000 controls using retrospective/case-control design.

We next estimate the parameters based on the reduced (and hence misspecified) logistic regression model

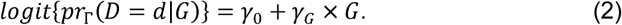

Shown in **Supplementary Table 1** are probability of the disease in the population, bias in 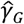 as the estimates of *β*_*G*_ = 0, variance, MSE, and FDR. The estimates in this setting are nearly unbiased with FDR that are nominal. Shown in **Table 1** is the setting when *β*_*G*_ = log (1.5). Here bias becomes more pronounced what also reduces the power to detect an effect. For example, when *β*_*O*_ = log(3) = 1.0986, *β*_0_ = −1, *σ* = 2, bias in 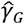 as the estimate of *β*_*G*_ is -0.18, while power to detect the effect is 0.76. Frequency of the disease in the population is 0.37. Shown in **Table 2** and **Supplementary Table 2** are the settings when *β*_*G*_ = log(2), log(2.5), log(3), log(5), log(8). Biases increase as the magnitude of the coefficient increases, however the bias because of its direction does not have impact on power to detect the effect. As illustrated in **Supplementary Table 3** the biases noted in samples with 3,000 cases and 3,000 controls persist in samples with 10,000 cases and 10,000 controls.

**Table 1:** Bias, variance and mean square error (MSE) of genetic effect estimates obtain using reduced model (2) when the data are simulated using full model (1). Shown is also probability of the disease in the population, i.e. *pr*_*B*_ (*D* = 1), and false discovery rate (FDR). The genotype is simulated to be Bernoulli(0.1), the omitted variable is simulated from Normal(0, *σ*^2^). We simulated the disease status from model (1) with parameters *β*_0_ = −1, −5; *β*_*G*_ = log(1.5), *β*_*O*_ = log(1), log(1.5), log(2), log(2.5), log(3), log(5), log(8). The results are based on 5,000 datasets of 3,000 cases and 3,000 controls.

|  |  |  |  |  |  |
| --- | --- | --- | --- | --- | --- |
| $\beta_o = \log(1)$ | | | | | |
| $(\beta_o, \sigma)$ | $pr_B(D = 1)$ | Bias | Variance | MSE | power |
| (-1,1) | 0.28 | 0.008 | 0.007 | 0.007 | 1 |
| (-1,2) | 0.28 | 0.008 | 0.007 | 0.007 | 1 |
| (-5,1) | 0.007 | -0.03 | 0.005 | 0.006 | 1 |
| (-5,2) | 0.007 | -0.03 | 0.005 | 0.006 | 1 |
| $\beta_o = \log(1.5) = 0.4055$ | | | | | |
| $(\beta_o, \sigma)$ | $pr_B(D = 1)$ | Bias | Variance | MSE | power |
| (-1,1) | 0.28 | -0.009 | 0.007 | 0.007 | 1 |
| (-1,2) | 0.30 | -0.05 | 0.007 | 0.01 | 1 |
| (-5,1) | 0.008 | -0.01 | 0.006 | 0.006 | 1 |
| (-5,2) | 0.01 | -0.03 | 0.006 | 0.006 | 1 |
| $\beta_o = \log(2) = 0.6931$ | | | | | |
| $(\beta_o, \sigma)$ | $pr_B(D = 1)$ | Bias | Variance | MSE | power |
| (-1,1) | 0.30 | -0.03 | 0.008 | 0.009 | 0.99 |
| (-1,2) | 0.33 | -0.10 | 0.007 | 0.02 | 0.95 |
| (-5,1) | 0.02 | -0.03 | 0.006 | 0.007 | 1 |
| (-5,2) | 0.03 | -0.03 | 0.006 | 0.007 | 1 |
| $\beta_o = \log(2.5) = 0.9163$ | | | | | |
| $(\beta_o, \sigma)$ | $pr_B(D = 1)$ | Bias | Variance | MSE | power |
| (-1,1) | 0.31 | -0.06 | 0.007 | 0.01 | 0.98 |
| (-1,2) | 0.35 | -0.15 | 0.008 | 0.03 | 0.86 |
| (-5,1) | 0.01 | -0.02 | 0.006 | 0.006 | 1 |
| (-5,2) | 0.03 | -0.06 | 0.006 | 0.01 | 0.99 |
| $\beta_o = \log(3) = 1.0986$ | | | | | |
| $(\beta_o, \sigma)$ | $pr_B(D = 1)$ | Bias | Variance | MSE | power |
| (-1,1) | 0.32 | -0.07 | 0.007 | 0.01 | 0.98 |
| (-1,2) | 0.37 | -0.18 | 0.007 | 0.04 | 0.76 |
| (-5,1) | 0.01 | -0.01 | 0.006 | 0.006 | 1 |
| (-5,2) | 0.04 | -0.11 | 0.006 | 0.02 | 0.96 |
| $\beta_o = \log(5) = 1.6094$ | | | | | |
| $(\beta_o, \sigma)$ | $pr_B(D = 1)$ | Bias | Variance | MSE | power |
| (-1,1) | 0.34 | -0.13 | 0.007 | 0.02 | 0.92 |
| (-1,2) | 0.40 | -0.23 | 0.007 | 0.06 | 0.54 |
| (-5,1) | 0.02 | -0.04 | 0.006 | 0.007 | 1 |
| (-5,2) | 0.09 | -0.19 | 0.007 | 0.04 | 0.73 |
| $\beta_o = \log(8) = 2.0794$ | | | | | |
| $(\beta_o, \sigma)$ | $pr_B(D = 1)$ | Bias | Variance | MSE | power |
| (-1,1) | 0.36 | -0.17 | 0.007 | 0.03 | 0.80 |
| (-1,2) | 0.42 | -0.27 | 0.007 | 0.08 | 0.37 |
| (-5,1) | 0.04 | -0.10 | 0.007 | 0.02 | 0.97 |
| (-5,2) | 0.14 | -0.25 | 0.007 | 0.07 | 0.48 |

**Table 2:**
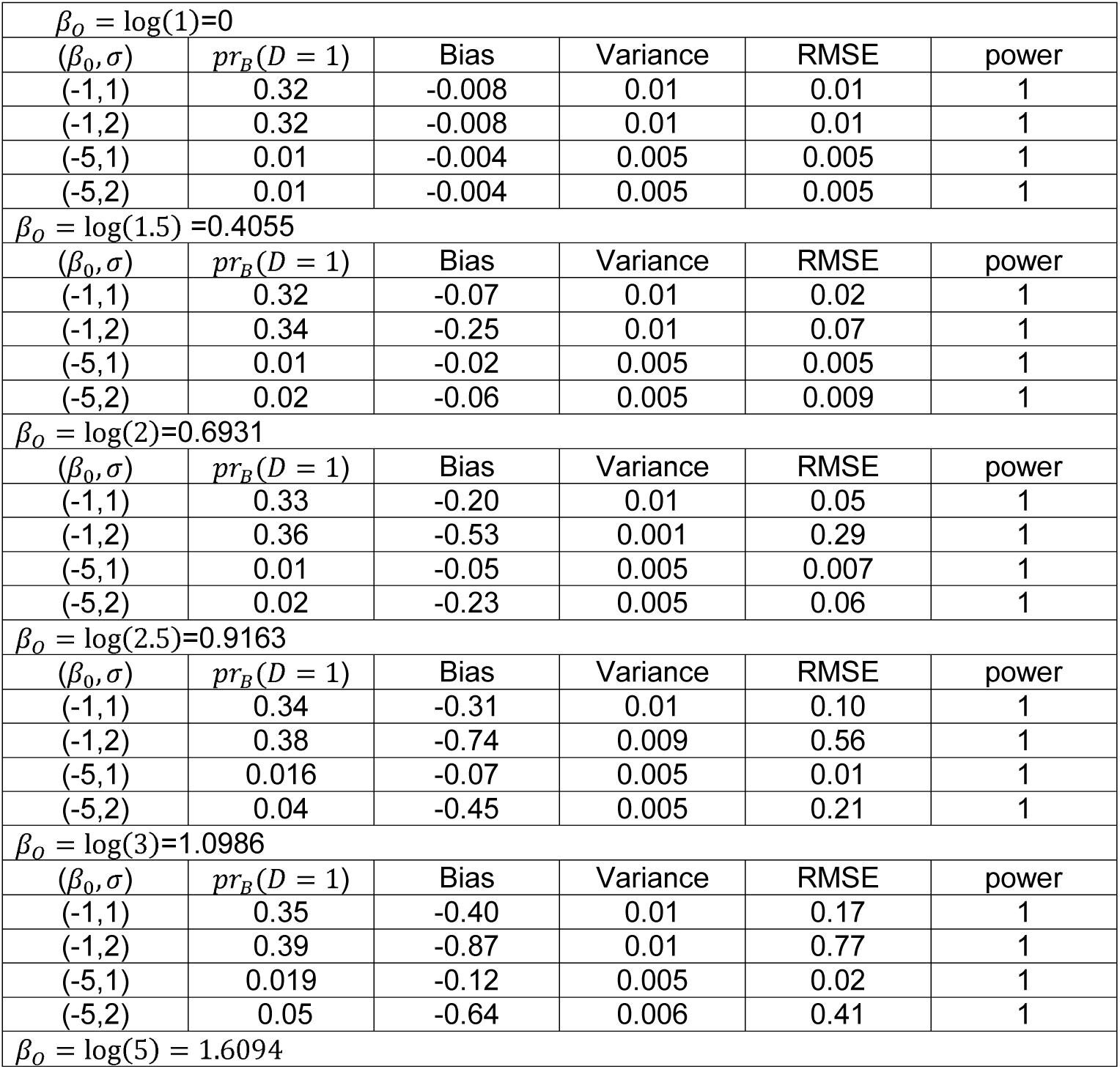
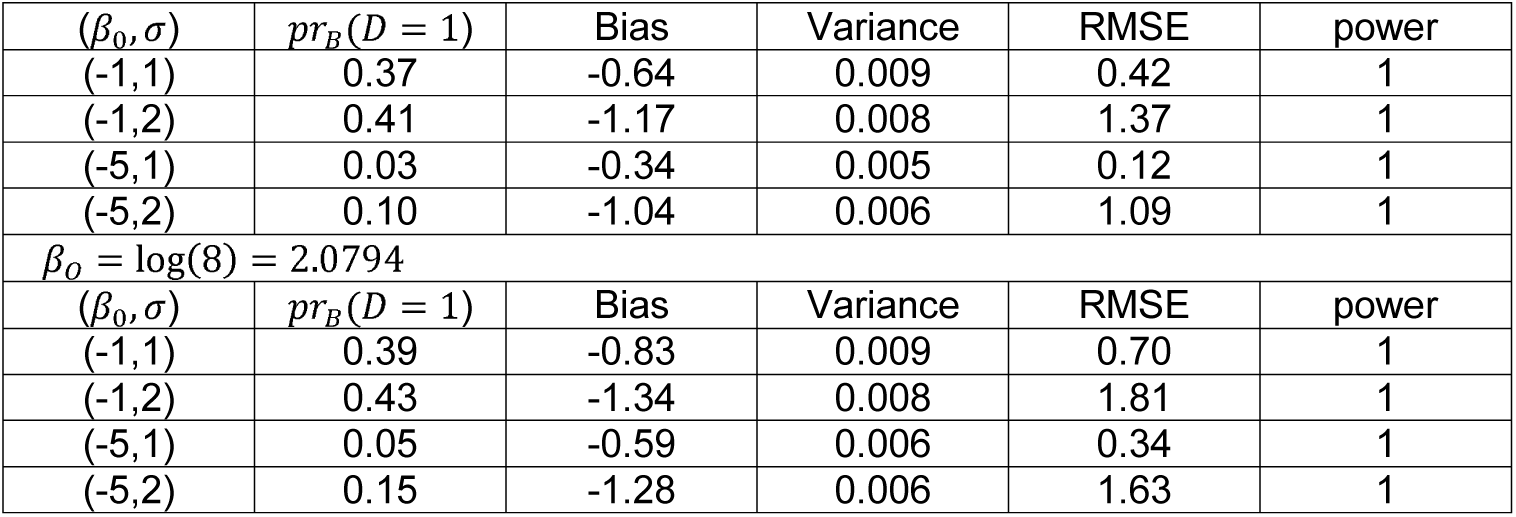
Bias, variance and mean square error (MSE) of genetic effect estimates obtain using reduced model (2) when the data are simulated using full model (1). Shown is also probability of the disease in the population, i.e. *pr*_*B*_ (*D* = 1), and false discovery rate (FDR). The genotype is simulated to be Bernoulli(0.1), the omitted variable is simulated from Normal(0, *σ*^2^). We simulated the disease status from model (1) with parameters *β*_0_= −1, −5; *β*_*G*_ = log(8), *β*_*O*_ = log(1), log(1.5), log(2), log(2.5), log(3), log(5), log(8). The results are based on 5,000 datasets of 3,000 cases and 3,000 controls.

|  |  |  |  |  |  |
| --- | --- | --- | --- | --- | --- |
| $\beta_o = \log(1)=0$ | | | | | |
| $(\beta_o, \sigma)$ | $pr_B(D = 1)$ | Bias | Variance | RMSE | power |
| (-1,1) | 0.32 | -0.008 | 0.01 | 0.01 | 1 |
| (-1,2) | 0.32 | -0.008 | 0.01 | 0.01 | 1 |
| (-5,1) | 0.01 | -0.004 | 0.005 | 0.005 | 1 |
| (-5,2) | 0.01 | -0.004 | 0.005 | 0.005 | 1 |
| $\beta_o = \log(1.5)=0.4055$ | | | | | |
| $(\beta_o, \sigma)$ | $pr_B(D = 1)$ | Bias | Variance | RMSE | power |
| (-1,1) | 0.32 | -0.07 | 0.01 | 0.02 | 1 |
| (-1,2) | 0.34 | -0.25 | 0.01 | 0.07 | 1 |
| (-5,1) | 0.01 | -0.02 | 0.005 | 0.005 | 1 |
| (-5,2) | 0.02 | -0.06 | 0.005 | 0.009 | 1 |
| $\beta_o = \log(2)=0.6931$ | | | | | |
| $(\beta_o, \sigma)$ | $pr_B(D = 1)$ | Bias | Variance | RMSE | power |
| (-1,1) | 0.33 | -0.20 | 0.01 | 0.05 | 1 |
| (-1,2) | 0.36 | -0.53 | 0.001 | 0.29 | 1 |
| (-5,1) | 0.01 | -0.05 | 0.005 | 0.007 | 1 |
| (-5,2) | 0.02 | -0.23 | 0.005 | 0.06 | 1 |
| $\beta_o = \log(2.5)=0.9163$ | | | | | |
| $(\beta_o, \sigma)$ | $pr_B(D = 1)$ | Bias | Variance | RMSE | power |
| (-1,1) | 0.34 | -0.31 | 0.01 | 0.10 | 1 |
| (-1,2) | 0.38 | -0.74 | 0.009 | 0.56 | 1 |
| (-5,1) | 0.016 | -0.07 | 0.005 | 0.01 | 1 |
| (-5,2) | 0.04 | -0.45 | 0.005 | 0.21 | 1 |
| $\beta_o = \log(3)=1.0986$ | | | | | |
| $(\beta_o, \sigma)$ | $pr_B(D = 1)$ | Bias | Variance | RMSE | power |
| (-1,1) | 0.35 | -0.40 | 0.01 | 0.17 | 1 |
| (-1,2) | 0.39 | -0.87 | 0.01 | 0.77 | 1 |
| (-5,1) | 0.019 | -0.12 | 0.005 | 0.02 | 1 |
| (-5,2) | 0.05 | -0.64 | 0.006 | 0.41 | 1 |
| $\beta_o = \log(5) = 1.6094$ | | | | | |

**Table 2:** Bias, variance and mean square error (MSE) of genetic effect estimates obtain
| $(\beta_0, \sigma)$ | $pr_B(D = 1)$ | Bias | Variance | RMSE | power |
| --- | --- | --- | --- | --- | --- |
| (-1,1) | 0.37 | -0.64 | 0.009 | 0.42 | 1 |
| (-1,2) | 0.41 | -1.17 | 0.008 | 1.37 | 1 |
| (-5,1) | 0.03 | -0.34 | 0.005 | 0.12 | 1 |
| (-5,2) | 0.10 | -1.04 | 0.006 | 1.09 | 1 |
| $\beta_0 = \log(8) = 2.0794$ | | | | | |
| $(\beta_0, \sigma)$ | $pr_B(D = 1)$ | Bias | Variance | RMSE | power |
| (-1,1) | 0.39 | -0.83 | 0.009 | 0.70 | 1 |
| (-1,2) | 0.43 | -1.34 | 0.008 | 1.81 | 1 |
| (-5,1) | 0.05 | -0.59 | 0.006 | 0.34 | 1 |
| (-5,2) | 0.15 | -1.28 | 0.006 | 1.63 | 1 |

#### Setting 2

We next conduct a simulation experiment to assess if the ratio of the parameters is estimated correctly when a continuous variable is omitted from the model. We simulate one genetic variant *G*_1_ from Bernoulli(0.1) and the other one *G*_2_ from Bernoulli(0.25) and O from Normal(0, *σ*^2^), where *σ* = 1,2. We simulate the true disease status from the logistic model :

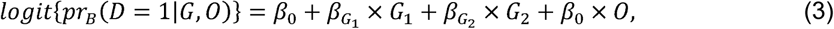

where we let *β*_0_= −1; −5 and we consider various pairs: 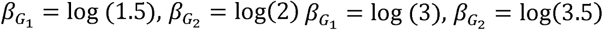; and we let *β*_*O*_ = log(2), log(2.5), log(3), log(5), log(8) across various settings.

#### Setting 3

We now examine a setting with many genetic variables and one omitted environmental variable. The goal of this simulation is to see if the relative order of the genetic variables is estimated correctly. We simulate *G*_1_ … *G*_*M*/2_ from Bernoulli(0.1) and *G*_*M*/2+1_ … *G*_*M*_ from Bernoulli(0.25), and *O* from Normal(0, *σ*^2^), where *σ* = 1,2.

Moreover, we simulate the disease status according to the risk model:

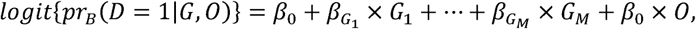

where we let 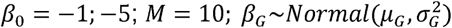, with 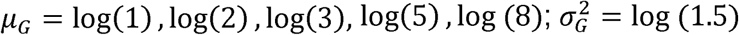 and *β*_*O*_ = log(1), log(1.5), log(2), log(2.5), log(3), log(5), log(8). We next estimate the parameters based on misspecified logistic regression model

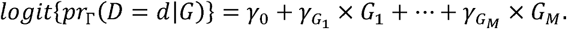

We would like to assess if the order of the genetic effect estimates is preserved. Suppose it does not matter what the magnitude of the estimate is, as long as the relative ordering in maintained. We define the order of the genetic effect by 1) Value of the coefficient; 2) P-value for the coefficient, that is, one ordering will be based just on the value of the coefficient estimate, and the second ordering just based on the p-value. Shown in **Table 3** and **Supplementary Table 4** are the results based on 5,000 samples of 3,000 cases and 3,000 controls. Shown in **Table 3**, the proportion of the genetic variants for which the ranks are the same. As illustrated in **Supplementary Table 4**, the proportion of the genetic variants for which the ranks are the same are very close to 1 when *β*_*O*_ is small.

**Table 3:** Proportions of genetic variants that received the same rank based on the full and reduced genetic models across all variants (ALL), top 10% and top 20%. We simulated 5,000 datasets with 3,000 cases and 3,000 controls. We simulated 10 genetic variants from Bernoulli(0.1) and disease status from the full model with coefficients *β*_*O*_ = log (3) and *µ*_*G*_ = log(1),log(2),log(3),log(5),log (8).

| $\mu_G = \log(1)$ | | | | | | | |
| --- | --- | --- | --- | --- | --- | --- | --- |
| $(\beta_0, \sigma)$ | $pr_B(D = d)$ | Ranks based on $\widehat{OR}$ | | | Ranks based on p values | | |
|  |  | ALL | TOP 10% | TOP 20% | ALL | TOP 10% | TOP 20% |
| (-1,1) | 0.33 | 0.89 | 1 | 0.99 | 0.74 | 0.89 | 0.89 |
| (-1,2) | 0.37 | 0.77 | 0.98 | 0.93 | 0.58 | 0.81 | 0.75 |
| (-5,1) | 0.01 | 0.89 | 1 | 0.99 | 0.75 | 0.89 | 0.87 |
| (-5,2) | 0.05 | 0.79 | 0.98 | 0.95 | 0.57 | 0.69 | 0.71 |
| $\mu_G = \log(2) = 0.693$ | | | | | | | |
| $(\beta_0, \sigma)$ | $pr_B(D = d)$ | Ranks based on $\widehat{OR}$ | | | Ranks based on p values | | |
|  |  | ALL | TOP 10% | TOP 20% | ALL | TOP 10% | TOP 20% |
| (-1,1) | 0.56 | 0.84 | 0.87 | 0.9 | 0.85 | 0.74 | 0.79 |
| (-1,2) | 0.55 | 0.71 | 0.73 | 0.77 | 0.70 | 0.53 | 0.61 |
| (-5,1) | 0.07 | 0.86 | 0.87 | 0.90 | 0.86 | 0.75 | 0.80 |
| (-5,2) | 0.12 | 0.74 | 0.73 | 0.79 | 0.73 | 0.53 | 0.62 |
| $\mu_G = \log(3) = 1.099$ | | | | | | | |
| $(\beta_0, \sigma)$ | $pr_B(D = d)$ | Ranks based on $\widehat{OR}$ | | | Ranks based on p values | | |
|  |  | ALL | TOP 10% | TOP 20% | ALL | TOP 10% | TOP 20% |

**Table 3:** Proportions of genetic variants that received the same rank based on the full
| (-1,1) | 0.66 | 0.82 | 0.85 | 0.88 | 0.87 | 0.92 | 0.95 |
| --- | --- | --- | --- | --- | --- | --- | --- |
| (-1,2) | 0.63 | 0.70 | 0.73 | 0.77 | 0.76 | 0.74 | 0.83 |
| (-5,1) | 0.13 | 0.85 | 0.84 | 0.88 | 0.92 | 0.77 | 0.88 |
| (-5,2) | 0.18 | 0.73 | 0.73 | 0.77 | 0.80 | 0.62 | 0.79 |
| $\mu_G = \log(5) = 1.609$ | | | | | | | |
| $(\beta_0, \sigma)$ | $pr_B(D = d)$ | Ranks based on $\widehat{OR}$ | | | Ranks based on p values | | |
|  |  | ALL | TOP 10% | TOP 20% | ALL | TOP 10% | TOP 20% |
| (-1,1) | 0.75 | 0.81 | 0.83 | 0.87 | 0.84 | 0.91 | 0.81 |
| (-1,2) | 0.71 | 0.69 | 0.71 | 0.76 | 0.72 | 0.84 | 0.74 |
| (-5,1) | 0.24 | 0.84 | 0.86 | 0.89 | 0.91 | 0.94 | 0.96 |
| (-5,2) | 0.28 | 0.71 | 0.72 | 0.77 | 0.80 | 0.85 | 0.88 |
| $\mu_G = \log(8) = 2.079$ | | | | | | | |
| $(\beta_0, \sigma)$ | $pr_B(D = d)$ | Ranks based on $\widehat{OR}$ | | | Ranks based on p values | | |
|  |  | ALL | TOP 10% | TOP 20% | ALL | TOP 10% | TOP 20% |
| (-1,1) | 0.79 | 0.79 | 0.79 | 0.84 | 0.86 | 0.87 | 0.88 |
| (-1,2) | 0.76 | 0.67 | 0.66 | 0.72 | 0.74 | 0.78 | 0.68 |
| (-5,1) | 0.35 | 0.83 | 0.86 | 0.89 | 0.88 | 0.96 | 0.88 |
| (-5,2) | 0.37 | 0.71 | 0.71 | 0.77 | 0.77 | 0.89 | 0.77 |

In summary, we conclude that in the context of the genetic association studies the issue of bias due to omitting variables needs to receive more attention because it can be pronounced, in either direction and can distort false positive rate and power to detect an effect.

### ESTIMATES OF THE REDUCED VS. FULL MODELS

Suppose we obtained estimates of the genetic effects from a case-control study that omits a variable, i.e. the estimates based on the reduced model (2). The risk of the disease is, however, determined by both the genetic effects *G* and the omitted variable *O*, i.e. the full model (1).

It can be easily seen that

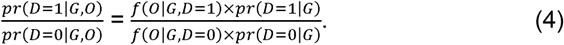

Hence

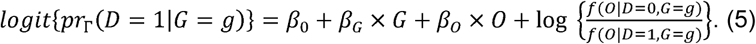

If the ratio 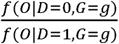 does not depend on *g*, i.e. a constant of *g*, then the estimate of *γ*_*G*_ is unbiased as the estimate of *β*_*G*_. Hence if the omitted variable and the genotype are independent conditionally on the disease status *D*, then the reduced model yields unbiased estimates of the genetic effects.

#### Bias recovery from assuming [*O*|*D*= *d, G* = *g*]

Interestingly, we derive that if [*O*|*D* = *d, G* = *g*] = *Normal*(*µ*_0_ + *µ*_*g*_ × *g* + *µ*_*d*_ × *d, σ*^2^), then it can be easily seen that

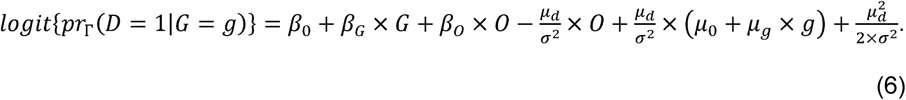

By equation (6), we can derive 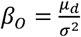 and 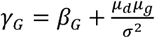. Therefore, the difference between *γ*_*G*_ and *β*_*G*_ is positive if *µ*_*d*_ × *µ*_*g*_ > 0; and the difference between *γ*_*G*_ and *β*_*G*_ is negative if *µ*_*d*_ × *µ*_*g*_ < 0. In particular, if *β*_*O*_ =0, or equivalently, *µ*_*d*_ = 0, which means that given genotype *G*, the disease *D* is conditionally independent of the omitted variable *O*, then estimate of *γ*_*G*_ is an unbiased estimate of *β*_*G*_.

#### Bias recovery from [*O*] and *pr*(*D*= 1)

In the following, we propose an approach to derive unbiased estimates of *β*_0_, *β*_*G*_ and *β*_*O*_ by solving a system of estimating equations when the auxiliary information of the omitted variable *O* is present and the rate of disease is known. Differently from the above discussions, we assume that the omitted variable *O* follows a normal distribution Normal(0,*σ*^2^) and *G* follows the Bernoulli distribution; and *O* and *G* are independent.

Based on the true model (1) and the fact that the rate of disease *pr*(*D* = 1) is known, we can obtain

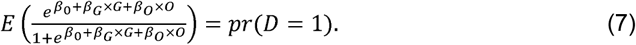

Suppose that what is available about the omitted variable *O* from the literature is the estimate from the following reduced model

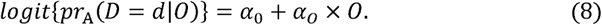

Under the logistic regression model (8), *α*_0_ and *α*_*o*_ are the solutions to the expected score equations and thus we can derive

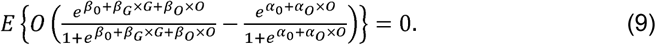

In a similar way, under the logistic regression model (2), *γ*_0_ and *γ*_*G*_ are the solutions to the expected score equations and thus we can derive

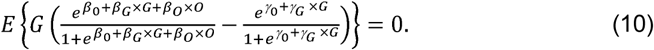

Since *α*_0_, *α*_*o*_ and *pr*(*D* = 1) are known from the literature, we can calculate *σ* by solving 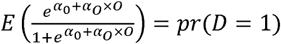. Based on the observed samples of *G* and *D*, we can derive unbiased estimates for *γ*_*G*_ and *pr*(*G* = 1) and then an unbiased estimate for *γ*_0_ can be obtained by solving 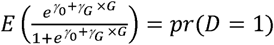. Applying numerical approximation to (7), (9) and (10), we can derive three estimating equations that only involve three unknown parameters *β*_0_, *β*_*G*_ and *β*_*O*_. Consequently, we can derive unbiased estimates for *β*_0_, *β*_*G*_ and *β*_*O*_ by solving the three estimating equations.

### SIMULATION STUDIES

The goal of the simulation studies is to assess bias in the estimates and the derivation (6) and the system of equations (7), (9) and (10). We simulate the genetic variable *G* from Bernoulli(0.1).

#### Setting 4

We are first interested to assess the equation (6). Hence simulated genotype from Bernoulli(0.1), then assumed *µ*_0_ = 0, *µ*_*g*_ = log(1.5), *µ*_*d*_ = − log(1.5), log(1), log(1.5), *σ*^2^ = 1, *β*_0_ = −1, −3.5, *β*_*G*_ = − log(2.5), − log(1.5), log(1.5), log (2.5) and 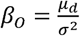. Next we generate the disease status according to model (4) for 5,000 datasets with 3,000 cases and 3,000 controls. Shown in **Table 4** and **Supplementary Table 5** are biases estimated based on (6), empirical bias, variance, MSE and power. The results suggest that the empirical bias is similar to the bias obtained through (6).

**Table 4:** Bias approximation obtained using (6), i.e.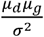, rate of the disease in the population *pr*_*B*_(*D* = *d*), bias, variance and mean squared error (MSE) of the estimates obtained from the reduced model. We simulated 5,000 datasets with 3,000 cases and 3,000 controls. We simulated genotype from Bernoulli(0.1), then assumed *µ*_0_ = 0, *µ*_g_=log(1.5),*µ*_*d*_ = − log(1.5),log (1.5),*σ*^2^ = 1, *β*_0_ = − 1,− 3.5 *β*_*G*_ *=* − log(2.5), − log(1.5),log(1.5),log(2.5).

| $\beta_0 = -3.5$ | | | | | | | |
| --- | --- | --- | --- | --- | --- | --- | --- |
| $\beta_G$ | $\mu_d$ | $\frac{\mu_d \mu_g}{\sigma^2}$ | $pr_B(D = 1)$ | Bias | Variance | MSE | power |
| $-\log(2.5)$ | $-\log(1.5)$ | -0.16 | 0.03 | -0.19 | 0.003 | 0.04 | 1 |
| $-\log(2.5)$ | $\log(1)$ | 0 | 0.03 | 0.01 | 0.003 | 0.003 | 1 |
| $-\log(2.5)$ | $\log(1.5)$ | 0.16 | 0.03 | 0.17 | 0.003 | 0.03 | 1 |
| $\log(2.5)$ | $-\log(1.5)$ | -0.16 | 0.04 | -0.16 | 0.002 | 0.03 | 1 |
| $\log(2.5)$ | $\log(1)$ | 0 | 0.03 | 0.01 | 0.002 | 0.002 | 1 |
| $\log(2.5)$ | $\log(1.5)$ | 0.16 | 0.04 | 0.17 | 0.002 | 0.03 | 1 |
| $\beta_0 = -1$ | | | | | | | |
| $\beta_G$ | $\mu_d$ | $\frac{\mu_d \mu_g}{\sigma^2}$ | $pr_B(D = 1)$ | Bias | Variance | MSE | power |
| $-\log(2.5)$ | $-\log(1.5)$ | -0.16 | 0.27 | -0.17 | 0.003 | 0.03 | 1 |
| $-\log(2.5)$ | $\log(1)$ | 0 | 0.25 | 0.01 | 0.003 | 0.003 | 1 |
| $-\log(2.5)$ | $\log(1.5)$ | 0.16 | 0.27 | 0.18 | 0.003 | 0.03 | 1 |
| $\log(2.5)$ | $-\log(1.5)$ | -0.16 | 0.30 | -0.16 | 0.002 | 0.03 | 1 |
| log(2.5) | log(1) | 0 | 0.29 | 0.002 | 0.002 | 0.002 | 1 |
| log(2.5) | log(1.5) | 0.16 | 0.31 | 0.17 | 0.002 | 0.03 | 1 |
| $\beta_0 = -3.5$ | | | | | | | |
| $\beta_G$ | $\mu_d$ | $\frac{\mu_d \mu_g}{\sigma^2}$ | $pr_B(D = 1)$ | Bias | Variance | MSE | power |
| $-\log(1.5)$ | $-\log(1.5)$ | -0.16 | 0.03 | -0.13 | 0.002 | 0.02 | 1 |
| $-\log(1.5)$ | log(1) | 0 | 0.03 | 0.02 | 0.002 | 0.003 | 1 |
| $-\log(1.5)$ | log(1.5) | 0.16 | 0.03 | 0.18 | 0.002 | 0.04 | 1 |
| log(1.5) | $-\log(1.5)$ | -0.16 | 0.03 | -0.15 | 0.002 | 0.02 | 1 |
| log(1.5) | log(1) | 0 | 0.03 | 0.01 | 0.002 | 0.002 | 1 |
| log(1.5) | log(1.5) | 0.16 | 0.03 | 0.17 | 0.002 | 0.03 | 1 |
| $\beta_0 = -1$ | | | | | | | |
| $\beta_G$ | $\mu_d$ | $\frac{\mu_d \mu_g}{\sigma^2}$ | $pr_B(D = 1)$ | Bias | Variance | MSE | power |
| $-\log(1.5)$ | $-\log(1.5)$ | -0.16 | 0.27 | -0.17 | 0.002 | 0.03 | 1 |
| $-\log(1.5)$ | log(1) | 0 | 0.26 | 0.01 | 0.002 | 0.003 | 1 |
| $-\log(1.5)$ | log(1.5) | 0.16 | 0.28 | 0.18 | 0.002 | 0.03 | 1 |
| log(1.5) | $-\log(1.5)$ | -0.16 | 0.29 | -0.17 | 0.002 | 0.03 | 1 |
| log(1.5) | log(1) | 0 | 0.28 | 0.005 | 0.002 | 0.002 | 1 |
| log(1.5) | log(1.5) | 0.16 | 0.30 | 0.17 | 0.002 | 0.031 | 1 |

#### Setting 5

We now assess the solution according to system of equations (9)-(11). We simulate the genetic variant from Bernoulli(0.1) and the omitted variable from Normal(0,1). And next we generate the disease status according to model (1) with coefficients *β*_0_ = −1, −5; *β*_*G*_ = log(2.5), log(3), log(5), log (8), *β*_*O*_ = log(5), log(8) for 5,000 datasets with 3,000 cases and 3,000 controls. Results shown in **Table 5** demonstrate that the numerical solution to the system of equations (7), (9) and (10) is nearly unbiased.

**Table 5:** Bias, Variance and Mean Squared Error (MSE) for the genetic effect estimates corrected based on the system of equations (9)-(11). We simulated 5,000 datasets of 3,000 cases and 3,000 controls. The genetic variant is simulated Bernoulli (0.10), the omitted variable is simulated from Normal(0,1) and the disease status is simulated based on model (2) with coefficients *β*_0_ = − 1, − 5; *β*_*G*_ = log(2.5), log(3), log(5), log (8), *β* _0_= log(5), log(8).

|  |  |  |  |  |
| --- | --- | --- | --- | --- |
| $\beta_G = \log(2.5)$ | | | | |
| $(\beta_0, \beta_o)$ | $pr_B(D = d)$ | Bias | Variance | MSE |
| (-1, log(5)) | 0.35 | -0.006 | 0.02 | 0.02 |
| (-5, log(5)) | 0.02 | 0.005 | 0.01 | 0.01 |
| (-1, log(8)) | 0.37 | 0.009 | 0.03 | 0.03 |
| (-5, log(8)) | 0.04 | 0.02 | 0.02 | 0.02 |
| $\beta_G = \log(3)$ | | | | |
| $(\beta_0, \beta_o)$ | $pr_B(D = d)$ | Bias | Variance | MSE |
| (-1, log(5)) | 0.35 | 0.002 | 0.02 | 0.02 |
| (-5, log(5)) | 0.02 | 0.007 | 0.01 | 0.01 |
| (-1, log(8)) | 0.37 | -0.002 | 0.03 | 0.03 |
| (-5, log(8)) | 0.04 | 0.004 | 0.02 | 0.02 |
| $\beta_G = \log(5)$ | | | | |

**Table 5:** Bias, Variance and Mean Squared Error (MSE) for the genetic effect estimates
| $(\beta_0, \beta_o)$ | $pr_B(D = d)$ | Bias | Variance | MSE |
| --- | --- | --- | --- | --- |
| $(-1, \log(5))$ | 0.36 | 0.01 | 0.02 | 0.02 |
| $(-5, \log(5))$ | 0.03 | 0.006 | 0.02 | 0.02 |
| $(-1, \log(8))$ | 0.38 | -0.02 | 0.03 | 0.03 |
| $(-5, \log(8))$ | 0.04 | -0.003 | 0.02 | 0.02 |
| $\beta_G = \log(8)$ | | | | |
| $(\beta_0, \beta_o)$ | $pr_B(D = d)$ | Bias | Variance | MSE |
| $(-1, \log(5))$ | 0.37 | 0.002 | 0.03 | 0.03 |
| $(-5, \log(5))$ | 0.03 | 0.03 | 0.02 | 0.02 |
| $(-1, \log(8))$ | 0.39 | -0.008 | 0.04 | 0.04 |
| $(-5, \log(8))$ | 0.05 | 0.02 | 0.03 | 0.03 |

### ALZHEIMER’S DISEASE STUDY

We are interested to assess what happens to the genetic effect estimates when a continuous variable is omitted from the model, i.e. how well (6) informs bias and if the system of equations (9)-(10) is capable to restore the genetic estimates. We hence consider two datasets. The Alzheimer’s Disease Neuroimaging Initiative (ADNI) dataset includes more extensive evaluations on a smaller subset of cases and controls. We hence assess how the genetic effect estimates change when continuous variables available in the dataset are omitted. Next, we consider a larger dataset generated by the Alzheimer’s Disease Genetics Consortium (ADGC) where extensive evaluations are available on a small subset. We hence assess how knowledge from the literature or from the ADNI data can be applied to inform how the genetic effect estimates change with omission of continuous variables.

#### ADNI

The set consists of 423 cases and 192 controls. We mapped the genetic variants to a set serving amyloid and tau proteins that are relevant to AD pathophysiology based on the Genecards database (https://www.genecards.org/). After preprocessing, the set contains 2,438 SNPs. The average (SD) age of cases is 74.29 (7.4), and 75.41 (4.91) in controls, p=0.058. 262(61.9%) of cases are ApoE *ε*4 carriers and 49 (25.5%) of controls are ApoE *ε*4 carriers, p<0.001. **Supplementary Table 7A-7M** further describe the sets of cases and controls and **Web-based Supplementary Materials Section B** provide extended details on the analyses of ADNI dataset.

To assess what happens to the genetic effect estimates when a continuous variable is omitted from the model, we consider several possible models and the logistic regression results, including coefficient estimates (log(OR)), standard errors (SE) and p-values are reported in **Supplementary Table 8**. We first consider a full Model 1 with age, sex, education, ApoE *ε*4 status, MMSE and a reduced model that omits MMSE (model 1A) and that omits ApoE *ε*4 status (model 1B). We next considered a full model 2 where we added ratio of hippocampus volume to whole brain volume to model 1 with the corresponding reduced model that omits the ratio of brain volumes. We observed that the difference in log(OR) estimates between the reduced and full models were on the order of ≥1*SE. For example, log(OR) for ApoE *ε*4 status changed from 1.27 (SE=0.25) to 1.62 (SE=0.20) in full model 1 vs. reduced model 1A; and from 1.005 (SE=0.268) to 1.27 (SE=0.25) in the full model 2 vs. reduced model 2A.

We next assumed that the full includes age, gender, education ApoE *ε*4 status, MMSE, and the ratio between hippocampus volume and whole brain volume, plus each of the genetic variants (Model 3). The reduced model 3A omits the ratio between brain volumes. On average, we observed that the difference in the log(OR) estimates of SNPs obtained in full model 3 vs. 3A is 0.006, with 25^th^ percentile -0.001 and 75^th^ percentile that is 0.005, minimum of -0.35 and maximum of 0.18.

We observed in the following how the SNPs rank in the full model 3 and reduced model 3A. Among the top 10 significant SNPs (ranked by p-value), 80% of the SNPs are the same in the full and reduced models, among the top 30 significant SNPs, 56.67% of the SNPs are the same and among the top 50 significant SNPs, 58% of the SNPs are the same. Hence overall, the conclusion about what SNPs should be carried to the validation set would be different based on these two models.

We also note that for all the models the distribution of p-values across all SNPs did not differ significantly from Uniform(0,1), i.e. p-values for Kolmogorov-Smirnov test are >0.05.

#### ADGC

The set consists of 2,794 cases and 667 controls (Set 1), where subsets contained data on age, sex, education, ApoE *ε*4 status (Set 2). We mapped the genetic variants to a set serving innate immune system that are relevant to AD pathophysiology (Lobach et al, 2019). After processing, the set contains 157 SNPs. The average (SD) age of cases is 70.78 (8.82), and 75.19 (8.27) in controls, p<0.001. The average (SD) education of cases is 14.08 (3.38), and 15.93 (2.72) in controls, p<0.001.1005 (48.9%) of cases are men, 109 (32.8%) of controls are men, p<0.001. 1327 (64.6%) of cases are ApoE *ε*4 carriers and 96 (28.9%) of controls are ApoE *ε*4, p<0.001. The dataset and analyses are described in extensive details in **Web-based Supplementary Materials Section C**.

We first assessed estimates in the full and reduced models based on a subset of data that includes age, sex, education, ApoE *ε*4. We observed that estimates of SNPs differed between the full (age, sex, education, ApoE *ε*4, SNP) and reduced models (omits age) by on average 0.01, 25^th^ percentile = -0.02, 75^th^ percentile = 0.04, minimum of 0.17 and maximum of 0.58.

We also note that for all the models the distribution of p-values across all SNPs did not differ significantly from Uniform(0,1), i.e. p-values for Kolmogorov-Smirnov test are >0.05.

We are next interested to asses the degree and directionality to which estimates of ApoE *ε*4 status change with the omission of age, MMSE, education, hippocampal volume and the ratio of the hippocampal volume to the whole brain volume. We therefore consider the set of 2,794 cases and 667 controls. We first estimate *γ*_ε4_ from a univariable model to be 0.16 (SE=0.01), p<0.001. We next learn the conditional distributions [*O*|*D* = *d, ε*4] of each of the omitted variables from the ADNI dataset, where we define the set of cases to be the set with diagnosis dementia and the set of controls to be the set with diagnosis of cognitively normal. Then we apply the relationship (6) to estimate the difference in the estimates due to omitting the variable as 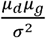. As the result, we estimate that omission of age decreases the log(OR) for ApoE *ε*4 status by 0.10, omission of MMSE increases the estimate by 0.10, omission of education increases the estimate by 0.04, omission of hippocampal brain volume increases the estimate by 0.06, and omission of the ratio between hippocampal brain volume and whole brain volume increases the estimate by 0.06.

We next assessed how the coefficient for ApoE *ε*4 status changes when MMSE is omitted from the model using system of equations (7)-(10). From the literature, we assumed that MMSE is distributed normally with mean 27 and standard deviation of 1.8; frequency of the disease in the population that is 10% and OR for MMSE that is 0.8 (95% CI: 0.55-1.1). In the reduced model the log(OR) for ApoE *ε*4 status is 1.5 (SE=0.13), p-value<0.001. Using the system of equations (7)-(10) we arrived at the log(OR) estimate that varied between 1.45 and 1.79 for various settings of the initial values that we considered.

## DISCUSSION

In the genetic association studies, we interested to accurately estimate either the parameters or the order of the magnitude of the parameters, because the estimates would determine our understanding about the underlying pathophysiologic mechanisms, risk prediction and can lead to the estimates of heritability, population attributable risk to the genetics, etc. Massive amounts of genetic data available in various databases can be utilized to estimate the genetic associations. Yet, the set of non-genetic variables is often brief.

We show that omitting a continuous variable associated with the disease status can result in substantial bias of parameter estimates in either direction. We derived two possible approaches for understanding the bias. The fist is explicit and is based on knowing [*O*|*D* = *d, G* = *g*]. The second is numerical and requires knowing the estimates from a univariable model with the omitted variable as the predictor (8) and knowing rate of the disease in the population. The two approaches that we developed differ in their assumptions. One assumes a Normal distribution for the conditional density of the omitted variable [*O*|*D* = *d, G* = *g*], i.e. assumes that the distribution of the omitted variable is a mixture of normals. The second in the system of equations assumes that the distribution of the omitted variable is normal.

Both of the approaches that we considered require knowing the set of variables in the full (true) model, what might not be feasible practically in many settings. In the analyses of Alzheimer’s disease studies we assumed various models to be the true (full) models and based on these assumptions assessed the directionality and magnitude of bias. Overall, the main contribution of our work is the justification that omitting a continuous variable from the logistic regression model can result in bias in either direction.

In some settings it is of interest to correctly estimate the order of the magnitude of the genetic effects to be able to rank the genetic markers according to strength of their association. In these settings, if the bias affects the estimates proportionally, then the bias would not change the ordering of the genetic effect estimates.

We found that if the genetic variable and the omitted variable are independent conditionally on the disease status, then omitting the variable does not result in bias of the genetic effects. This assumption is not equivalent to independence between the genotype and the omitted variable in the population.

The arguments that we’ve developed are based on the logistic link model and normality of the omitted variable. These derivations do not naturally extend to other link functions and other forms of the omitted variable.

Pirinen et al (2012) showed that for rare diseases inclusion of the key covariates can reduce power, while for common diseases inclusion of the key covariates can increase power. Our findings are similar in that the bias can either reduce or increase the magnitude of the effect. Specifically, if the omitted variable is normally distributed with [*O*|*D* = *d, G* = *g*] = *Normal* (*µ*_0_ + *µ*_*g*_ ×*g* + *µ*_*d*_ × *d, σ*^2^) then the bias is a function of *µ*_*g*_, *µ*_*d*_ and *σ*^2^. Based on this relationship we also see that a rare disease is not immune to the bias.

## Supporting information

Web-based supplement

## ACKNOWLEDGEMENTS

Dr. Lobach is supported by 5R21AG043710-02.

Data used in the preparation of this article were obtained from the Alzheimer’s Disease Neuroimaging Initiative (ADNI) database (adni.loni.usc.edu). The ADNI was launched in 2003 as a public-private partnership, led by Principal Investigator Michael W. Weiner, MD. The primary goal of ADNI has been to test whether serial magnetic resonance imaging (MRI), positron emission tomography (PET), other biological markers, and clinical and neuropsychological assessment can be combined to measure the progression of mild cognitive impairment (MCI) and early Alzheimer’s disease (AD). For up-to-date information, see www.adni-info.org.

Genotyping is performed by Alzheimer’s Disease Genetics Consortium (ADGC), U01 AG032984, RC2 AG036528. Phenotypic collection is coordinated by the National Alzheimer’s Coordinating Center (NACC), U01 AG016976

Samples from the National Cell Repository for Alzheimer’s Disease (NCRAD), which receives government support under a cooperative agreement grant (U24 AG21886) awarded by the National Institute on Aging (NIA), were used in this study. We thank contributors who collected samples used in this study, as well as patients and their families, whose help and participation made this work possible.

Data for this study were prepared, archived, and distributed by the National Institute on Aging Alzheimer’s Disease Data Storage Site (NIAGADS) at the University of Pennsylvania (U24-AG041689-01)

Data collection and sharing for this project was funded by the Alzheimer’s Disease Neuroimaging Initiative (ADNI) (National Institutes of Health Grant U01 AG024904) and DOD ADNI (Department of Defense award number W81XWH-12-2-0012). ADNI is funded by the National Institute on Aging, the National Institute of Biomedical Imaging and Bioengineering, and through generous contributions from the following: AbbVie, Alzheimer’s Association; Alzheimer’s Drug Discovery Foundation; Araclon Biotech; BioClinica, Inc.; Biogen; Bristol-Myers Squibb Company; CereSpir, Inc.; Cogstate; Eisai Inc.; Elan Pharmaceuticals, Inc.; Eli Lilly and Company; EuroImmun; F. Hoffmann-La Roche Ltd and its affiliated company Genentech, Inc.; Fujirebio; GE Healthcare; IXICO Ltd.; Janssen Alzheimer Immunotherapy Research & Development, LLC.; Johnson & Johnson Pharmaceutical Research & Development LLC.; Lumosity; Lundbeck; Merck & Co., Inc.; Meso Scale Diagnostics, LLC.; NeuroRx Research; Neurotrack Technologies; Novartis Pharmaceuticals Corporation; Pfizer Inc.; Piramal Imaging; Servier; Takeda Pharmaceutical Company; and Transition Therapeutics. The Canadian Institutes of Health Research is providing funds to support ADNI clinical sites in Canada. Private sector contributions are facilitated by the Foundation for the National Institutes of Health (www.fnih.org). The grantee organization is the Northern California Institute for Research and Education, and the study is coordinated by the Alzheimer’s Therapeutic Research Institute at the University of Southern California. ADNI data are disseminated by the Laboratory for Neuro Imaging at the University of Southern California.

