## Supplementary material for "Genetic effect estimates in case-control studies when a continuous variable is omitted from the model": Web-based supplement

**Web-based supplementary materials for**

A. **Simulation studies results**

Shown in this section are tables describing results of simulation studies.

| $\beta_{O}=\log\left( 1 \right)$ | | | | | |
| --- | --- | --- | --- | --- | --- |
| ($\beta_{0},\sigma$) | ${pr}_{B}(D=1$) | Bias | Variance | MSE | FDR |
| (-1,1) | 0.27 | 0.005 | 0.008 | 0.008 | 0.05 |
| (-1,2) | 0.27 | 0.005 | 0.008 | 0.0078 | 0.05 |
| (-5,1) | 0.007 | -0.06 | 0.006 | 0.01 | 0.08 |
| (-5,2) | 0.006 | -0.06 | 0.006 | 0.01 | 0.08 |
| $\beta_{O}=\log\left( 1.5 \right)=0.4055$ | | | | | |
| ($\beta_{0},\sigma$) | ${pr}_{B}(D=1$) | Bias | Variance | MSE | FDR |
| (-1,1) | 0.28 | -0.009 | 0.007 | 0.007 | 0.048 |
| (-1,2) | 0.29 | 0.003 | 0.008 | 0.008 | 0.05 |
| (-5,1) | 0.007 | -0.04 | 0.006 | 0.008 | 0.045 |
| (-5,2) | 0.009 | -0.05 | 0.006 | 0.009 | 0.07 |
| $\beta_{O}=\log\left( 2 \right)$=0.6931 | | | | | |
| ($\beta_{0},\sigma$) | ${pr}_{B}(D=1$) | Bias | Variance | MSE | FDR |
| (-1,1) | 0.29 | 0.007 | 0.007 | 0.007 | 0.05 |
| (-1,2) | 0.32 | 0.003 | 0.008 | 0.008 | 0.06 |
| (-5,1) | 0.009 | -0.05 | 0.006 | 0.008 | 0.06 |
| (-5,2) | 0.02 | -0.009 | 0.007 | 0.007 | 0.04 |
| $\beta_{O}=\log\left( 2.5 \right)$=0.9163 | | | | | |
| ($\beta_{0},\sigma$) | ${pr}_{B}(D=1$) | Bias | Variance | MSE | FDR |
| (-1,1) | 0.30 | 0.005 | 0.007 | 0.007 | 0.05 |
| (-1,2) | 0.35 | 0.0008 | 0.008 | 0.008 | 0.05 |
| (-5,1) | 0.01 | -0.04 | 0.007 | 0.008 | 0.05 |
| (-5,2) | 0.03 | 0.02 | 0.007 | 0.008 | 0.05 |
| $\beta_{O}=\log\left( 3 \right)$=1.0986 | | | | | |
| ($\beta_{0},\sigma$) | ${pr}_{B}(D=1$) | Bias | Variance | MSE | FDR |
| (-1,1) | 0.31 | 0.0008 | 0.007 | 0.007 | 0.048 |
| (-1,2) | 0.36 | 0.0001 | 0.007 | 0.007 | 0.047 |
| (-5,1) | 0.01 | -0.02 | 0.007 | 0.007 | 0.04 |
| (-5,2) | 0.04 | 0.02 | 0.007 | 0.007 | 0.048 |
| $\beta_{O}=\log\left( 5 \right)$=1.6094 | | | | | |
| ($\beta_{0},\sigma$) | ${pr}_{B}(D=1$) | Bias | Variance | MSE | FDR |
| (-1,1) | 0.33 | 0.002 | 0.007 | 0.007 | 0.05 |
| (-1,2) | 0.39 | -0.0009 | 0.007 | 0.007 | 0.048 |
| (-5,1) | 0.02 | 0.02 | 0.007 | 0.007 | 0.046 |
| (-5,2) | 0.09 | 0.001 | 0.007 | 0.007 | 0.047 |
| $\beta_{O}=\log\left( 8 \right)$=2.0794 | | | | | |
| ($\beta_{0},\sigma$) | ${pr}_{B}(D=1$) | Bias | Variance | MSE | FDR |
| (-1,1) | 0.36 | -0.0005 | 0.007 | 0.0072 | 0.048 |
| (-1,2) | 0.41 | -0.005 | 0.008 | 0.0075 | 0.049 |
| (-5,1) | 0.04 | 0.01 | 0.008 | 0.0073 | 0.05 |
| (-5,2) | 0.13 | -0.006 | 0.001 | 0.0071 | 0.04 |

**Supplementary Table 1**: Bias, variance and mean square error (MSE) of genetic effect estimates obtain using reduced model (2) when the data are simulated using full model (1). Shown is also probability of the disease in the population, i.e. ${pr}_{B}(D=1$), and false discovery rate (FDR). The genotype is simulated to be Bernoulli(0.1), the omitted variable is simulated from Normal(0,$\sigma^{2}$). We simulated the disease status from model (1) with parameters $\beta_{0}=-1,-5; \beta_{G}=\log\left( 1 \right)=0, \beta_{O}=\log\left( 1 \right),\log\left( 1.5 \right),\log\left( 2 \right),\log\left( 2.5 \right),\log\left( 3 \right),\log\left( 5 \right),\log\left( 8 \right).$ The results are based on 5,000 datasets of 3,000 cases and 3,000 controls.

| $\beta_{G}=\log\left( 2.5 \right)$=0.9163, $\beta_{O}=\log\left( 1 \right)$ | | | | | |
| --- | --- | --- | --- | --- | --- |
| ($\beta_{0},\sigma$) | ${pr}_{B}(D=1$) | Bias | Variance | MSE | power |
| (-1,1) | 0.29 | 0.006 | 0.007 | 0.007 | 1 |
| (-1,2) | 0.29 | 0.006 | 0.007 | 0.007 | 1 |
| (-5,1) | 0.008 | -0.01 | 0.005 | 0.005 | 1 |
| (-5,2) | 0.008 | -0.01 | 0.005 | 0.005 | 1 |
| $\beta_{G}=\log\left( 2.5 \right)$=0.9163, $\beta_{O}=\log\left( 1.5 \right)$ =0.4055 | | | | | |
| ($\beta_{0},\sigma$) | ${pr}_{B}(D=1$) | Bias | Variance | MSE | power |
| (-1,1) | 0.30 | -0.04 | 0.007 | 0.009 | 1 |
| (-1,2) | 0.31 | -0.12 | 0.007 | 0.02 | 1 |
| (-5,1) | 0.008 | -0.02 | 0.005 | 0.006 | 1 |
| (-5,2) | 0.01 | -0.03 | 0.005 | 0.006 | 1 |
| $\beta_{G}=\log\left( 2.5 \right)$=0.9163, $\beta_{O}=\log\left( 2 \right)$=0.6931 | | | | | |
| ($\beta_{0},\sigma$) | ${pr}_{B}(D=1$) | Bias | Variance | RMSE | power |
| (-1,1) | 0.31 | -0.09 | 0.007 | 0.02 | 1 |
| (-1,2) | 0.34 | -0.24 | 0.007 | 0.064 | 1 |
| (-5,1) | 0.01 | -0.02 | 0.005 | 0.006 | 1 |
| (-5,2) | 0.02 | -0.06 | 0.006 | 0.009 | 1 |
| $\beta_{G}=\log\left( 2.5 \right)$=0.9163, $\beta_{O}=\log\left( 2.5 \right)$=0.9163 | | | | | |
| ($\beta_{0},\sigma$) | ${pr}_{B}(D=1$) | Bias | Variance | RMSE | power |
| (-1,1) | 0.32 | -0.14 | 0.007 | 0.03 | 1 |
| (-1,2) | 0.36 | -0.33 | 0.007 | 0.12 | 1 |
| (-5,1) | 0.01 | -0.02 | 0.005 | 0.006 | 1 |
| (-5,2) | 0.03 | -0.18 | 0.006 | 0.04 | 1 |
| $\beta_{G}=\log\left( 2.5 \right)$=0.9163, $\beta_{O}=\log\left( 3 \right)$=1.0986 | | | | | |
| ($\beta_{0},\sigma$) | ${pr}_{B}(D=1$) | Bias | Variance | RMSE | power |
| (-1,1) | 0.33 | -0.18 | 0.007 | 0.04 | 1 |
| (-1,2) | 0.37 | -0.39 | 0.007 | 0.16 | 1 |
| (-5,1) | 0.01 | -0.03 | 0.005 | 0.006 | 1 |
| (-5,2) | 0.04 | -0.26 | 0.006 | 0.07 | 1 |
| $\beta_{G}=\log\left( 2.5 \right)$=0.9163, $\beta_{O}=\log\left( 5 \right)=1.6094$ | | | | | |
| ($\beta_{0},\sigma$) | ${pr}_{B}(D=1$) | Bias | Variance | RMSE | power |
| (-1,1) | 0.35 | -0.29 | 0.007 | 0.09 | 1 |
| (-1,2) | 0.40 | -0.52 | 0.008 | 0.28 | 1 |
| (-5,1) | 0.02 | -0.12 | 0.006 | 0.02 | 1 |
| (-5,2) | 0.09 | -0.44 | 0.007 | 0.2 | 1 |
| $\beta_{G}=\log\left( 2.5 \right)$=0.9163, $\beta_{O}=\log\left( 8 \right)=2.0794$ | | | | | |
| ($\beta_{0},\sigma$) | ${pr}_{B}(D=1$) | Bias | Variance | RMSE | power |
| (-1,1) | 0.37 | -0.38 | 0.007 | 0.15 | 1 |
| (-1,2) | 0.42 | -0.60 | 0.007 | 0.36 | 0.97 |
| (-5,1) | 0.04 | -0.23 | 0.006 | 0.06 | 1 |
| (-5,2) | 0.14 | -0.56 | 0.007 | 0.32 | 0.99 |
| $\beta_{G}=\log\left( 3 \right)$=1.0986, $\beta_{O}=\log\left( 1 \right)$ | | | | | |
| ($\beta_{0},\sigma$) | ${pr}_{B}(D=1$) | Bias | Variance | RMSE | power |
| (-1,1) | 0.29 | -0.01 | 0.008 | 0.008 | 1 |
| (-1,2) | 0.29 | -0.01 | 0.008 | 0.008 | 1 |
| (-5,1) | 0.008 | -0.01 | 0.005 | 0.005 | 1 |
| (-5,2) | 0.008 | -0.01 | 0.005 | 0.005 | 1 |
| $\beta_{G}=\log\left( 3 \right)$=1.0986, $\beta_{O}=\log\left( 1.5 \right)$ =0.4055 | | | | | |
| ($\beta_{0},\sigma$) | ${pr}_{B}(D=1$) | Bias | Variance | RMSE | power |
| (-1,1) | 0.30 | -0.05 | 0.007 | 0.01 | 1 |
| (-1,2) | 0.32 | -0.14 | 0.008 | 0.03 | 1 |
| (-5,1) | 0.009 | -0.01 | 0.005 | 0.005 | 1 |
| (-5,2) | 0.01 | -0.03 | 0.005 | 0.006 | 1 |
| $\beta_{G}=\log\left( 3 \right)$=1.0986, $\beta_{O}=\log\left( 2 \right)$=0.6931 | | | | | |
| ($\beta_{0},\sigma$) | ${pr}_{B}(D=1$) | Bias | Variance | RMSE | power |
| (-1,1) | 0.31 | -0.11 | 0.008 | 0.02 | 1 |
| (-1,2) | 0.34 | -0.28 | 0.008 | 0.09 | 1 |
| (-5,1) | 0.01 | -0.03 | 0.005 | 0.006 | 1 |
| (-5,2) | 0.02 | -0.09 | 0.005 | 0.01 | 1 |
| $\beta_{G}=\log\left( 3 \right)$=1.0986, $\beta_{O}=\log\left( 2.5 \right)$=0.9163 | | | | | |
| ($\beta_{0},\sigma$) | ${pr}_{B}(D=1$) | Bias | Variance | RMSE | power |
| (-1,1) | 0.32 | -0.16 | 0.007 | 0.03 | 1 |
| (-1,2) | 0.36 | -0.39 | 0.007 | 0.16 | 1 |
| (-5,1) | 0.01 | -0.03 | 0.005 | 0.007 | 1 |
| (-5,2) | 0.030 | -0.21 | 0.005 | 0.05 | 1 |
| $\beta_{G}=\log\left( 3 \right)$=1.0986, $\beta_{O}=\log\left( 3 \right)$=1.0986 | | | | | |
| ($\beta_{0},\sigma$) | ${pr}_{B}(D=1$) | Bias | Variance | RMSE | power |
| (-1,1) | 0.33 | -0.21 | 0.007 | 0.05 | 1 |
| (-1,2) | 0.38 | -0.47 | 0.007 | 0.23 | 1 |
| (-5,1) | 0.01 | -0.04 | 0.005 | 0.007 | 1 |
| (-5,2) | 0.04 | -0.31 | 0.006 | 0.10 | 1 |
| $\beta_{G}=\log\left( 3 \right)$=1.0986, $\beta_{O}=\log\left( 5 \right)=1.6094$ | | | | | |
| ($\beta_{0},\sigma$) | ${pr}_{B}(D=1$) | Bias | Variance | RMSE | power |
| (-1,1) | 0.35 | -0.34 | 0.008 | 0.13 | 1 |
| (-1,2) | 0.40 | -0.62 | 0.007 | 0.39 | 1 |
| (-5,1) | 0.02 | -0.15 | 0.005 | 0.03 | 1 |
| (-5,2) | 0.09 | -0.54 | 0.006 | 0.29 | 1 |
| $\beta_{G}=\log\left( 3 \right)$=1.0986, $\beta_{O}=\log\left( 8 \right)=2.0794$ | | | | | |
| ($\beta_{0},\sigma$) | ${pr}_{B}(D=1$) | Bias | Variance | RMSE | power |
| (-1,1) | 0.37 | -0.44 | 0.008 | 0.20 | 1 |
| (-1,2) | 0.42 | -0.71 | 0.007 | 0.51 | 1 |
| (-5,1) | 0.04 | -0.28 | 0.006 | 0.08 | 1 |
| (-5,2) | 0.14 | -0.67 | 0.007 | 0.46 | 1 |
| $\beta_{G}=\log\left( 5 \right)=1.6094$, $\beta_{O}=\log\left( 1 \right)$ | | | | | |
| ($\beta_{0},\sigma$) | ${pr}_{B}(D=1$) | Bias | Variance | RMSE | power |
| (-1,1) | 0.31 | -0.01 | 0.009 | 0.009 | 1 |
| (-1,2) | 0.31 | -0.01 | 0.009 | 0.009 | 1 |
| (-5,1) | 0.009 | -0.01 | 0.005 | 0.005 | 1 |
| (-5,2) | 0.009 | -0.01 | 0.005 | 0.005 | 1 |
| $\beta_{G}=\log\left( 5 \right)=1.6094$, $\beta_{O}=\log\left( 1.5 \right)$ =0.4055 | | | | | |
| ($\beta_{0},\sigma$) | ${pr}_{B}(D=1$) | Bias | Variance | RMSE | power |
| (-1,1) | 0.313 | -0.07 | 0.008 | 0.01 | 1 |
| (-1,2) | 0.33 | -0.20 | 0.008 | 0.05 | 1 |
| (-5,1) | 0.01 | -0.02 | 0.005 | 0.005 | 1 |
| (-5,2) | 0.01 | -0.03 | 0.005 | 0.006 | 1 |
| $\beta_{G}=\log\left( 5 \right)=1.6094$, $\beta_{O}=\log\left( 2 \right)$=0.6931 | | | | | |
| ($\beta_{0},\sigma$) | ${pr}_{B}(D=1$) | Bias | Variance | RMSE | power |
| (-1,1) | 0.32 | -0.16 | 0.009 | 0.03 | 1 |
| (-1,2) | 0.35 | -0.43 | 0.008 | 0.19 | 1 |
| (-5,1) | 0.01 | -0.03 | 0.005 | 0.006 | 1 |
| (-5,2) | 0.02 | -0.17 | 0.005 | 0.03 | 1 |
| $\beta_{G}=\log\left( 5 \right)=1.6094$, $\beta_{O}=\log\left( 2.5 \right)$=0.9163 | | | | | |
| ($\beta_{0},\sigma$) | ${pr}_{B}(D=1$) | Bias | Variance | RMSE | power |
| (-1,1) | 0.33 | -0.24 | 0.008 | 0.06 | 1 |
| (-1,2) | 0.37 | -0.574 | 0.008 | 0.34 | 1 |
| (-5,1) | 0.014 | -0.05 | 0.005 | 0.007 | 1 |
| (-5,2) | 0.03 | -0.32 | 0.005 | 0.11 | 1 |
| $\beta_{G}=\log\left( 5 \right)=1.6094$, $\beta_{O}=\log\left( 3 \right)$=1.0986 | | | | | |
| ($\beta_{0},\sigma$) | ${pr}_{B}(D=1$) | Bias | Variance | RMSE | power |
| (-1,1) | 0.34 | -0.31 | 0.008 | 0.11 | 1 |
| (-1,2) | 0.38 | -0.68 | 0.008 | 0.47 | 1 |
| (-5,1) | 0.02 | -0.08 | 0.005 | 0.01 | 1 |
| (-5,2) | 0.05 | -0.48 | 0.005 | 0.23 | 1 |
| $\beta_{G}=\log\left( 5 \right)=1.6094$, $\beta_{O}=\log\left( 5 \right)=1.6094$ | | | | | |
| ($\beta_{0},\sigma$) | ${pr}_{B}(D=1$) | Bias | Variance | RMSE | power |
| (-1,1) | 0.36 | -0.50 | 0.008 | 0.26 | 1 |
| (-1,2) | 0.41 | -0.91 | 0.008 | 0.83 | 1 |
| (-5,1) | 0.03 | -0.23 | 0.005 | 0.06 | 1 |
| (-5,2) | 0.10 | -0.79 | 0.006 | 0.63 | 1 |
| $\beta_{G}=\log\left( 5 \right)=1.6094$, $\beta_{O}=\log\left( 8 \right)=2.0794$ | | | | | |
| ($\beta_{0},\sigma$) | ${pr}_{B}(D=1$) | Bias | Variance | RMSE | power |
| (-1,1) | 0.38 | -0.65 | 0.008 | 0.43 | 1 |
| (-1,2) | 0.43 | -1.04 | 0.008 | 1.08 | 1 |
| (-5,1) | 0.04 | -0.43 | 0.006 | 0.19 | 1 |
| (-5,2) | 0.14 | -0.98 | 0.007 | 0.97 | 1 |

**Supplementary Table 2**: Bias, variance and mean square error (MSE) of genetic effect estimates obtain using reduced model (2) when the data are simulated using full model (1). Shown is also probability of the disease in the population, i.e. ${pr}_{B}(D=1$), and false discovery rate (FDR). The genotype is simulated to be Bernoulli(0.1), the omitted variable is simulated from Normal(0,$\sigma^{2}$). We simulated the disease status from model (1) with parameters $\beta_{0}=-1,-5; \beta_{G}=\log\left( 2 \right),\log\left( 3 \right),log(5), \beta_{O}=\log\left( 1 \right),\log\left( 1.5 \right),\log\left( 2 \right),\log\left( 2.5 \right),\log\left( 3 \right),\log\left( 5 \right),\log\left( 8 \right).$ The results are based on 5,000 datasets of 3,000 cases and 3,000 controls.

| $\beta_{G}=\log\left( 1 \right)$ , $\beta_{O}=\log\left( 1 \right)$ | | | | | |
| --- | --- | --- | --- | --- | --- |
| ($\beta_{0},\sigma$) | ${pr}_{B}(D=1$) | Bias | Variance | RMSE | FDR |
| (-1,1) | 0.2691 | -0.0064 | 0.0023 | 0.0023 | 0.0562 |
| (-1,2) | 0.269 | 0.0031 | 0.0022 | 0.0022 | 0.0532 |
| (-5,1) | 0.0066 | 0.0051 | 0.002 | 0.002 | 0.0404 |
| (-5,2) | 0.0066 | 0.0051 | 0.002 | 0.002 | 0.0404 |
| $\beta_{G}=\log\left( 1 \right)$ , $\beta_{O}=\log\left( 1.5 \right)=0.4055$ | | | | | |
| ($\beta_{0},\sigma$) | ${pr}_{B}(D=1$) | Bias | Variance | RMSE | FDR |
| (-1,1) | 0.2761 | 0.0012 | 0.0022 | 0.0022 | 0.0504 |
| (-1,2) | 0.2935 | -6e-04 | 0.0022 | 0.0022 | 0.0496 |
| (-5,1) | 0.0072 | 0.0121 | 0.0021 | 0.0023 | 0.0538 |
| (-5,2) | 0.0092 | 0.0215 | 0.0021 | 0.0026 | 0.0716 |
| $\beta_{G}=\log\left( 1 \right)$ , $\beta_{O}=\log\left( 2 \right)$=0.6931 | | | | | |
| ($\beta_{0},\sigma$) | ${pr}_{B}(D=1$) | Bias | Variance | RMSE | FDR |
| (-1,1) | 0.2879 | -0.0020 | 0.0022 | 0.0022 | 0.0458 |
| (-1,2) | 0.3235 | 0.0007 | 0.0022 | 0.0022 | 0.0464 |
| (-5,1) | 0.0084 | 0.0189 | 0.0020 | 0.0024 | 0.0552 |
| (-5,2) | 0.0161 | 0.0089 | 0.0021 | 0.0022 | 0.0494 |
| $\beta_{G}=\log\left( 1 \right)$ , $\beta_{O}=\log\left( 2.5 \right)$=0.9163 | | | | | |
| ($\beta_{0},\sigma$) | ${pr}_{B}(D=1$) | Bias | Variance | RMSE | FDR |
| (-1,1) | 0.2991 | -0.0013 | 0.0022 | 0.0022 | 0.0468 |
| (-1,2) | 0.3449 | -0.0002 | 0.0022 | 0.0022 | 0.0440 |
| (-5,1) | 0.0100 | 0.0189 | 0.0020 | 0.0024 | 0.0572 |
| (-5,2) | 0.0268 | 0.0032 | 0.0021 | 0.0021 | 0.0436 |
| $\beta_{G}=\log\left( 1 \right)$ , $\beta_{O}=\log\left( 3 \right)$=1.0986 | | | | | |
| ($\beta_{0},\sigma$) | ${pr}_{B}(D=1$) | Bias | Variance | RMSE | FDR |
| (-1,1) | 0.3085 | 0.0014 | 0.0022 | 0.0022 | 0.0470 |
| (-1,2) | 0.3600 | 0.0006 | 0.0022 | 0.0022 | 0.0440 |
| (-5,1) | 0.0118 | 0.0140 | 0.0021 | 0.0023 | 0.0510 |
| (-5,2) | 0.0395 | 0.0015 | 0.0021 | 0.0021 | 0.0414 |
| $\beta_{G}=\log\left( 1 \right)$ , $\beta_{O}=\log\left( 5 \right)$=1.6094 | | | | | |
| ($\beta_{0},\sigma$) | ${pr}_{B}(D=1$) | Bias | Variance | RMSE | FDR |
| (-1,1) | 0.3345 | 0.0010 | 0.0022 | 0.0022 | 0.0460 |
| (-1,2) | 0.3925 | 0.0009 | 0.0022 | 0.0022 | 0.0452 |
| (-5,1) | 0.0208 | 0.0061 | 0.0021 | 0.0021 | 0.0438 |
| (-5,2) | 0.0872 | 0.0017 | 0.0021 | 0.0021 | 0.0446 |
| $\beta_{G}=\log\left( 1 \right)$ , $\beta_{O}=\log\left( 8 \right)$=2.0794 | | | | | |
| ($\beta_{0},\sigma$) | ${pr}_{B}(D=1$) | Bias | Variance | RMSE | FDR |
| (-1,1) | 0.3555 | 0.0017 | 0.0022 | 0.0022 | 0.0448 |
| (-1,2) | 0.4125 | -0.0008 | 0.0022 | 0.0022 | 0.0464 |
| (-5,1) | 0.0350 | 0.0004 | 0.0021 | 0.0021 | 0.0464 |
| (-5,2) | 0.1346 | -0.0013 | 0.0021 | 0.0021 | 0.0460 |
| $\beta_{G}=\log\left( 1.5 \right)=0.4055$ , $\beta_{O}=\log\left( 1 \right)$ | | | | | |
| ($\beta_{0},\sigma$) | ${pr}_{B}(D=1$) | Bias | Variance | RMSE | power |
| (-1,1) | 0.2777 | -0.0026 | 0.0021 | 0.0021 | 1 |
| (-1,2) | 0.2777 | 0.0014 | 0.0021 | 0.0021 | 1 |
| (-5,1) | 0.0070 | 0.0160 | 0.0018 | 0.0021 | 1 |
| (-5,2) | 0.0070 | 0.0160 | 0.0018 | 0.0021 | 1 |
| $\beta_{G}=\log\left( 1.5 \right)=0.4055$ , $\beta_{O}=\log\left( 1.5 \right)=0.4055$ | | | | | |
| ($\beta_{0},\sigma$) | ${pr}_{B}(D=1$) | Bias | Variance | RMSE | power |
| (-1,1) | 0.2845 | -0.0113 | 0.0021 | 0.0023 | 1 |
| (-1,2) | 0.3014 | -0.0454 | 0.0021 | 0.0042 | 1 |
| (-5,1) | 0.0076 | 0.0164 | 0.0018 | 0.0021 | 1 |
| (-5,2) | 0.0096 | 0.0123 | 0.0019 | 0.002 | 1 |
| $\beta_{G}=\log\left( 1.5 \right)=0.4055$ , $\beta_{O}=\log\left( 2 \right)$=0.6931 | | | | | |
| ($\beta_{0},\sigma$) | ${pr}_{B}(D=1$) | Bias | Variance | RMSE | power |
| (-1,1) | 0.2960 | -0.0398 | 0.0021 | 0.0037 | 1 |
| (-1,2) | 0.3304 | -0.1055 | 0.0021 | 0.0133 | 1 |
| (-5,1) | 0.0088 | 0.0162 | 0.0018 | 0.0020 | 1 |
| (-5,2) | 0.0168 | -0.0139 | 0.0019 | 0.0021 | 1 |
| $\beta_{G}=\log\left( 1.5 \right)=0.4055$ , $\beta_{O}=\log\left( 2.5 \right)$=0.9163 | | | | | |
| ($\beta_{0},\sigma$) | ${pr}_{B}(D=1$) | Bias | Variance | RMSE | power |
| (-1,1) | 0.3068 | -0.0609 | 0.0021 | 0.0058 | 1 |
| (-1,2) | 0.3510 | -0.1453 | 0.0021 | 0.0233 | 1 |
| (-5,1) | 0.0105 | 0.0091 | 0.0018 | 0.0019 | 1 |
| (-5,2) | 0.0279 | -0.0604 | 0.0019 | 0.0056 | 1 |
| $\beta_{G}=\log\left( 1.5 \right)=0.4055$ , $\beta_{O}=\log\left( 3 \right)$=1.0986 | | | | | |
| ($\beta_{0},\sigma$) | ${pr}_{B}(D=1$) | Bias | Variance | RMSE | power |
| (-1,1) | 0.3159 | -0.0767 | 0.0021 | 0.0080 | 1 |
| (-1,2) | 0.3656 | -0.1698 | 0.0022 | 0.0310 | 0.9996 |
| (-5,1) | 0.0124 | 0.0019 | 0.0019 | 0.0019 | 1 |
| (-5,2) | 0.0408 | -0.0954 | 0.0019 | 0.0110 | 1 |
| $\beta_{G}=\log\left( 1.5 \right)=0.4055$ , $\beta_{O}=\log\left( 5 \right)$=1.6094 | | | | | |
| ($\beta_{0},\sigma$) | ${pr}_{B}(D=1$) | Bias | Variance | RMSE | power |
| (-1,1) | 0.3410 | -0.1259 | 0.0021 | 0.0180 | 1 |
| (-1,2) | 0.3968 | -0.2279 | 0.0022 | 0.0541 | 0.9634 |
| (-5,1) | 0.0217 | -0.0341 | 0.0019 | 0.0031 | 1 |
| (-5,2) | 0.0891 | -0.1910 | 0.0020 | 0.0385 | 0.9982 |
| $\beta_{G}=\log\left( 1.5 \right)=0.4055$ , $\beta_{O}=\log\left( 8 \right)$=2.0794 | | | | | |
| ($\beta_{0},\sigma$) | ${pr}_{B}(D=1$) | Bias | Variance | RMSE | power |
| (-1,1) | 0.3611 | -0.1624 | 0.0021 | 0.0285 | 1 |
| (-1,2) | 0.4160 | -0.2624 | 0.0022 | 0.0710 | 0.8652 |
| (-5,1) | 0.0363 | -0.0846 | 0.0020 | 0.0091 | 1 |
| (-5,2) | 0.1366 | -0.2405 | 0.0021 | 0.0599 | 0.9440 |
| $\beta_{G}=\log\left( 2 \right)$=0.6931 , $\beta_{O}=\log\left( 1 \right)$ | | | | | |
| ($\beta_{0},\sigma$) | ${pr}_{B}(D=1$) | Bias | Variance | RMSE | power |
| (-1,1) | 0.2845 | 0.0024 | 0.0022 | 0.0022 | 1 |
| (-1,2) | 0.2845 | 0.0024 | 0.0022 | 0.0022 | 1 |
| (-5,1) | 0.0073 | 0.0159 | 0.0017 | 0.0020 | 1 |
| (-5,2) | 0.0073 | 0.0159 | 0.0017 | 0.0020 | 1 |
| $\beta_{G}=\log\left( 2 \right)$=0.6931 , $\beta_{O}=\log\left( 1.5 \right)=0.4055$ | | | | | |
| ($\beta_{0},\sigma$) | ${pr}_{B}(D=1$) | Bias | Variance | RMSE | power |
| (-1,1) | 0.291 | -0.0227 | 0.0022 | 0.0027 | 1 |
| (-1,2) | 0.3075 | -0.0824 | 0.0021 | 0.0089 | 1 |
| (-5,1) | 0.0079 | 0.0139 | 0.0017 | 0.0019 | 1 |
| (-5,2) | 0.01 | 0.0045 | 0.0017 | 0.0018 | 1 |
| $\beta_{G}=\log\left( 2 \right)$=0.6931 , $\beta_{O}=\log\left( 2 \right)$=0.6931 | | | | | |
| ($\beta_{0},\sigma$) | ${pr}_{B}(D=1$) | Bias | Variance | RMSE | power |
| (-1,1) | 0.3021 | -0.0641 | 0.0021 | 0.0063 | 1 |
| (-1,2) | 0.3355 | -0.1806 | 0.0022 | 0.0348 | 1 |
| (-5,1) | 0.0092 | 0.0070 | 0.0017 | 0.0017 | 1 |
| (-5,2) | 0.0175 | -0.0391 | 0.0018 | 0.0033 | 1 |
| $\beta_{G}=\log\left( 2 \right)$=0.6931 , $\beta_{O}=\log\left( 2.5 \right)$=0.9163 | | | | | |
| ($\beta_{0},\sigma$) | ${pr}_{B}(D=1$) | Bias | Variance | RMSE | power |
| (-1,1) | 0.3125 | -0.1004 | 0.0022 | 0.0122 | 1 |
| (-1,2) | 0.3555 | -0.2480 | 0.0022 | 0.0637 | 1 |
| (-5,1) | 0.0109 | -0.0001 | 0.0017 | 0.0017 | 1 |
| (-5,2) | 0.0288 | -0.1111 | 0.0018 | 0.0142 | 1 |
| $\beta_{G}=\log\left( 2 \right)$=0.6931 , $\beta_{O}=\log\left( 3 \right)$=1.0986 | | | | | |
| ($\beta_{0},\sigma$) | ${pr}_{B}(D=1$) | Bias | Variance | RMSE | power |
| (-1,1) | 0.3215 | -0.1309 | 0.0022 | 0.0193 | 1 |
| (-1,2) | 0.3696 | -0.2941 | 0.0022 | 0.0887 | 1 |
| (-5,1) | 0.0129 | -0.0096 | 0.0018 | 0.0019 | 1 |
| (-5,2) | 0.0420 | -0.1766 | 0.0019 | 0.0331 | 1 |
| $\beta_{G}=\log\left( 2 \right)$=0.6931 , $\beta_{O}=\log\left( 5 \right)$=1.6094 | | | | | |
| ($\beta_{0},\sigma$) | ${pr}_{B}(D=1$) | Bias | Variance | RMSE | power |
| (-1,1) | 0.3458 | -0.2151 | 0.0022 | 0.0484 | 1 |
| (-1,2) | 0.3999 | -0.3897 | 0.0022 | 0.1540 | 1 |
| (-5,1) | 0.0225 | -0.0737 | 0.0018 | 0.0072 | 1 |
| (-5,2) | 0.0906 | -0.3297 | 0.0020 | 0.1107 | 1 |
| $\beta_{G}=\log\left( 2 \right)$=0.6931 , $\beta_{O}=\log\left( 8 \right)$=2.0794 | | | | | |
| ($\beta_{0},\sigma$) | ${pr}_{B}(D=1$) | Bias | Variance | RMSE | power |
| (-1,1) | 0.3653 | -0.2802 | 0.0022 | 0.0807 | 1 |
| (-1,2) | 0.4186 | -0.4467 | 0.0022 | 0.2017 | 0.9998 |
| (-5,1) | 0.0374 | -0.1562 | 0.0019 | 0.0263 | 1 |
| (-5,2) | 0.1381 | -0.4173 | 0.0020 | 0.1762 | 1 |
| $\beta_{G}=\log\left( 2.5 \right)$=0.9163 , $\beta_{O}=\log\left( 1 \right)$ | | | | | |
| ($\beta_{0},\sigma$) | ${pr}_{B}(D=1$) | Bias | Variance | RMSE | power |
| (-1,1) | 0.2900 | 0.0024 | 0.0022 | 0.0022 | 1 |
| (-1,2) | 0.2900 | 0.0024 | 0.0022 | 0.0022 | 1 |
| (-5,1) | 0.0076 | 0.0156 | 0.0016 | 0.0019 | 1 |
| (-5,2) | 0.0076 | 0.0156 | 0.0016 | 0.0019 | 1 |
| $\beta_{G}=\log\left( 2.5 \right)$=0.9163, $\beta_{O}=\log\left( 1.5 \right)=0.4055$ | | | | | |
| ($\beta_{0},\sigma$) | ${pr}_{B}(D=1$) | Bias | Variance | RMSE | power |
| (-1,1) | 0.2964 | -0.0325 | 0.0022 | 0.0032 | 1 |
| (-1,2) | 0.3123 | -0.1109 | 0.0022 | 0.0145 | 1 |
| (-5,1) | 0.0083 | 0.0189 | 0.0016 | 0.002 | 1 |
| (-5,2) | 0.0105 | 0.0024 | 0.0016 | 0.0016 | 1 |
| $\beta_{G}=\log\left( 2.5 \right)$=0.9163, $\beta_{O}=\log\left( 2 \right)$=0.6931 | | | | | |
| ($\beta_{0},\sigma$) | ${pr}_{B}(D=1$) | Bias | Variance | RMSE | power |
| (-1,1) | 0.3070 | -0.0854 | 0.0022 | 0.0095 | 1 |
| (-1,2) | 0.3396 | -0.2394 | 0.0022 | 0.0595 | 1 |
| (-5,1) | 0.0096 | 0.0066 | 0.0016 | 0.0017 | 1 |
| (-5,2) | 0.0181 | -0.0587 | 0.0017 | 0.0051 | 1 |
| $\beta_{G}=\log\left( 2.5 \right)$=0.9163, $\beta_{O}=\log\left( 2.5 \right)$=0.9163 | | | | | |
| ($\beta_{0},\sigma$) | ${pr}_{B}(D=1$) | Bias | Variance | RMSE | power |
| (-1,1) | 0.3172 | -0.1347 | 0.0022 | 0.0204 | 1 |
| (-1,2) | 0.3591 | -0.3278 | 0.0022 | 0.1097 | 1 |
| (-5,1) | 0.0114 | -0.0061 | 0.0016 | 0.0017 | 1 |
| (-5,2) | 0.0297 | -0.1578 | 0.0018 | 0.0267 | 1 |
| $\beta_{G}=\log\left( 2.5 \right)$=0.9163, $\beta_{O}=\log\left( 3 \right)$=1.0986 | | | | | |
| ($\beta_{0},\sigma$) | ${pr}_{B}(D=1$) | Bias | Variance | RMSE | power |
| (-1,1) | 0.3260 | -0.1753 | 0.0022 | 0.0330 | 1 |
| (-1,2) | 0.3728 | -0.3890 | 0.0022 | 0.1535 | 1 |
| (-5,1) | 0.0134 | -0.0176 | 0.0017 | 0.0020 | 1 |
| (-5,2) | 0.0430 | -0.2440 | 0.0018 | 0.0613 | 1 |
| $\beta_{G}=\log\left( 2.5 \right)$=0.9163, $\beta_{O}=\log\left( 5 \right)$=1.6094 | | | | | |
| ($\beta_{0},\sigma$) | ${pr}_{B}(D=1$) | Bias | Variance | RMSE | power |
| (-1,1) | 0.3496 | -0.2850 | 0.0022 | 0.0834 | 1 |
| (-1,2) | 0.4023 | -0.5154 | 0.0022 | 0.2678 | 1 |
| (-5,1) | 0.0232 | -0.1063 | 0.0017 | 0.0130 | 1 |
| (-5,2) | 0.0918 | -0.4421 | 0.0019 | 0.1973 | 1 |
| $\beta_{G}=\log\left( 2.5 \right)$=0.9163, $\beta_{O}=\log\left( 8 \right)$=2.0794 | | | | | |
| ($\beta_{0},\sigma$) | ${pr}_{B}(D=1$) | Bias | Variance | RMSE | power |
| (-1,1) | 0.3686 | -0.3704 | 0.0022 | 0.1394 | 1 |
| (-1,2) | 0.4205 | -0.5913 | 0.0022 | 0.3519 | 1 |
| (-5,1) | 0.0384 | -0.2144 | 0.0018 | 0.0478 | 1 |
| (-5,2) | 0.1394 | -0.5519 | 0.0020 | 0.3066 | 1 |
| $\beta_{G}=\log\left( 3 \right)$=1.0986 , $\beta_{O}=\log\left( 1 \right)$ | | | | | |
| ($\beta_{0},\sigma$) | ${pr}_{B}(D=1$) | Bias | Variance | RMSE | power |
| (-1,1) | 0.2945 | -0.0010 | 0.0023 | 0.0023 | 1 |
| (-1,2) | 0.2945 | -0.0010 | 0.0023 | 0.0023 | 1 |
| (-5,1) | 0.0080 | 0.0149 | 0.0016 | 0.0018 | 1 |
| (-5,2) | 0.0080 | 0.0149 | 0.0016 | 0.0018 | 1 |
| $\beta_{G}=\log\left( 3 \right)$=1.0986, $\beta_{O}=\log\left( 1.5 \right)=0.4055$ | | | | | |
| ($\beta_{0},\sigma$) | ${pr}_{B}(D=1$) | Bias | Variance | RMSE | power |
| (-1,1) | 0.3007 | -0.0405 | 0.0022 | 0.0039 | 1 |
| (-1,2) | 0.3163 | -0.1331 | 0.0023 | 0.02 | 1 |
| (-5,1) | 0.0086 | 0.0156 | 0.0016 | 0.0018 | 1 |
| (-5,2) | 0.0109 | -0.0011 | 0.0017 | 0.0017 | 1 |
| $\beta_{G}=\log\left( 3 \right)$=1.0986, $\beta_{O}=\log\left( 2 \right)$=0.6931 | | | | | |
| ($\beta_{0},\sigma$) | ${pr}_{B}(D=1$) | Bias | Variance | RMSE | power |
| (-1,1) | 0.3112 | -0.1031 | 0.0023 | 0.0129 | 1 |
| (-1,2) | 0.3430 | -0.2876 | 0.0023 | 0.0850 | 1 |
| (-5,1) | 0.0100 | 0.0033 | 0.0016 | 0.0016 | 1 |
| (-5,2) | 0.0188 | -0.0777 | 0.0017 | 0.0077 | 1 |
| $\beta_{G}=\log\left( 3 \right)$=1.0986, $\beta_{O}=\log\left( 2.5 \right)$=0.9163 | | | | | |
| ($\beta_{0},\sigma$) | ${pr}_{B}(D=1$) | Bias | Variance | RMSE | power |
| (-1,1) | 0.3210 | -0.1626 | 0.0023 | 0.0287 | 1 |
| (-1,2) | 0.3620 | -0.3938 | 0.0023 | 0.1573 | 1 |
| (-5,1) | 0.0119 | -0.0100 | 0.0016 | 0.0017 | 1 |
| (-5,2) | 0.0305 | -0.1947 | 0.0017 | 0.0397 | 1 |
| $\beta_{G}=\log\left( 3 \right)$=1.0986, $\beta_{O}=\log\left( 3 \right)$=1.0986 | | | | | |
| ($\beta_{0},\sigma$) | ${pr}_{B}(D=1$) | Bias | Variance | RMSE | power |
| (-1,1) | 0.3296 | -0.2127 | 0.0023 | 0.0475 | 1 |
| (-1,2) | 0.3754 | -0.4668 | 0.0022 | 0.2202 | 1 |
| (-5,1) | 0.0140 | -0.0262 | 0.0017 | 0.0023 | 1 |
| (-5,2) | 0.0439 | -0.2982 | 0.0018 | 0.0907 | 1 |
| $\beta_{G}=\log\left( 3 \right)$=1.0986, $\beta_{O}=\log\left( 5 \right)$=1.6094 | | | | | |
| ($\beta_{0},\sigma$) | ${pr}_{B}(D=1$) | Bias | Variance | RMSE | power |
| (-1,1) | 0.3528 | -0.3421 | 0.0023 | 0.1193 | 1 |
| (-1,2) | 0.4043 | -0.6181 | 0.0022 | 0.3842 | 1 |
| (-5,1) | 0.0240 | -0.1350 | 0.0017 | 0.0199 | 1 |
| (-5,2) | 0.0929 | -0.5334 | 0.0019 | 0.2864 | 1 |
| $\beta_{G}=\log\left( 3 \right)$=1.0986, $\beta_{O}=\log\left( 8 \right)$=2.0794 | | | | | |
| ($\beta_{0},\sigma$) | ${pr}_{B}(D=1$) | Bias | Variance | RMSE | power |
| (-1,1) | 0.3713 | -0.4440 | 0.0022 | 0.1994 | 1 |
| (-1,2) | 0.4221 | -0.7093 | 0.0022 | 0.5053 | 1 |
| (-5,1) | 0.0392 | -0.2657 | 0.0018 | 0.0723 | 1 |
| (-5,2) | 0.1405 | -0.6624 | 0.0020 | 0.4408 | 1 |
| $\beta_{G}=\log\left( 5 \right)$=1.6094 , $\beta_{O}=\log\left( 1 \right)$ | | | | | |
| ($\beta_{0},\sigma$) | ${pr}_{B}(D=1$) | Bias | Variance | RMSE | power |
| (-1,1) | 0.3068 | -0.0007 | 0.0027 | 0.0027 | 1 |
| (-1,2) | 0.3068 | -0.0007 | 0.0027 | 0.0027 | 1 |
| (-5,1) | 0.0092 | 0.0128 | 0.0015 | 0.0017 | 1 |
| (-5,2) | 0.0092 | 0.0128 | 0.0015 | 0.0017 | 1 |
| $\beta_{G}=\log\left( 5 \right)$=1.6094, $\beta_{O}=\log\left( 1.5 \right)=0.4055$ | | | | | |
| ($\beta_{0},\sigma$) | ${pr}_{B}(D=1$) | Bias | Variance | RMSE | power |
| (-1,1) | 0.3126 | -0.059 | 0.0026 | 0.0061 | 1 |
| (-1,2) | 0.3271 | -0.2007 | 0.0026 | 0.0429 | 1 |
| (-5,1) | 0.01 | 0.0081 | 0.0015 | 0.0016 | 1 |
| (-5,2) | 0.0125 | -0.0189 | 0.0016 | 0.0019 | 1 |
| $\beta_{G}=\log\left( 5 \right)$=1.6094, $\beta_{O}=\log\left( 2 \right)$=0.6931 | | | | | |
| ($\beta_{0},\sigma$) | ${pr}_{B}(D=1$) | Bias | Variance | RMSE | power |
| (-1,1) | 0.3223 | -0.1545 | 0.0026 | 0.0265 | 1 |
| (-1,2) | 0.3520 | -0.4271 | 0.0025 | 0.1849 | 1 |
| (-5,1) | 0.0116 | -0.0068 | 0.0016 | 0.0016 | 1 |
| (-5,2) | 0.0210 | -0.1425 | 0.0016 | 0.0219 | 1 |
| $\beta_{G}=\log\left( 5 \right)$=1.6094, $\beta_{O}=\log\left( 2.5 \right)$=0.9163 | | | | | |
| ($\beta_{0},\sigma$) | ${pr}_{B}(D=1$) | Bias | Variance | RMSE | power |
| (-1,1) | 0.3315 | -0.2420 | 0.0026 | 0.0611 | 1 |
| (-1,2) | 0.3700 | -0.5799 | 0.0024 | 0.3388 | 1 |
| (-5,1) | 0.0136 | -0.0292 | 0.0016 | 0.0024 | 1 |
| (-5,2) | 0.0333 | -0.3145 | 0.0017 | 0.1006 | 1 |
| $\beta_{G}=\log\left( 5 \right)$=1.6094, $\beta_{O}=\log\left( 3 \right)$=1.0986 | | | | | |
| ($\beta_{0},\sigma$) | ${pr}_{B}(D=1$) | Bias | Variance | RMSE | power |
| (-1,1) | 0.3395 | -0.3163 | 0.0025 | 0.1026 | 1 |
| (-1,2) | 0.3826 | -0.6863 | 0.0024 | 0.4735 | 1 |
| (-5,1) | 0.0159 | -0.0632 | 0.0016 | 0.0056 | 1 |
| (-5,2) | 0.0470 | -0.4640 | 0.0017 | 0.2170 | 1 |
| $\beta_{G}=\log\left( 5 \right)$=1.6094, $\beta_{O}=\log\left( 5 \right)$=1.6094 | | | | | |
| ($\beta_{0},\sigma$) | ${pr}_{B}(D=1$) | Bias | Variance | RMSE | power |
| (-1,1) | 0.3612 | -0.5068 | 0.0025 | 0.2593 | 1 |
| (-1,2) | 0.4098 | -0.9075 | 0.0023 | 0.8259 | 1 |
| (-5,1) | 0.0265 | -0.2231 | 0.0016 | 0.0514 | 1 |
| (-5,2) | 0.0963 | -0.7903 | 0.0019 | 0.6264 | 1 |
| $\beta_{G}=\log\left( 5 \right)$=1.6094, $\beta_{O}=\log\left( 8 \right)$=2.0794 | | | | | |
| ($\beta_{0},\sigma$) | ${pr}_{B}(D=1$) | Bias | Variance | RMSE | power |
| (-1,1) | 0.3788 | -0.6532 | 0.0024 | 0.4291 | 1 |
| (-1,2) | 0.4266 | -1.0406 | 0.0023 | 1.0852 | 1 |
| (-5,1) | 0.0422 | -0.4169 | 0.0017 | 0.1755 | 1 |
| (-5,2) | 0.1438 | -0.9750 | 0.0020 | 0.9525 | 1 |
| $\beta_{G}=\log\left( 8 \right)$=2.0794, $\beta_{O}=\log\left( 1 \right)$ | | | | | |
| ($\beta_{0},\sigma$) | ${pr}_{B}(D=1$) | Bias | Variance | RMSE | power |
| (-1,1) | 0.3169 | 0.0014 | 0.0035 | 0.0035 | 1 |
| (-1,2) | 0.3167 | -0.0004 | 0.0035 | 0.0035 | 1 |
| (-5,1) | 0.0111 | 0.0108 | 0.0015 | 0.0017 | 1 |
| (-5,2) | 0.0111 | 0.0108 | 0.0015 | 0.0017 | 1 |
| $\beta_{G}=\log\left( 8 \right)$=2.0794, $\beta_{O}=\log\left( 1.5 \right)=0.4055$ | | | | | |
| ($\beta_{0},\sigma$) | ${pr}_{B}(D=1$) | Bias | Variance | RMSE | power |
| (-1,1) | 0.3222 | -0.0742 | 0.0032 | 0.0087 | 1 |
| (-1,2) | 0.3362 | -0.2511 | 0.003 | 0.066 | 1 |
| (-5,1) | 0.012 | 0.0028 | 0.0016 | 0.0016 | 1 |
| (-5,2) | 0.0149 | -0.0368 | 0.0016 | 0.003 | 1 |
| $\beta_{G}=\log\left( 8 \right)$=2.0794, $\beta_{O}=\log\left( 2 \right)$=0.6931 | | | | | |
| ($\beta_{0},\sigma$) | ${pr}_{B}(D=1$) | Bias | Variance | RMSE | power |
| (-1,1) | 0.3317 | -0.1917 | 0.0033 | 0.0400 | 1 |
| (-1,2) | 0.3600 | -0.5462 | 0.0029 | 0.3012 | 1 |
| (-5,1) | 0.0138 | -0.0204 | 0.0016 | 0.0020 | 1 |
| (-5,2) | 0.0240 | -0.2104 | 0.0016 | 0.0459 | 1 |
| $\beta_{G}=\log\left( 8 \right)$=2.0794, $\beta_{O}=\log\left( 2.5 \right)$=0.9163 | | | | | |
| ($\beta_{0},\sigma$) | ${pr}_{B}(D=1$) | Bias | Variance | RMSE | power |
| (-1,1) | 0.3407 | -0.3064 | 0.0031 | 0.0970 | 1 |
| (-1,2) | 0.3770 | -0.7472 | 0.0027 | 0.5611 | 1 |
| (-5,1) | 0.0160 | -0.0579 | 0.0016 | 0.0049 | 1 |
| (-5,2) | 0.0366 | -0.4383 | 0.0017 | 0.1938 | 1 |
| $\beta_{G}=\log\left( 8 \right)$=2.0794, $\beta_{O}=\log\left( 3 \right)$=1.0986 | | | | | |
| ($\beta_{0},\sigma$) | ${pr}_{B}(D=1$) | Bias | Variance | RMSE | power |
| (-1,1) | 0.3483 | -0.3981 | 0.0030 | 0.1615 | 1 |
| (-1,2) | 0.3890 | -0.8862 | 0.0026 | 0.7879 | 1 |
| (-5,1) | 0.0185 | -0.1045 | 0.0016 | 0.0125 | 1 |
| (-5,2) | 0.0505 | -0.6276 | 0.0017 | 0.3956 | 1 |
| $\beta_{G}=\log\left( 8 \right)$=2.0794, $\beta_{O}=\log\left( 5 \right)$=1.6094 | | | | | |
| ($\beta_{0},\sigma$) | ${pr}_{B}(D=1$) | Bias | Variance | RMSE | power |
| (-1,1) | 0.3687 | -0.6513 | 0.0028 | 0.4270 | 1 |
| (-1,2) | 0.4147 | -1.1727 | 0.0025 | 1.3777 | 1 |
| (-5,1) | 0.0297 | -0.3203 | 0.0016 | 0.1042 | 1 |
| (-5,2) | 0.0998 | -1.0364 | 0.0019 | 1.0761 | 1 |
| $\beta_{G}=\log\left( 8 \right)$=2.0794, $\beta_{O}=\log\left( 8 \right)$=2.0794 | | | | | |
| ($\beta_{0},\sigma$) | ${pr}_{B}(D=1$) | Bias | Variance | RMSE | power |
| (-1,1) | 0.3855 | -0.8444 | 0.0026 | 0.7156 | 1 |
| (-1,2) | 0.4306 | -1.3449 | 0.0024 | 1.8111 | 1 |
| (-5,1) | 0.0457 | -0.5678 | 0.0017 | 0.3241 | 1 |
| (-5,2) | 0.1471 | -1.2681 | 0.0020 | 1.6101 | 1 |

**Supplementary Table 3**: Probability of the disease in the population (${pr}_{B}(D=1$)), Bias, Variance, Root Mean Squared Error of the estimates of main effects of the genotype from misspecified model (2), False Discovery Rate (FDR). The genotype is simulated to be Bernoulli(0.1), the omitted variable is simulated from Normal(0,$\sigma^{2}$). In the true model (1) the parameters are $\beta_{0}=-1,-5; \beta_{G}=\log\left( 1 \right),\log\left( 1.5 \right),\log\left( 2 \right),\log\left( 2.5 \right),\log\left( 3 \right),\log\left( 5 \right),log(8), \beta_{O}=\log\left( 1 \right),\log\left( 1.5 \right),\log\left( 2 \right),\log\left( 2.5 \right),\log\left( 3 \right),\log\left( 5 \right),\log\left( 8 \right).$ The results are based on 5,000 datasets of 10,000 cases and 10,000 controls.

| $\beta_{O}=\log\left( 1 \right),\mu_{G}=\log\left( 1 \right)$ | | | | | | | |
| --- | --- | --- | --- | --- | --- | --- | --- |
| ($\beta_{0},\sigma$) | ${pr}_{B}(D=d$) | Ranks based on estimates | | | Ranks based on p values | | |
|  |  | ALL | TOP 10% | TOP 20% | ALL | TOP 10% | TOP 20% |
| (-1,1) | 0.3025 | 0.9982 | 1 | 0.9999 | 0.9938 | 0.997 | 0.998 |
| (-1,2) | 0.302 | 0.9979 | 1 | 1 | 0.9936 | 0.9966 | 0.998 |
| (-5,1) | 0.0106 | 0.9982 | 1 | 1 | 0.9945 | 0.999 | 0.998 |
| (-5,2) | 0.0106 | 0.9982 | 1 | 1 | 0.9945 | 0.999 | 0.998 |
| $\beta_{O}=\log\left( 1 \right),\mu_{G}=\log\left( 2 \right)=0693$ | | | | | | | |
| ($\beta_{0},\sigma$) | ${pr}_{B}(D=d$) | Ranks based on estimates | | | Ranks based on p values | | |
|  |  | ALL | TOP 10% | TOP 20% | ALL | TOP 10% | TOP 20% |
| (-1,1) | 0.5715 | 0.9964 | 0.9972 | 0.9977 | 0.9971 | 0.9958 | 0.9967 |
| (-1,2) | 0.5715 | 0.996 | 0.997 | 0.9978 | 0.9972 | 0.9962 | 0.9969 |
| (-5,1) | 0.048 | 0.9971 | 0.9976 | 0.9981 | 0.9969 | 0.9952 | 0.9954 |
| (-5,2) | 0.048 | 0.9971 | 0.9976 | 0.9981 | 0.9969 | 0.9952 | 0.9954 |
| $\beta_{O}=\log\left( 1 \right),\mu_{G}=\log\left( 3 \right)=1.099$ | | | | | | | |
| ($\beta_{0},\sigma$) | ${pr}_{B}(D=d$) | Ranks based on estimates | | | Ranks based on p values | | |
|  |  | ALL | TOP 10% | TOP 20% | ALL | TOP 10% | TOP 20% |
| (-1,1) | 0.6762 | 0.9955 | 0.9966 | 0.9968 | 0.996 | 0.998 | 0.9988 |
| (-1,2) | 0.6762 | 0.9962 | 0.997 | 0.998 | 0.9957 | 0.9982 | 0.999 |
| (-5,1) | 0.1117 | 0.9955 | 0.9964 | 0.9967 | 0.9981 | 0.9956 | 0.9978 |
| (-5,2) | 0.1117 | 0.9955 | 0.9964 | 0.9967 | 0.9981 | 0.9956 | 0.9978 |
| $\beta_{O}=\log\left( 1 \right),\mu_{G}=\log\left( 5 \right)=1.609$ | | | | | | | |
| ($\beta_{0},\sigma$) | ${pr}_{B}(D=d$) | Ranks based on estimates | | | Ranks based on p values | | |
|  |  | ALL | TOP 10% | TOP 20% | ALL | TOP 10% | TOP 20% |
| (-1,1) | 0.7609 | 0.9942 | 0.9964 | 0.9973 | 0.9952 | 0.998 | 0.9937 |
| (-1,2) | 0.7609 | 0.9946 | 0.9966 | 0.997 | 0.9947 | 0.9974 | 0.993 |
| (-5,1) | 0.2283 | 0.9956 | 0.9972 | 0.9974 | 0.9983 | 0.9982 | 0.9991 |
| (-5,2) | 0.2283 | 0.9956 | 0.9972 | 0.9974 | 0.9983 | 0.9982 | 0.9991 |
| $\beta_{O}=\log\left( 1 \right),\mu_{G}=\log\left( 8 \right)=2.079$ | | | | | | | |
| ($\beta_{0},\sigma$) | ${pr}_{B}(D=d$) | Ranks based on estimates | | | Ranks based on p values | | |
|  |  | ALL | TOP 10% | TOP 20% | ALL | TOP 10% | TOP 20% |
| (-1,1) | 0.7997 | 0.9941 | 0.9934 | 0.9923 | 0.9968 | 0.9968 | 0.9975 |
| (-1,2) | 0.8007 | 0.996 | 0.9952 | 0.994 | 0.9962 | 0.9962 | 0.9973 |
| (-5,1) | 0.3177 | 0.9962 | 0.9952 | 0.9948 | 0.9959 | 0.9936 | 0.9953 |
| (-5,2) | 0.3177 | 0.9962 | 0.9952 | 0.9948 | 0.9959 | 0.9936 | 0.9953 |

| $\beta_{O}=\log\left( 1.5 \right)=0.405,\mu_{G}=\log\left( 1 \right)$ | | | | | | | |
| --- | --- | --- | --- | --- | --- | --- | --- |
| ($\beta_{0},\sigma$) | ${pr}_{B}(D=d$) | Ranks based on estimates | | | Ranks based on p values | | |
|  |  | ALL | TOP 10% | TOP 20% | ALL | TOP 10% | TOP 20% |
| (-1,1) | 0.3276 | 0.944 | 0.9594 | 0.9647 | 0.8578 | 0.842 | 0.8166 |
| (-1,2) | 0.3398 | 0.8881 | 0.9262 | 0.9347 | 0.7444 | 0.7338 | 0.6804 |
| (-5,1) | 0.0105 | 0.9458 | 0.9436 | 0.9582 | 0.8558 | 0.8126 | 0.8126 |
| (-5,2) | 0.0131 | 0.8949 | 0.8786 | 0.9129 | 0.7173 | 0.6246 | 0.6214 |
| $\beta_{O}=\log\left( 1.5 \right)=0.405,\mu_{G}=\log\left( 2 \right)=0.693$ | | | | | | | |
| ($\beta_{0},\sigma$) | ${pr}_{B}(D=d$) | Ranks based on estimates | | | Ranks based on p values | | |
|  |  | ALL | TOP 10% | TOP 20% | ALL | TOP 10% | TOP 20% |
| (-1,1) | 0.5706 | 0.9344 | 0.9462 | 0.9585 | 0.9494 | 0.914 | 0.9352 |
| (-1,2) | 0.5672 | 0.8804 | 0.9084 | 0.929 | 0.8964 | 0.815 | 0.8536 |
| (-5,1) | 0.0507 | 0.9486 | 0.9538 | 0.9649 | 0.9512 | 0.9144 | 0.9253 |
| (-5,2) | 0.0584 | 0.8973 | 0.9086 | 0.93 | 0.9021 | 0.8246 | 0.8559 |
| $\beta_{O}=\log\left( 1.5 \right)=0.405,\mu_{G}=\log\left( 3 \right)=1.099$ | | | | | | | |
| ($\beta_{0},\sigma$) | ${pr}_{B}(D=d$) | Ranks based on estimates | | | Ranks based on p values | | |
|  |  | ALL | TOP 10% | TOP 20% | ALL | TOP 10% | TOP 20% |
| (-1,1) | 0.6741 | 0.932 | 0.9484 | 0.9604 | 0.9426 | 0.9802 | 0.9876 |
| (-1,2) | 0.6673 | 0.8688 | 0.8984 | 0.9178 | 0.9009 | 0.9596 | 0.9739 |
| (-5,1) | 0.1149 | 0.9425 | 0.95 | 0.9601 | 0.9715 | 0.9262 | 0.9618 |
| (-5,2) | 0.1239 | 0.885 | 0.8896 | 0.9135 | 0.9411 | 0.8266 | 0.9116 |
| $\beta_{O}=\log\left( 1.5 \right)=0.405,\mu_{G}=\log\left( 5 \right)=1.609$ | | | | | | | |
| ($\beta_{0},\sigma$) | ${pr}_{B}(D=d$) | Ranks based on estimates | | | Ranks based on p values | | |
|  |  | ALL | TOP 10% | TOP 20% | ALL | TOP 10% | TOP 20% |
| (-1,1) | 0.7588 | 0.9211 | 0.9406 | 0.9524 | 0.9265 | 0.9614 | 0.908 |
| (-1,2) | 0.752 | 0.8534 | 0.877 | 0.9053 | 0.8656 | 0.9338 | 0.8333 |
| (-5,1) | 0.2309 | 0.9352 | 0.9492 | 0.9609 | 0.9667 | 0.9802 | 0.9893 |
| (-5,2) | 0.2374 | 0.8762 | 0.8814 | 0.9065 | 0.9351 | 0.9576 | 0.9746 |
| $\beta_{O}=\log\left( 1.5 \right)=0.405,\mu_{G}=\log\left( 8 \right)=2.079$ | | | | | | | |
| ($\beta_{0},\sigma$) | ${pr}_{B}(D=d$) | Ranks based on estimates | | | Ranks based on p values | | |
|  |  | ALL | TOP 10% | TOP 20% | ALL | TOP 10% | TOP 20% |
| (-1,1) | 0.8068 | 0.9052 | 0.9156 | 0.939 | 0.9451 | 0.9438 | 0.9592 |
| (-1,2) | 0.8009 | 0.8367 | 0.8274 | 0.8749 | 0.8934 | 0.8936 | 0.9119 |
| (-5,1) | 0.3416 | 0.9328 | 0.9462 | 0.9605 | 0.9526 | 0.9866 | 0.9567 |
| (-5,2) | 0.3455 | 0.8702 | 0.8802 | 0.9103 | 0.9088 | 0.9708 | 0.9097 |

| $\beta_{O}=\log\left( 2 \right)=0.693,\mu_{G}=\log\left( 1 \right)$ | | | | | | | |
| --- | --- | --- | --- | --- | --- | --- | --- |
| ($\beta_{0},\sigma$) | ${pr}_{B}(D=d$) | Ranks based on estimates | | | Ranks based on p values | | |
|  |  | ALL | TOP 10% | TOP 20% | ALL | TOP 10% | TOP 20% |
| (-1,1) | 0.3356 | 0.9045 | 0.9344 | 0.9431 | 0.7735 | 0.7626 | 0.7114 |
| (-1,2) | 0.361 | 0.8117 | 0.8378 | 0.8646 | 0.6129 | 0.5794 | 0.5175 |
| (-5,1) | 0.0121 | 0.9113 | 0.8956 | 0.927 | 0.7602 | 0.6772 | 0.6771 |
| (-5,2) | 0.0222 | 0.8133 | 0.8358 | 0.8638 | 0.5717 | 0.505 | 0.4855 |
| $\beta_{O}=\log\left( 2 \right)=0.693,\mu_{G}=\log\left( 2 \right)=0.693$ | | | | | | | |
| ($\beta_{0},\sigma$) | ${pr}_{B}(D=d$) | Ranks based on estimates | | | Ranks based on p values | | |
|  |  | ALL | TOP 10% | TOP 20% | ALL | TOP 10% | TOP 20% |
| (-1,1) | 0.5681 | 0.8928 | 0.9244 | 0.9411 | 0.9101 | 0.8472 | 0.8764 |
| (-1,2) | 0.5605 | 0.8024 | 0.841 | 0.866 | 0.8135 | 0.6812 | 0.7467 |
| (-5,1) | 0.0557 | 0.9107 | 0.9098 | 0.9311 | 0.9171 | 0.8532 | 0.872 |
| (-5,2) | 0.0783 | 0.8276 | 0.8316 | 0.872 | 0.8305 | 0.6942 | 0.7466 |
| $\beta_{O}=\log\left( 2 \right)=0.693,\mu_{G}=\log\left( 3 \right)=1.099$ | | | | | | | |
| ($\beta_{0},\sigma$) | ${pr}_{B}(D=d$) | Ranks based on estimates | | | Ranks based on p values | | |
|  |  | ALL | TOP 10% | TOP 20% | ALL | TOP 10% | TOP 20% |
| (-1,1) | 0.6697 | 0.8872 | 0.9146 | 0.9313 | 0.9086 | 0.9586 | 0.9752 |
| (-1,2) | 0.6537 | 0.7933 | 0.826 | 0.8632 | 0.842 | 0.8942 | 0.9317 |
| (-5,1) | 0.1207 | 0.9002 | 0.903 | 0.9237 | 0.9502 | 0.858 | 0.9277 |
| (-5,2) | 0.145 | 0.8112 | 0.7968 | 0.8429 | 0.8907 | 0.7238 | 0.8554 |
| $\beta_{O}=\log\left( 2 \right)=0.693,\mu_{G}=\log\left( 5 \right)=1.609$ | | | | | | | |
| ($\beta_{0},\sigma$) | ${pr}_{B}(D=d$) | Ranks based on estimates | | | Ranks based on p values | | |
|  |  | ALL | TOP 10% | TOP 20% | ALL | TOP 10% | TOP 20% |
| (-1,1) | 0.7541 | 0.8674 | 0.8836 | 0.9094 | 0.8835 | 0.9436 | 0.8541 |
| (-1,2) | 0.737 | 0.7725 | 0.8054 | 0.8448 | 0.8 | 0.8914 | 0.7729 |
| (-5,1) | 0.2351 | 0.8928 | 0.9018 | 0.9259 | 0.9437 | 0.9652 | 0.9801 |
| (-5,2) | 0.2527 | 0.8083 | 0.8208 | 0.8565 | 0.8857 | 0.9254 | 0.9475 |
| $\beta_{O}=\log\left( 2 \right)=0.693,\mu_{G}=\log\left( 8 \right)=2.079$ | | | | | | | |
| ($\beta_{0},\sigma$) | ${pr}_{B}(D=d$) | Ranks based on estimates | | | Ranks based on p values | | |
|  |  | ALL | TOP 10% | TOP 20% | ALL | TOP 10% | TOP 20% |
| (-1,1) | 0.8026 | 0.8509 | 0.8632 | 0.8968 | 0.9077 | 0.9186 | 0.9338 |
| (-1,2) | 0.7881 | 0.7571 | 0.7508 | 0.8112 | 0.8269 | 0.8322 | 0.8276 |
| (-5,1) | 0.3444 | 0.8883 | 0.8972 | 0.926 | 0.9198 | 0.97 | 0.9215 |
| (-5,2) | 0.354 | 0.7886 | 0.8126 | 0.8444 | 0.8503 | 0.945 | 0.8534 |

| $\beta_{O}=\log\left( 2.5 \right)=0.916,\mu_{G}=\log\left( 1 \right)$ | | | | | | | |
| --- | --- | --- | --- | --- | --- | --- | --- |
| ($\beta_{0},\sigma$) | ${pr}_{B}(D=d$) | Ranks based on estimates | | | Ranks based on p values | | |
|  |  | ALL | TOP 10% | TOP 20% | ALL | TOP 10% | TOP 20% |
| (-1,1) | 0.3243 | 0.9051 | 0.9984 | 0.9931 | 0.7761 | 0.9026 | 0.9087 |
| (-1,2) | 0.3618 | 0.8099 | 0.9906 | 0.9664 | 0.6237 | 0.8318 | 0.7896 |
| (-5,1) | 0.0151 | 0.9082 | 0.997 | 0.9917 | 0.7725 | 0.909 | 0.8943 |
| (-5,2) | 0.0349 | 0.8264 | 0.988 | 0.9713 | 0.6156 | 0.7468 | 0.7683 |
| $\beta_{O}=\log\left( 2.5 \right)=0.916,\mu_{G}=\log\left( 2 \right)=0.693$ | | | | | | | |
| ($\beta_{0},\sigma$) | ${pr}_{B}(D=d$) | Ranks based on estimates | | | Ranks based on p values | | |
|  |  | ALL | TOP 10% | TOP 20% | ALL | TOP 10% | TOP 20% |
| (-1,1) | 0.566 | 0.8673 | 0.9006 | 0.9252 | 0.8785 | 0.7814 | 0.8298 |
| (-1,2) | 0.555 | 0.7508 | 0.7898 | 0.819 | 0.7476 | 0.5788 | 0.6649 |
| (-5,1) | 0.0613 | 0.8799 | 0.8878 | 0.9078 | 0.8883 | 0.7958 | 0.830 |
| (-5,2) | 0.0996 | 0.7815 | 0.7836 | 0.8349 | 0.7825 | 0.6134 | 0.6811 |
| $\beta_{O}=\log\left( 2.5 \right)=0.916,\mu_{G}=\log\left( 3 \right)=1.099$ | | | | | | | |
| ($\beta_{0},\sigma$) | ${pr}_{B}(D=d$) | Ranks based on estimates | | | Ranks based on p values | | |
|  |  | ALL | TOP 10% | TOP 20% | ALL | TOP 10% | TOP 20% |
| (-1,1) | 0.6655 | 0.8488 | 0.8768 | 0.9046 | 0.8844 | 0.9378 | 0.9614 |
| (-1,2) | 0.6415 | 0.7398 | 0.7694 | 0.8126 | 0.7976 | 0.8158 | 0.8821 |
| (-5,1) | 0.1271 | 0.8682 | 0.8652 | 0.8985 | 0.9302 | 0.81 | 0.9028 |
| (-5,2) | 0.1655 | 0.7592 | 0.751 | 0.7965 | 0.8486 | 0.6768 | 0.8258 |
| $\beta_{O}=\log\left( 2.5 \right)=0.916,\mu_{G}=\log\left( 5 \right)=1.609$ | | | | | | | |
| ($\beta_{0},\sigma$) | ${pr}_{B}(D=d$) | Ranks based on estimates | | | Ranks based on p values | | |
|  |  | ALL | TOP 10% | TOP 20% | ALL | TOP 10% | TOP 20% |
| (-1,1) | 0.7498 | 0.8362 | 0.8494 | 0.8841 | 0.8533 | 0.92 | 0.8232 |
| (-1,2) | 0.7226 | 0.7259 | 0.7546 | 0.7998 | 0.7541 | 0.861 | 0.7561 |
| (-5,1) | 0.2397 | 0.8672 | 0.872 | 0.902 | 0.9272 | 0.9568 | 0.9702 |
| (-5,2) | 0.2677 | 0.751 | 0.7628 | 0.8013 | 0.8401 | 0.8878 | 0.9183 |
| $\beta_{O}=\log\left( 2.5 \right)=0.916,\mu_{G}=\log\left( 8 \right)=2.079$ | | | | | | | |
| ($\beta_{0},\sigma$) | ${pr}_{B}(D=d$) | Ranks based on estimates | | | Ranks based on p values | | |
|  |  | ALL | TOP 10% | TOP 20% | ALL | TOP 10% | TOP 20% |
| (-1,1) | 0.7986 | 0.8126 | 0.8286 | 0.8682 | 0.8793 | 0.8838 | 0.8995 |
| (-1,2) | 0.7748 | 0.7069 | 0.7076 | 0.7647 | 0.7804 | 0.819 | 0.7475 |
| (-5,1) | 0.3468 | 0.8545 | 0.8746 | 0.9042 | 0.8962 | 0.9658 | 0.8934 |
| (-5,2) | 0.3628 | 0.7433 | 0.7522 | 0.7989 | 0.8026 | 0.911 | 0.7982 |

| $\beta_{O}=\log\left( 3 \right)=1.099,\mu_{G}=\log\left( 1 \right)$ | | | | | | | |
| --- | --- | --- | --- | --- | --- | --- | --- |
| ($\beta_{0},\sigma$) | ${pr}_{B}(D=d$) | Ranks based on estimates | | | Ranks based on p values | | |
|  |  | ALL | TOP 10% | TOP 20% | ALL | TOP 10% | TOP 20% |
| (-1,1) | 0.3321 | 0.8864 | 0.9972 | 0.9868 | 0.7402 | 0.887 | 0.888 |
| (-1,2) | 0.3744 | 0.7693 | 0.9768 | 0.9332 | 0.5785 | 0.8144 | 0.746 |
| (-5,1) | 0.0176 | 0.8912 | 0.9968 | 0.9905 | 0.7458 | 0.8912 | 0.8726 |
| (-5,2) | 0.0483 | 0.7935 | 0.9752 | 0.9525 | 0.5673 | 0.6922 | 0.7145 |
| $\beta_{O}=\log\left( 3 \right)=1.099,\mu_{G}=\log\left( 2 \right)=0.693$ | | | | | | | |
| ($\beta_{0},\sigma$) | ${pr}_{B}(D=d$) | Ranks based on estimates | | | Ranks based on p values | | |
|  |  | ALL | TOP 10% | TOP 20% | ALL | TOP 10% | TOP 20% |
| (-1,1) | 0.564 | 0.8404 | 0.8718 | 0.9 | 0.852 | 0.7366 | 0.7933 |
| (-1,2) | 0.551 | 0.7127 | 0.7276 | 0.7716 | 0.6962 | 0.5332 | 0.6126 |
| (-5,1) | 0.0671 | 0.8606 | 0.8668 | 0.8995 | 0.864 | 0.7494 | 0.7964 |
| (-5,2) | 0.1186 | 0.7382 | 0.7322 | 0.7863 | 0.7285 | 0.5346 | 0.6195 |
| $\beta_{O}=\log\left( 3 \right)=1.099,\mu_{G}=\log\left( 3 \right)=1.099$ | | | | | | | |
| ($\beta_{0},\sigma$) | ${pr}_{B}(D=d$) | Ranks based on estimates | | | Ranks based on p values | | |
|  |  | ALL | TOP 10% | TOP 20% | ALL | TOP 10% | TOP 20% |
| (-1,1) | 0.6613 | 0.8225 | 0.8508 | 0.8814 | 0.8667 | 0.9226 | 0.9499 |
| (-1,2) | 0.6319 | 0.7041 | 0.7252 | 0.7737 | 0.7599 | 0.7424 | 0.8303 |
| (-5,1) | 0.1334 | 0.8473 | 0.8382 | 0.8771 | 0.9152 | 0.77 | 0.8808 |
| (-5,2) | 0.1836 | 0.7261 | 0.728 | 0.7743 | 0.8045 | 0.6194 | 0.7902 |
| $\beta_{O}=\log\left( 3 \right)=1.099,\mu_{G}=\log\left( 5 \right)=1.609$ | | | | | | | |
| ($\beta_{0},\sigma$) | ${pr}_{B}(D=d$) | Ranks based on estimates | | | Ranks based on p values | | |
|  |  | ALL | TOP 10% | TOP 20% | ALL | TOP 10% | TOP 20% |
| (-1,1) | 0.7451 | 0.8147 | 0.8336 | 0.8703 | 0.8361 | 0.9114 | 0.8068 |
| (-1,2) | 0.7106 | 0.692 | 0.7128 | 0.7637 | 0.7199 | 0.8384 | 0.7362 |
| (-5,1) | 0.2441 | 0.8408 | 0.859 | 0.8875 | 0.9076 | 0.9388 | 0.9609 |
| (-5,2) | 0.2808 | 0.713 | 0.7194 | 0.7666 | 0.8043 | 0.8472 | 0.8797 |
| $\beta_{O}=\log\left( 3 \right)=1.099,\mu_{G}=\log\left( 8 \right)=2.079$ | | | | | | | |
| ($\beta_{0},\sigma$) | ${pr}_{B}(D=d$) | Ranks based on estimates | | | Ranks based on p values | | |
|  |  | ALL | TOP 10% | TOP 20% | ALL | TOP 10% | TOP 20% |
| (-1,1) | 0.7948 | 0.786 | 0.7946 | 0.8422 | 0.8605 | 0.8708 | 0.8762 |
| (-1,2) | 0.763 | 0.6692 | 0.6602 | 0.7241 | 0.7398 | 0.7812 | 0.6765 |
| (-5,1) | 0.3496 | 0.8347 | 0.8552 | 0.8884 | 0.8779 | 0.961 | 0.8796 |
| (-5,2) | 0.3706 | 0.7083 | 0.712 | 0.7698 | 0.7704 | 0.8922 | 0.7705 |

| $\beta_{O}=\log\left( 5 \right)=1.609,\mu_{G}=\log\left( 1 \right)$ | | | | | | | |
| --- | --- | --- | --- | --- | --- | --- | --- |
| ($\beta_{0},\sigma$) | ${pr}_{B}(D=d$) | Ranks based on estimates | | | Ranks based on p values | | |
|  |  | ALL | TOP 10% | TOP 20% | ALL | TOP 10% | TOP 20% |
| (-1,1) | 0.3696 | 0.7811 | 0.8398 | 0.8532 | 0.5759 | 0.5534 | 0.4854 |
| (-1,2) | 0.4135 | 0.6227 | 0.571 | 0.6497 | 0.3813 | 0.3038 | 0.275 |
| (-5,1) | 0.028 | 0.7903 | 0.8218 | 0.8432 | 0.5323 | 0.486 | 0.4548 |
| (-5,2) | 0.0992 | 0.628 | 0.6174 | 0.6645 | 0.3607 | 0.2958 | 0.2742 |
| $\beta_{O}=\log\left( 5 \right)=1.609,\mu_{G}=\log\left( 2 \right)=0.693$ | | | | | | | |
| ($\beta_{0},\sigma$) | ${pr}_{B}(D=d$) | Ranks based on estimates | | | Ranks based on p values | | |
|  |  | ALL | TOP 10% | TOP 20% | ALL | TOP 10% | TOP 20% |
| (-1,1) | 0.5579 | 0.7738 | 0.8208 | 0.8523 | 0.7778 | 0.6204 | 0.704 |
| (-1,2) | 0.5402 | 0.6177 | 0.566 | 0.6445 | 0.5652 | 0.4066 | 0.4889 |
| (-5,1) | 0.0883 | 0.8027 | 0.7946 | 0.8451 | 0.8077 | 0.6646 | 0.7162 |
| (-5,2) | 0.1747 | 0.6389 | 0.62 | 0.676 | 0.6038 | 0.4316 | 0.5073 |
| $\beta_{O}=\log\left( 5 \right)=1.609,\mu_{G}=\log\left( 3 \right)=1.099$ | | | | | | | |
| ($\beta_{0},\sigma$) | ${pr}_{B}(D=d$) | Ranks based on estimates | | | Ranks based on p values | | |
|  |  | ALL | TOP 10% | TOP 20% | ALL | TOP 10% | TOP 20% |
| (-1,1) | 0.6478 | 0.7655 | 0.7968 | 0.8364 | 0.8175 | 0.858 | 0.908 |
| (-1,2) | 0.6077 | 0.6144 | 0.5728 | 0.6477 | 0.6586 | 0.5552 | 0.6895 |
| (-5,1) | 0.1549 | 0.7844 | 0.7828 | 0.8256 | 0.8755 | 0.7072 | 0.8433 |
| (-5,2) | 0.2335 | 0.6384 | 0.6274 | 0.681 | 0.701 | 0.521 | 0.7035 |
| $\beta_{O}=\log\left( 5 \right)=1.609,\mu_{G}=\log\left( 5 \right)=1.609$ | | | | | | | |
| ($\beta_{0},\sigma$) | ${pr}_{B}(D=d$) | Ranks based on estimates | | | Ranks based on p values | | |
|  |  | ALL | TOP 10% | TOP 20% | ALL | TOP 10% | TOP 20% |
| (-1,1) | 0.7301 | 0.7486 | 0.7746 | 0.8147 | 0.7738 | 0.8802 | 0.766 |
| (-1,2) | 0.6785 | 0.6076 | 0.6166 | 0.6686 | 0.6481 | 0.74 | 0.6693 |
| (-5,1) | 0.26 | 0.7753 | 0.8058 | 0.8349 | 0.8603 | 0.9002 | 0.9285 |
| (-5,2) | 0.3153 | 0.6187 | 0.6006 | 0.6574 | 0.7078 | 0.7168 | 0.7533 |
| $\beta_{O}=\log\left( 5 \right)=1.609,\mu_{G}=\log\left( 8 \right)=2.079$ | | | | | | | |
| ($\beta_{0},\sigma$) | ${pr}_{B}(D=d$) | Ranks based on estimates | | | Ranks based on p values | | |
|  |  | ALL | TOP 10% | TOP 20% | ALL | TOP 10% | TOP 20% |
| (-1,1) | 0.7816 | 0.7257 | 0.7246 | 0.7797 | 0.8036 | 0.8372 | 0.7848 |
| (-1,2) | 0.7301 | 0.5956 | 0.5804 | 0.6482 | 0.6579 | 0.7316 | 0.5681 |
| (-5,1) | 0.3582 | 0.7673 | 0.7844 | 0.8249 | 0.8255 | 0.923 | 0.8189 |
| (-5,2) | 0.3912 | 0.612 | 0.586 | 0.6553 | 0.681 | 0.7748 | 0.6591 |

| $\beta_{O}=\log\left( 8 \right)=2.079,\mu_{G}=\log\left( 1 \right)$ | | | | | | | |
| --- | --- | --- | --- | --- | --- | --- | --- |
| ($\beta_{0},\sigma$) | ${pr}_{B}(D=d$) | Ranks based on estimates | | | Ranks based on p values | | |
|  |  | ALL | TOP 10% | TOP 20% | ALL | TOP 10% | TOP 20% |
| (-1,1) | 0.3856 | 0.7273 | 0.7668 | 0.7981 | 0.5042 | 0.4634 | 0.4122 |
| (-1,2) | 0.4291 | 0.5526 | 0.4922 | 0.5687 | 0.3116 | 0.2642 | 0.2369 |
| (-5,1) | 0.0443 | 0.7363 | 0.7944 | 0.8093 | 0.4615 | 0.436 | 0.3875 |
| (-5,2) | 0.1465 | 0.5695 | 0.5546 | 0.6094 | 0.3105 | 0.2792 | 0.246 |
| $\beta_{O}=\log\left( 8 \right)=2.079,\mu_{G}=\log\left( 2 \right)=0.693$ | | | | | | | |
| ($\beta_{0},\sigma$) | ${pr}_{B}(D=d$) | Ranks based on estimates | | | Ranks based on p values | | |
|  |  | ALL | TOP 10% | TOP 20% | ALL | TOP 10% | TOP 20% |
| (-1,1) | 0.5522 | 0.7231 | 0.7444 | 0.7929 | 0.7031 | 0.5386 | 0.6239 |
| (-1,2) | 0.5336 | 0.546 | 0.4578 | 0.5575 | 0.4777 | 0.3456 | 0.4151 |
| (-5,1) | 0.1124 | 0.7477 | 0.736 | 0.7967 | 0.745 | 0.562 | 0.6372 |
| (-5,2) | 0.22 | 0.5617 | 0.5308 | 0.5909 | 0.5064 | 0.3662 | 0.4308 |
| $\beta_{O}=\log\left( 8 \right)=2.079,\mu_{G}=\log\left( 3 \right)=1.099$ | | | | | | | |
| ($\beta_{0},\sigma$) | ${pr}_{B}(D=d$) | Ranks based on estimates | | | Ranks based on p values | | |
|  |  | ALL | TOP 10% | TOP 20% | ALL | TOP 10% | TOP 20% |
| (-1,1) | 0.6353 | 0.7218 | 0.7332 | 0.7901 | 0.7715 | 0.757 | 0.8454 |
| (-1,2) | 0.5905 | 0.544 | 0.464 | 0.5568 | 0.571 | 0.4378 | 0.5733 |
| (-5,1) | 0.1778 | 0.7322 | 0.731 | 0.7793 | 0.824 | 0.6584 | 0.8134 |
| (-5,2) | 0.2719 | 0.5553 | 0.4806 | 0.566 | 0.5998 | 0.424 | 0.5881 |
| $\beta_{O}=\log\left( 8 \right)=2.079,\mu_{G}=\log\left( 5 \right)=1.609$ | | | | | | | |
| ($\beta_{0},\sigma$) | ${pr}_{B}(D=d$) | Ranks based on estimates | | | Ranks based on p values | | |
|  |  | ALL | TOP 10% | TOP 20% | ALL | TOP 10% | TOP 20% |
| (-1,1) | 0.7148 | 0.6976 | 0.6894 | 0.7475 | 0.7276 | 0.832 | 0.7267 |
| (-1,2) | 0.6542 | 0.5467 | 0.5278 | 0.5881 | 0.5932 | 0.621 | 0.606 |
| (-5,1) | 0.2767 | 0.7275 | 0.7386 | 0.7892 | 0.8149 | 0.8574 | 0.8918 |
| (-5,2) | 0.3426 | 0.5489 | 0.492 | 0.5731 | 0.6248 | 0.5758 | 0.6412 |
| $\beta_{O}=\log\left( 8 \right)=2.079,\mu_{G}=\log\left( 8 \right)=2.079$ | | | | | | | |
| ($\beta_{0},\sigma$) | ${pr}_{B}(D=d$) | Ranks based on estimates | | | Ranks based on p values | | |
|  |  | ALL | TOP 10% | TOP 20% | ALL | TOP 10% | TOP 20% |
| (-1,1) | 0.767 | 0.6815 | 0.6742 | 0.7332 | 0.7528 | 0.7996 | 0.6945 |
| (-1,2) | 0.7026 | 0.5305 | 0.5056 | 0.5637 | 0.5943 | 0.6764 | 0.5211 |
| (-5,1) | 0.368 | 0.7185 | 0.7512 | 0.7907 | 0.7797 | 0.9058 | 0.784 |
| (-5,2) | 0.4067 | 0.5416 | 0.483 | 0.5684 | 0.6157 | 0.6666 | 0.5726 |

**Supplementary Table 4**: Proportions of genetic variants that received the same rank based on the full and reduced genetic models across all variants (ALL), top 10% and top 20%. We simulated 5,000 datasets with 3,000 cases and 3,000 controls. We simulated 10 genetic variants from Bernoulli(0.1) and disease status from the full model with coefficients $\beta_{O}=log(3)$ and $\mu_{G}=log(1), log \left( 2 \right),\log\left( 3 \right),\log\left( 5 \right),log(8).$

| $\beta_{0}=-3.5,\sigma=1,\mu_{0}=0$,$\mu_{g}=log(1.5$) | | | | | | | |
| --- | --- | --- | --- | --- | --- | --- | --- |
| $\beta_{G}$ | $\mu_{d}$ | $\frac{\mu_{d}\mu_{g}}{\sigma^{2}}$ | ${pr}_{B}(D=d$) | Bias | Variance | MSE | power |
| $-\log\left( 2 \right)$ | $-\log\left( 1.5 \right)$ | -0.1644 | 0.0301 | -0.1554 | 0.0027 | 0.0268 | 1 |
| $-\log\left( 2 \right)$ | $\log\left( 1 \right)$ | 0 | 0.0279 | 0.0060 | 0.0024 | 0.0025 | 1 |
| $-\log\left( 2 \right)$ | $\log\left( 1.5 \right)$ | 0.1644 | 0.0306 | 0.1908 | 0.0022 | 0.0386 | 1 |
| $\log\left( 2 \right)$ | $-\log\left( 1.5 \right)$ | -0.1644 | 0.0340 | -0.1627 | 0.0017 | 0.0282 | 1 |
| $\log\left( 2 \right)$ | $\log\left( 1 \right)$ | 0 | 0.0322 | 0.0121 | 0.0016 | 0.0017 | 1 |
| $\log\left( 2 \right)$ | $\log\left( 1.5 \right)$ | 0.1644 | 0.0036 | 0.1729 | 0.0016 | 0.0315 | 1 |
| $\beta_{0}=-1,\sigma=1,\mu_{0}=0$,$\mu_{g}=log(1.5$) | | | | | | | |
| $\beta_{G}$ | $\mu_{d}$ | $\frac{\mu_{d}\mu_{g}}{\sigma^{2}}$ | ${pr}_{B}(D=d$) | Bias | Variance | MSE | power |
| $-\log\left( 2 \right)$ | $-\log\left( 1.5 \right)$ | -0.1644 | 0.2706 | -0.1586 | 0.0029 | 0.0281 | 1 |
| $-\log\left( 2 \right)$ | $\log\left( 1 \right)$ | 0 | 0.2570 | 0.0096 | 0.0027 | 0.0028 | 1 |
| $-\log\left( 2 \right)$ | $\log\left( 1.5 \right)$ | 0.1644 | 0.2753 | 0.1748 | 0.0025 | 0.0331 | 1 |
| $\log\left( 2 \right)$ | $-\log\left( 1.5 \right)$ | -0.1644 | 0.2966 | -0.1598 | 0.0021 | 0.0277 | 1 |
| $\log\left( 2 \right)$ | $\log\left( 1 \right)$ | 0 | 0.2838 | 0.0025 | 0.0021 | 0.0021 | 1 |
| $\log\left( 2 \right)$ | $\log\left( 1.5 \right)$ | 0.1644 | 0.3046 | 0.1655 | 0.0022 | 0.0296 | 1 |
| $\beta_{0}=-3.5,\sigma=1,\mu_{0}=0$,$\mu_{g}=log(1.5$) | | | | | | | |
| $\beta_{G}$ | $\mu_{d}$ | $\frac{\mu_{d}\mu_{g}}{\sigma^{2}}$ | ${pr}_{B}(D=d$) | Bias | Variance | MSE | FDR |
| $\log\left( 1 \right)$ | $-\log\left( 1.5 \right)$ | -0.1644 | 0.0315 | -0.1487 | 0.0020 | 0.0241 | 0.8882 |
| $\log\left( 1 \right)$ | $\log\left( 1 \right)$ | 0 | 0.0294 | 0.0118 | 0.0018 | 0.0020 | 0.0354 |
| $\log\left( 1 \right)$ | $\log\left( 1.5 \right)$ | 0.1644 | 0.0324 | 0.1632 | 0.0018 | 0.0284 | 0.9542 |
| $\beta_{0}=-3.5,\sigma=1,\mu_{0}=0$,$\mu_{g}=log(1.5$) | | | | | | | |
| $\beta_{G}$ | $\mu_{d}$ | $\frac{\mu_{d}\mu_{g}}{\sigma^{2}}$ | ${pr}_{B}(D=d$) | Bias | Variance | MSE | power |
| $\log\left( 8 \right)$ | $-\log\left( 1.5 \right)$ | -0.1644 | 0.0470 | -0.1638 | 0.0016 | 0.0285 | 1 |
| $\log\left( 8 \right)$ | $\log\left( 1 \right)$ | 0 | 0.0460 | 0.0053 | 0.0016 | 0.0016 | 1 |
| $\log\left( 8 \right)$ | $\log\left( 1.5 \right)$ | 0.1644 | 0.0525 | 0.1651 | 0.0017 | 0.0289 | 1 |
| $\beta_{0}=-1,\sigma=1,\mu_{0}=0$,$\mu_{g}=log(1.5$) | | | | | | | |
| $\beta_{G}$ | $\mu_{d}$ | $\frac{\mu_{d}\mu_{g}}{\sigma^{2}}$ | ${pr}_{B}(D=d$) | Bias | Variance | MSE | FDR |
| $\log\left( 1 \right)$ | $-\log\left( 1.5 \right)$ | -0.1644 | 0.2815 | -0.1572 | 0.0023 | 0.0270 | 0.9064 |
| $\log\left( 1 \right)$ | $\log\left( 1 \right)$ | 0 | 0.2683 | 0.0049 | 0.0022 | 0.0022 | 0.05 |
| $\log\left( 1 \right)$ | $\log\left( 1.5 \right)$ | 0.1644 | 0.2882 | 0.1712 | 0.0022 | 0.0315 | 0.959 |
| $\beta_{0}=-1,\sigma=1,\mu_{0}=0$,$\mu_{g}=log(1.5$) | | | | | | | |
| $\beta_{G}$ | $\mu_{d}$ | $\frac{\mu_{d}\mu_{g}}{\sigma^{2}}$ | ${pr}_{B}(D=d$) | Bias | Variance | MSE | power |
| $\log\left( 8 \right)$ | $-\log\left( 1.5 \right)$ | -0.1644 | 0.3294 | -0.1594 | 0.0031 | 0.0285 | 1 |
| $\log\left( 8 \right)$ | $\log\left( 1 \right)$ | 0 | 0.3162 | 0.0023 | 0.0032 | 0.0032 | 1 |
| $\log\left( 8 \right)$ | $\log\left( 1.5 \right)$ | 0.1644 | 0.3353 | 0.1650 | 0.0037 | 0.0309 | 1 |

**Supplementary Table 5**: Bias approximation obtained using (6), rate of the disease in the population ${pr}_{B}(D=d$), bias, variance and mean squared error (MSE) of the estimates obtained from the reduced model. We simulated 5,000 datasets with 3,000 cases and 3,000 controls. We simulated genotype from Bernoulli(0.1), then assumed $\mu_{0}=0,\mu_{g}=\log\left( 1.5 \right),\mu_{d}=-\log\left( 1.5 \right),log(1.5),\sigma^{2}=1$,$\beta_{0}=-1,-3.5,\beta_{G}=-\log\left( 2.5 \right),-\log\left( 1.5 \right),\log\left( 1.5 \right),log(2.5)$.

B. **Analyses of ADNI dataset**

We defined case-control status to indicate a case for diagnosis {AD,LMCI,EMCI} and controls for {SN, SMC}.

We first present a detailed description of the distribution of key variables (age, gender, education, race/ethnicity, ApoE e4 status, MMSE, hippocampus volume, whole brain volume and the ratio between hippocampus volume and whole brain volume) by case-control status in ADNI data in **Supplementary Tables 7A-L**. Next, **Supplementary Table 7M** presents p-values for comparing these variables between cases and controls derived by fitting a univariable logistic regression model for continuous variables and using Fisher’s exact test for categorical variables.

**Supplementary Table 8** presents estimates of log(OR), their standard errors (SE) and p-values for various settings of the assumed full (true) model and reduced models.

There are totally 2,438 SNPs have minor allele frequency of >5%. We fitted univariate logistic regression with AD,LMCI,EMCI vs CN,SMC as the binary outcome and each SNP as the predictor and found totally 114 SNPs with p-value<0.05.

We now assume that the full (true) model is the model that includes

**Model 3:** age, gender, education, ApoE $\varepsilon$4 status, MMSE and the ratio between hippocampus and whole brain volumes and each of the SNPs.

Shown on **Supplementary Figures 1-3** are histograms of the estimates of log(OR)s and shown on **Supplementary Figure 4** is the histogram of p-values for the estimates of SNPs. We next estimated **models 3A** that omit the ratio between the brain volumes. Shown on **Supplementary Figures 5-6** are the histograms of the estimates of log(OR)s and on **Supplementary Figure 7** is the histogram of p-values for the log(OR)s of SNPs.

Shown in **Supplementary Table 11** are the differences in log(OR) estimates obtained in the full model 3 vs. reduced model 3A. Recall that model 3 includes age, male, education, ApoE $\varepsilon$4 status, MMSE and the ratio between hippocampus and whole brain volumes and each of the 2,438 SNPs; and model 3A omits the ratio between hippocampus volume and whole brain volume. We note that the average difference in the log(OR) estimates of age is -0.03, in the estimates of sex is -0.21, education: -0.02, ApoE $\varepsilon$4 status: -0.27, MMSE: .05, and SNPs: 0.006. The minimum difference in the estimates for SNPs is -0.35, while the maximum is 0.18. **Supplementary Figure 8** displays histogram of the differences.

We furthermore compare the distribution of log(OR) estimates for SNPs obtained in Model 3 vs. 3A using Kolmogorov-Smirnov test and results are shown in **Supplementary Table 12.** Those distributions are different for age, sex, education ApoE $\varepsilon$4 status, MMSE, but not SNPs. P-values shown in **Supplementary Table 13** for Kolmogorov-Smirnov test comparing the distribution of p-values for SNPs within model 3 and within model 3A show that the distributions are not different from Uniform(0,1).

Among these 2,438 SNPs, totally 105 SNPs have p-values <0.05 in the full model

According to the p-values, the most 50 significant SNPs are

"rs3017528_A" "rs220229_A"	"rs2158354_C" "rs2830257_C" "rs2830451_A" "rs10516828_C" "rs2830292_C" "rs10740244_A" "rs919001_A"	"rs768952_A"	"rs2444757_A" "rs7863087_C" "rs1918733_G" "rs4456451_A" "rs3843671_T" "rs2777864_T" "rs220179_A"	"rs2282070_G" "rs12248673_T" "rs1893577_T"

"rs2160422_T" "rs107017_G"	"rs12773412_G" "rs2251540_A" "rs5760066_A"

"rs1296028_G" "rs458358_A"	"rs27566_C"	"rs255856_C"	"rs2182621_A"

"rs7138085_A" "rs10521438_A" "rs1927113_T" "rs2240219_A" "rs2774279_T"

"rs17638883_C" "rs4912732_G" "rs6480412_T" "rs7501783_G" "rs219661_G"

"rs11145477_A" "rs2837960_G" "rs2373383_G" "rs466815_G"	"rs951629_A"

"rs896994_C"	"rs7309381_G" "rs2411899_T" "rs2241494_G" "rs914185_T"

Moreover, there are 4 SNPs have p-values <0.001, that is,

"rs2830257_C" "rs3017528_A" "rs2158354_C" "rs220229_A"

In the reduced model, among these 2,438 SNPs, totally 120 SNPs have p-values <0.05. According to the p-values, the most 50 significant SNPs are

"rs220229_A"	"rs3017528_A" "rs10740244_A" "rs2830257_C" "rs919001_A"	"rs10516828_C" "rs1918733_G" "rs2158354_C" "rs2411899_T" "rs2830451_A" "rs2837960_G" "rs2291929_C" "rs4912732_G" "rs220179_A"	"rs2282070_G" "rs2830292_C" "rs10832908_G" "rs466815_G" "rs7221333_G" "rs2777864_T"

"rs222151_T"	"rs867366_G"	"rs4334084_G" "rs11138566_T" "rs12773412_G"

"rs27566_C"	"rs255856_C"	"rs12248673_T" "rs11680429_A" "rs3843671_T"

"rs7128017_T" "rs10521438_A" "rs1927113_T" "rs107017_G"	"rs17638883_C" "rs1893590_C" "rs6825804_T" "rs2444757_A" "rs750445_C"	"rs4456451_A" "rs867595_C"	"rs2839440_T" "rs2032257_A" "rs915841_C"	"rs915843_T"	"rs755736_A"	"rs2032258_C" "rs2160422_T" "rs8992_G"	"rs8133852_A"

Moreover, there are 4 SNPs have p-values <0.001, that is,

"rs2830257_C" "rs3017528_A" "rs2158354_C" "rs220229_A"

Among the top 10 significant SNPs, 80% of the SNPs are the same, among the top 30

significant SNPs, 60% of the SNPs are the same and among the top 50 significant SNPs, 58% of the SNPs are the same. Hence overall, the conclusion about what SNPs should be carried to the validation set would be different based on these two models.

| Age | N | Mean | Std | Min | Median | Max |
| --- | --- | --- | --- | --- | --- | --- |
| Control(CN,SMC) | 192 | 75.41 | 4.91 | 59.9 | 74.8 | 89.6 |
| Case(AD,LMCI,EMCI) | 423 | 74.29 | 7.4 | 54.4 | 74.5 | 90.9 |

**Supplementary Table 7A**: Age distribution among cases and controls in ADNI data

| Gender | Male | Female |
| --- | --- | --- |
| Control(CN,SMC) | 104(54.17%) | 88(45.83%) |
| Case(AD,LMCI,EMCI) | 260(61.47%) | 163(38.53%) |

**Supplementary Table 7B:** Gender distribution among cases and controls in ADNI data

| Education | 4 | 6 | 7 | 8 | 9 | 10 | 11 | 12 | 13 | 14 | 15 | 16 | 17 | 18 | 19 | 20 |
| --- | --- | --- | --- | --- | --- | --- | --- | --- | --- | --- | --- | --- | --- | --- | --- | --- |
| Control(CN,SMC) | 0 | 1 | 1 | 1 | 1 | 5 | 0 | 13 | 12 | 20 | 10 | 45 | 13 | 35 | 8 | 27 |
| Case(AD,LMCI,EMCI) | 1 | 2 | 2 | 8 | 3 | 12 | 7 | 63 | 24 | 47 | 13 | 99 | 14 | 74 | 14 | 40 |

**Supplementary Table 7C:** Gender distribution among cases and controls in ADNI data

| Education | N | Mean | Std | Min | Median | Max |
| --- | --- | --- | --- | --- | --- | --- |
| Control(CN,SMC) | 192 | 16.07 | 2.84 | 6 | 16 | 20 |
| Case(AD,LMCI,EMCI) | 423 | 15.27 | 3.1 | 4 | 16 | 20 |

**Supplementary Table 7D:** Education distribution among cases and controls in ADNI data

| Ethnicity | Not Hisp/Latino | Hisp/Latino |
| --- | --- | --- |
| Control(CN,SMC) | 190(98.96%) | 2(1.04%) |
| Case(AD,LMCI,EMCI) | 409(96.69%) | 14(3.31%) |

**Supplementary Table 7E**: Ethnicity distribution among cases and controls in ADNI data

| Race | Am Indian/Alaskan | Asian | Black | More than one | White |
| --- | --- | --- | --- | --- | --- |
| Control(CN,SMC) | 0 | 1(0.52%) | 14(7.29%) | 0 | 177(92.19%) |
| Case(AD,LMCI,EMCI) | 1(0.24%) | 8(1.89%) | 17(4.02%) | 3(0.71%) | 394(93.14%) |

**Supplementary Table 7F**: Race distribution among cases and controls in ADNI data

| ApoE4 | 0 | 1 |
| --- | --- | --- |
| Control(CN,SMC) | 143(74.48%) | 49(25.52%) |
| Case(AD,LMCI,EMCI) | 161(38.06%) | 262(61.94%) |

**Supplementary Table 7G**: ApoE e4 status distribution among cases and controls in ADNI data

| MMSE | N | Mean | Std | Min | Median | Max |
| --- | --- | --- | --- | --- | --- | --- |
| Control(CN,SMC) | 192 | 29.07 | 1.01 | 25 | 29 | 30 |
| Case(AD,LMCI,EMCI) | 423 | 25.95 | 2.57 | 20 | 26 | 30 |

**Supplementary Table 7H**: MMSE distribution among cases and controls in ADNI data

| Hippocampus volume | N | Mean | Std | Min | Median | Max |
| --- | --- | --- | --- | --- | --- | --- |
| Control(CN,SMC) | 192 | 7264.2 | 895.59 | 5264 | 7347 | 10769 |
| Case(AD,LMCI,EMCI) | 423 | 6138.97 | 1120.68 | 3091 | 6102 | 9572 |

**Supplementary Table 7I**: Hippocampus volume distribution among cases and controls in ADNI data

| Whole brain volume | N | Mean | Std | Min | Median | Max |
| --- | --- | --- | --- | --- | --- | --- |
| Control(CN,SMC) | 192 | 1015446 | 94334.91 | 746249 | 1018030 | 1229740 |
| Case(AD,LMCI,EMCI) | 423 | 985409.6 | 114475.2 | 738813 | 979010 | 1364690 |

**Supplementary Table 7J**: Whole brain volume distribution among cases and controls in ADNI data

| Hippocampus volume/ Whole brain volume | N | Mean | Std | Min | Median | Max |
| --- | --- | --- | --- | --- | --- | --- |
| Control(CN,SMC) | 192 | 0.007 | 0.001 | 0.005 | 0.007 | 0.01 |
| Case(AD,LMCI,EMCI) | 423 | 0.006 | 0.001 | 0.004 | 0.006 | 0.01 |

**Supplementary Table 7K**: Ratio of hippocampus volume to whole brain volume distribution among cases and controls in ADNI data

|  | Age | male | education | ethnicity | race |
| --- | --- | --- | --- | --- | --- |
| P-value | 0.058 | 0.09 | 0.003 | 0.17 | 0.19 |
|  | ApoE4 | MMSE | Hippocampus volume | Whole brain volume | Hippocampus volume/ Whole brain volume |
| P-value | <0.001 | <0.001 | <0.001 | 0.002 | <0.001 |

**Supplementary Table 7M**: P-values for comparing the characteristics between cases and controls in ADNI data

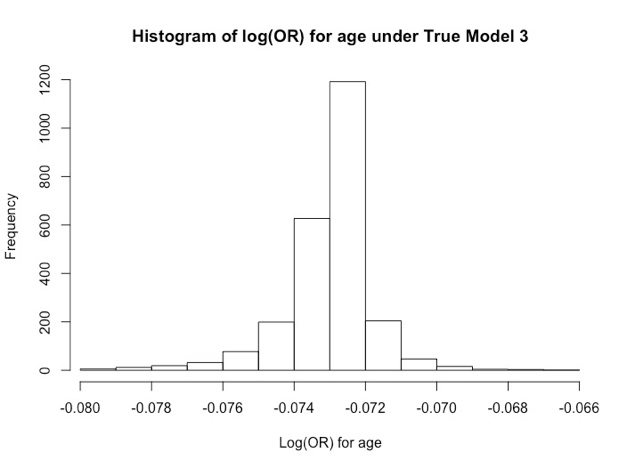

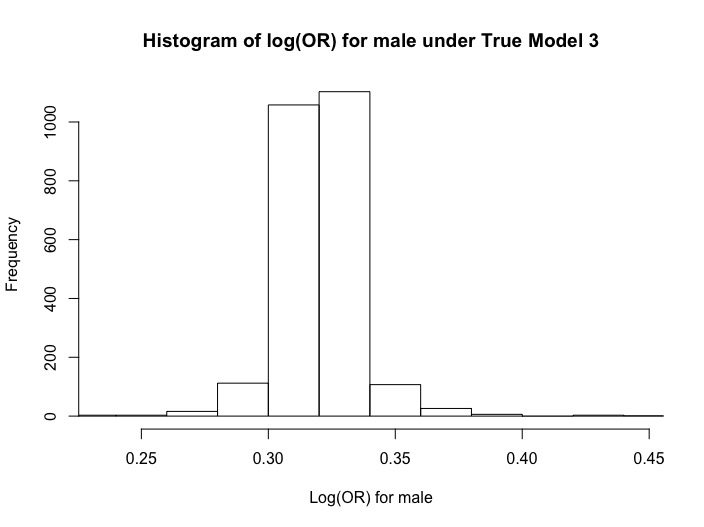

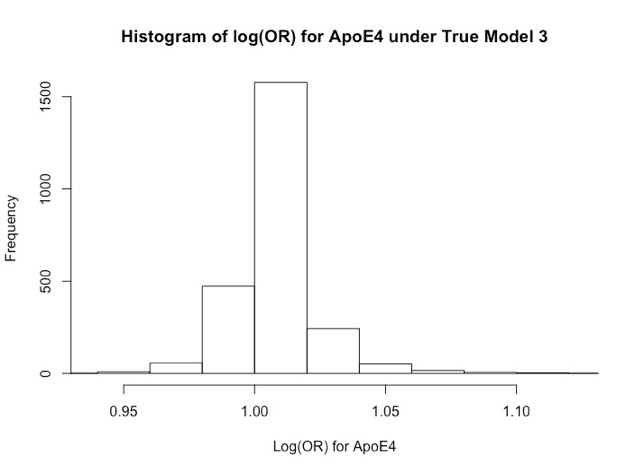

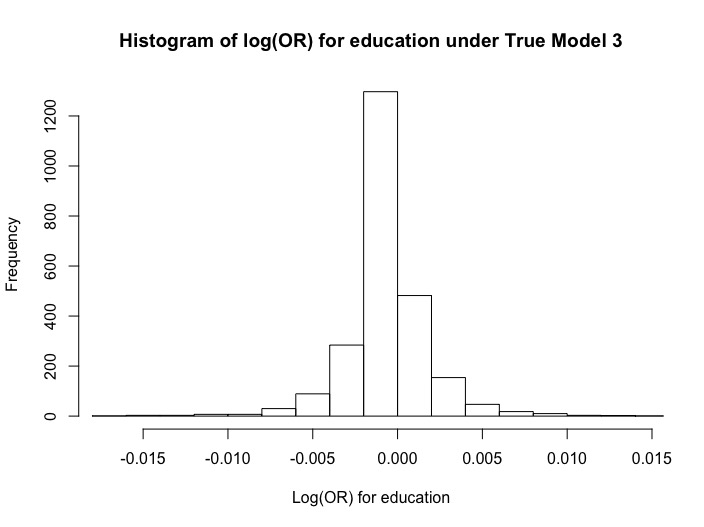

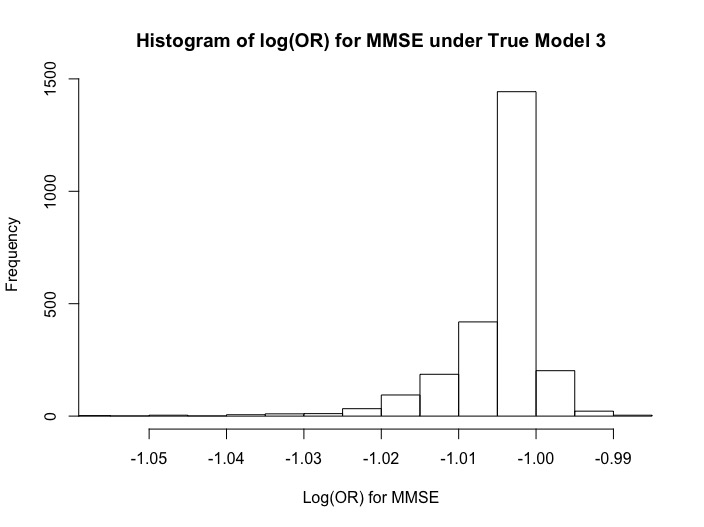

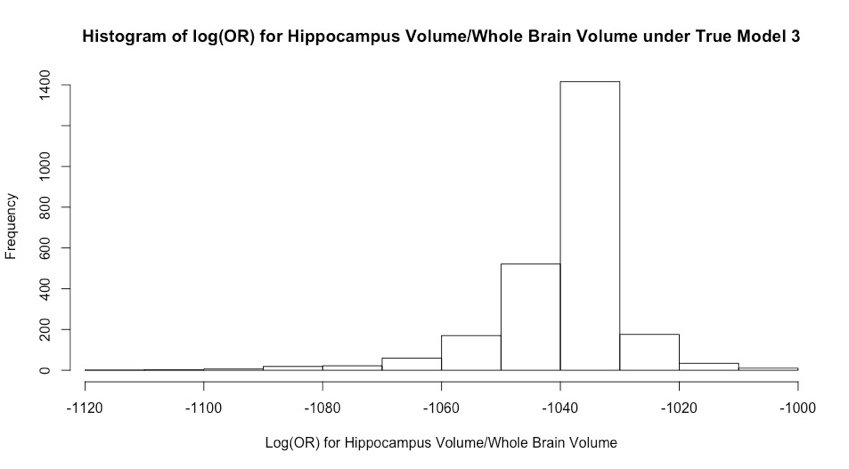

**Supplementary Figure 1**: Histograms of log(OR) estimates for age,male, ApoE $\varepsilon$4 status, education and MMSE, the ratio between hippocampus volume and whole brain volume across true models 3 that include age, male, education, ApoE $\varepsilon$4 status, MMSE and the ratio between hippocampus and whole brain volumes and each of the 2,438 SNPs.

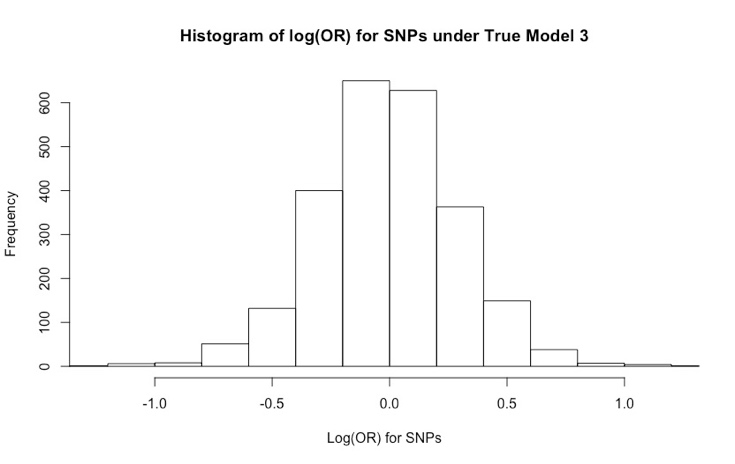

**Supplementary Figure 2:** Histograms of log(OR) estimates for SNPs across true models 3 that include age, male, education, ApoE $\varepsilon$4 status, MMSE and the ratio between hippocampus and whole brain volumes and each of the 2,438 SNPs.

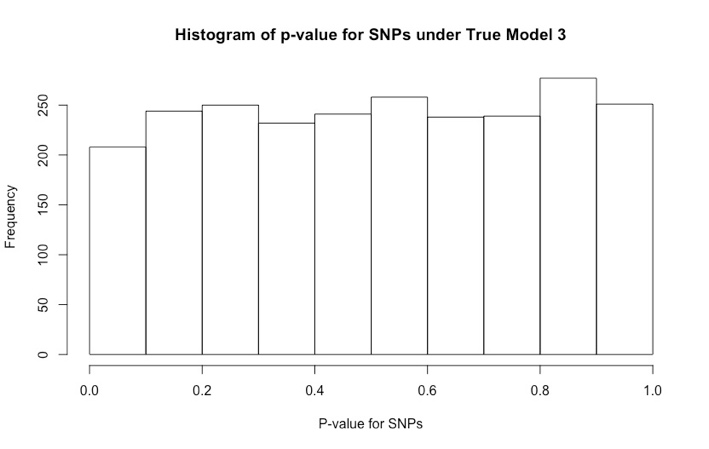

**Supplementary Figure 3:** Histograms of p-values for log(OR) estimates for SNPs across true models 3 that include age, male, education, ApoE $\varepsilon$4 status, MMSE and the ratio between hippocampus and whole brain volumes and each of the 2,438 SNPs.

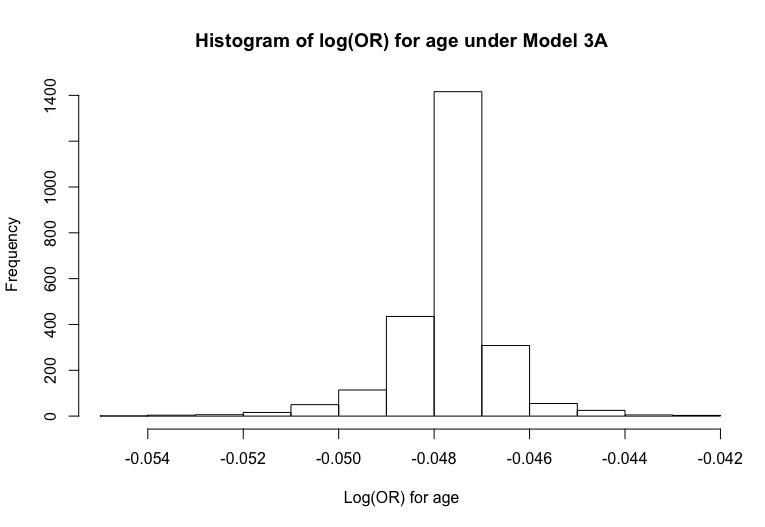

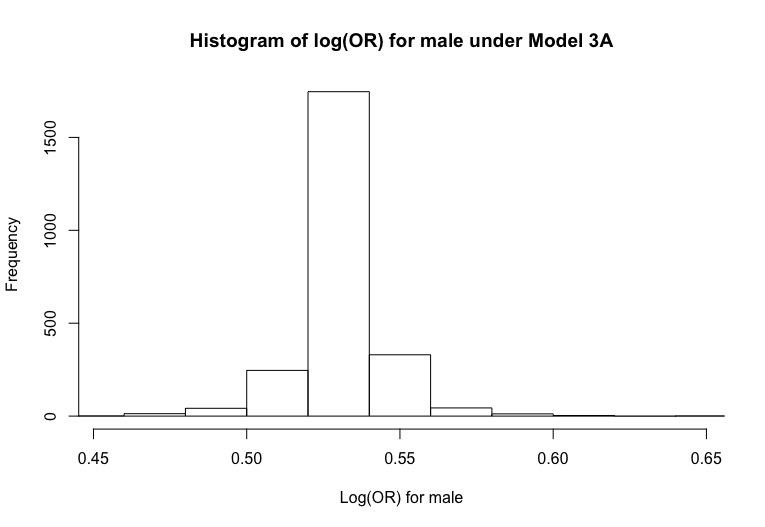

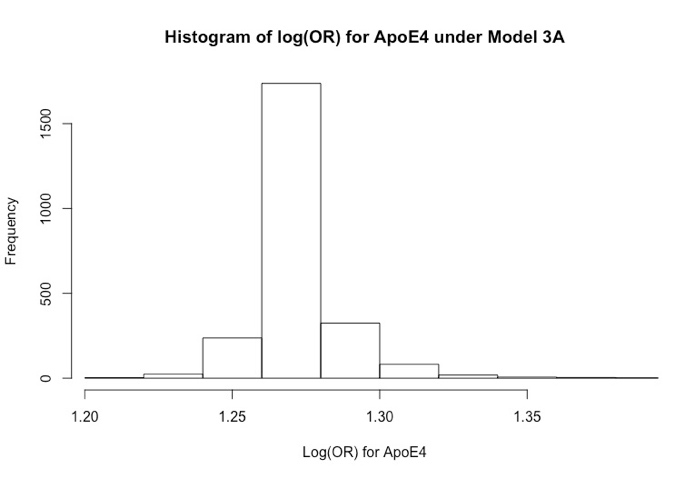

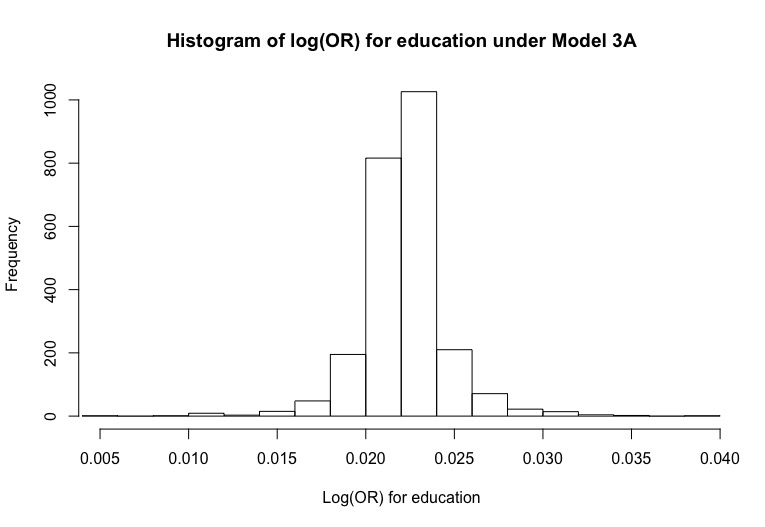

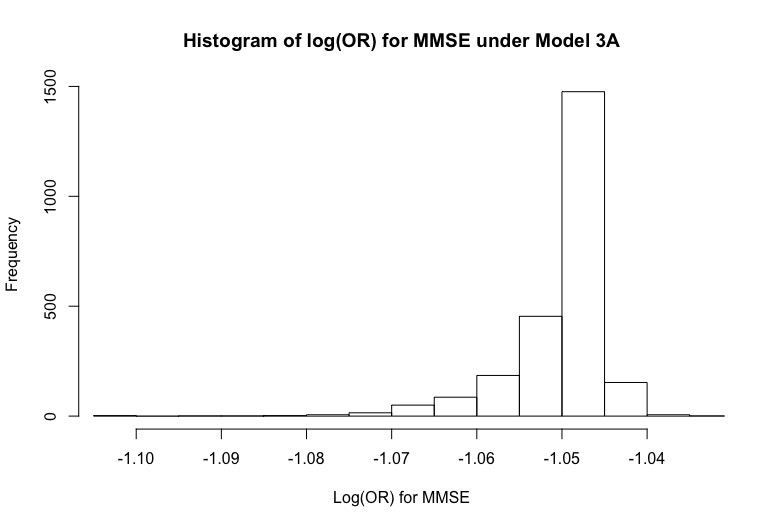

**Supplementary Figure 4**: Histograms of log(OR) estimates for age, male, ApoE $\varepsilon$4 status, education and MMSE across model that that include age, male, education, ApoE $\varepsilon$4 status, MMSE and omits the ratio between hippocampus and whole brain volumes and each of the 2,438 SNPs.

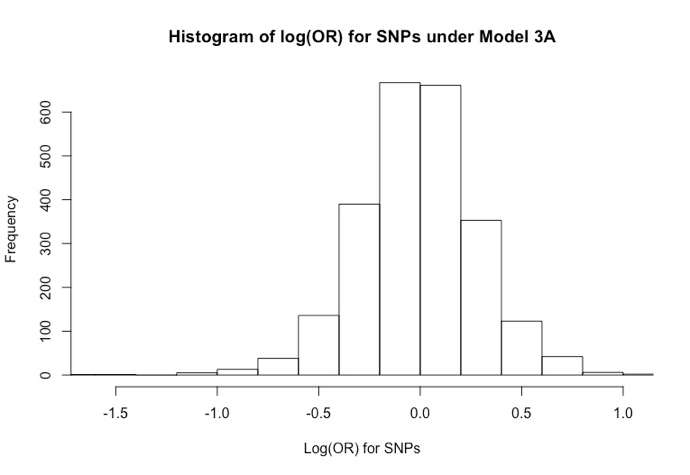

**Supplementary Figure 5:** Histograms of log(OR) estimates for SNPs across models that include age, male, education, ApoE $\varepsilon$4 status, MMSE and omits the ratio between hippocampus and whole brain volumes and each of the 2,438 SNPs.

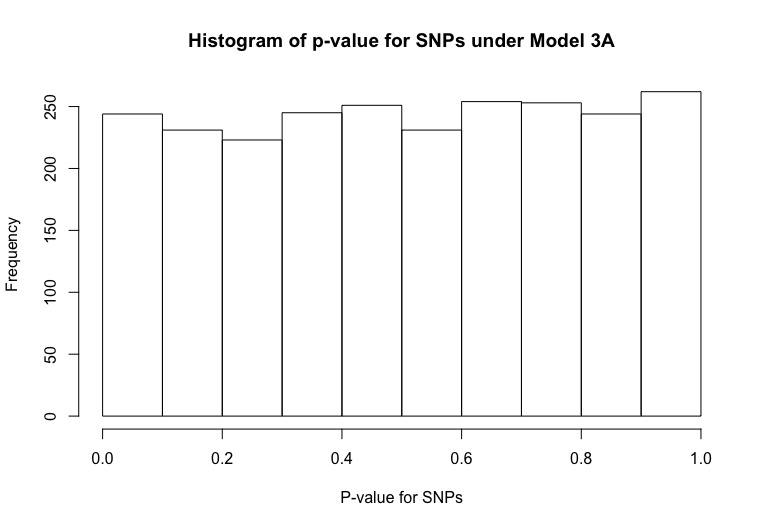

**Supplementary Figure 6:** Histograms of p-values for log(OR) estimates for SNPs across true models 3 that include age, male, education, ApoE $\varepsilon$4 status, MMSE and the ratio between hippocampus and whole brain volumes and each of the 2,438 SNPs.

|  | Age | Male | Education | ApoE4 | MMSE | Hippocampus Volume/Whole Brain Volume |
| --- | --- | --- | --- | --- | --- | --- |
| **Full Model 1: Age, male, education, ApoE4, MMSE** | | | | | | |
| Log(OR) | **-0.048** | **0.531** | **0.022** | **1.27** | **-1.047** |  |
| SE | 0.02 | 0.255 | 0.043 | 0.25 | 0.098 |  |
| p-value | 0.017 | 0.037 | 0.603 | <0.001 | <0.001 |  |
| Reduced Model 1A: omits variable MMSE | | | | | | |
| Log(OR) | **-0.021** | **0.499** | **-0.098** | **1.52** |  |  |
| SE | 0.014 | 0.196 | 0.033 | 0.196 |  |  |
| p-value | 0.145 | 0.011 | 0.003 | <0.001 |  |  |
| Reduced Model 1B: omits variable ApoE4 | | | | | | |
| Log(OR) | **-0.058** | **0.514** | **0.014** |  | **-1.069** |  |
| SE | 0.019 | 0.247 | 0.042 |  | 0.096 |  |
| p-value | 0.003 | 0.038 | 0.732 |  | <0.001 |  |
| **Full Model 2: Age, male, education, ApoE4, MMSE, Hippocampus Volume/Whole Brain Volume** | | | | | | |
| Log(OR) | **-0.073** | **0.320** | **-0.001** | **1.005** | **-1.002** | **-1036.353** |
| SE | 0.021 | 0.271 | 0.045 | 0.268 | 0.104 | 174.014 |
| p-value | 0.001 | 0.237 | 0.991 | <0.001 | <0.001 | <0.001 |
| Reduced Model 2A: Omits Hippocampus Volume/Whole Brain Volume | | | | | | |
| Log(OR)S | **-0.048** | **0.531** | **0.022** | **1.270** | **-1.047** | **-0.048** |
| SE | 0.020 | 0.255 | 0.043 | 0.250 | 0.098 | 0.020 |
| p-value | 0.017 | 0.037 | 0.603 | <0.001 | <0.001 | 0.017 |

**Supplementary Table 8**: Estimates of log(OR), their standard errors (SE) and p-values for various settings of the assumed full (true) model and various reduced models in ADNI data.

| Variables | Min | 1^st^ Qu. | Median | Mean | 3^rd^ Qu. | Max | p-value |
| --- | --- | --- | --- | --- | --- | --- | --- |
| Age | -0.0248 | -0.0254 | -0.0253 | -0.0254 | -0.0252 | -0.0238 | <0.001 |
| Male | -0.2191 | -0.2113 | -0.2112 | -0.2108 | -0.2110 | -0.2010 | <0.001 |
| Education | -0.0227 | -0.0227 | -0.0226 | -0.0227 | -0.0227 | -0.0245 | <0.001 |
| ApoE4 | -0.2655 | -0.2658 | -0.2653 | -0.2655 | -0.2647 | -0.2632 | <0.001 |
| MMSE | 0.0476 | 0.0450 | 0.0452 | 0.0452 | 0.0454 | 0.0474 | <0.001 |
| SNPs | 0.3512 | -0.001 | -0.0015 | 0.0064 | 0.0046 | 0.1758 | 0.4424 |

**Supplementary Table 11**: Summary statistics on the difference in log(OR) estimates obtained in the full model 3 and reduced model 3A. Model 3 includes age, male, education, ApoE $\varepsilon$4 status, MMSE and the ratio between hippocampus and whole brain volumes and each of the 2,438 SNPs; and model 3A omits the ratio between hippocampus volume and whole brain volume.

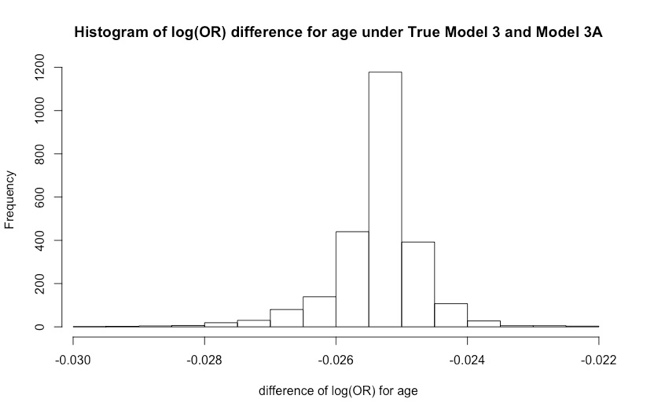

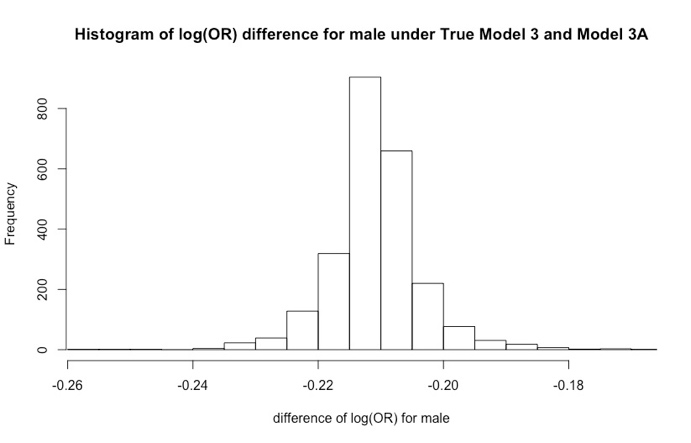

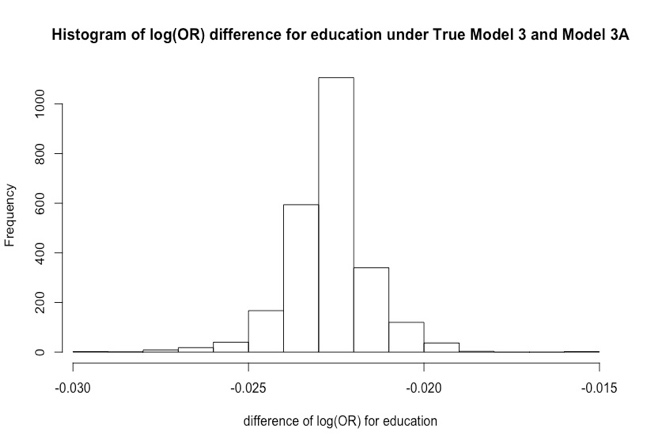

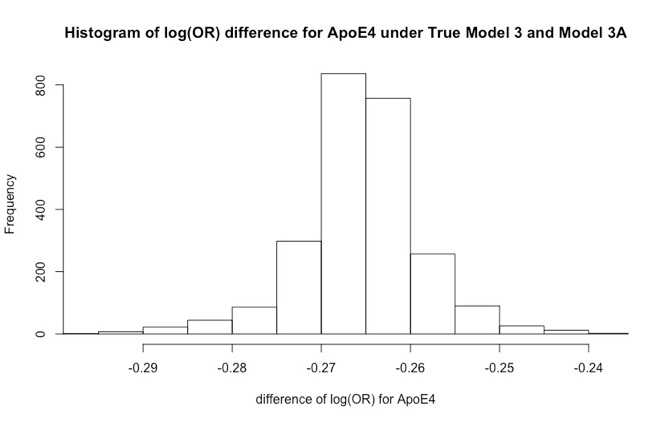

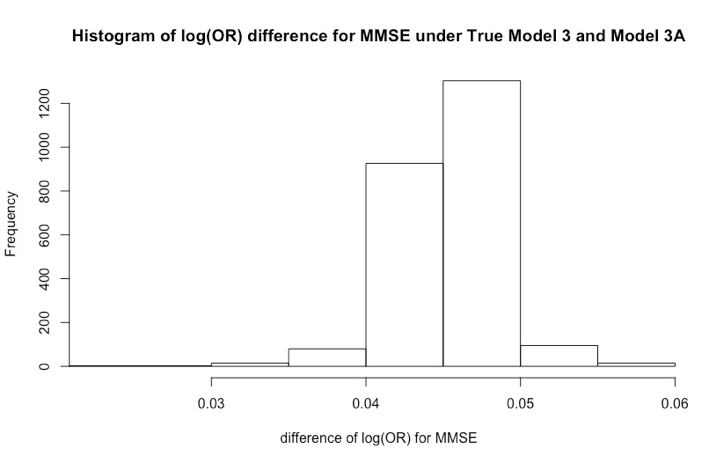

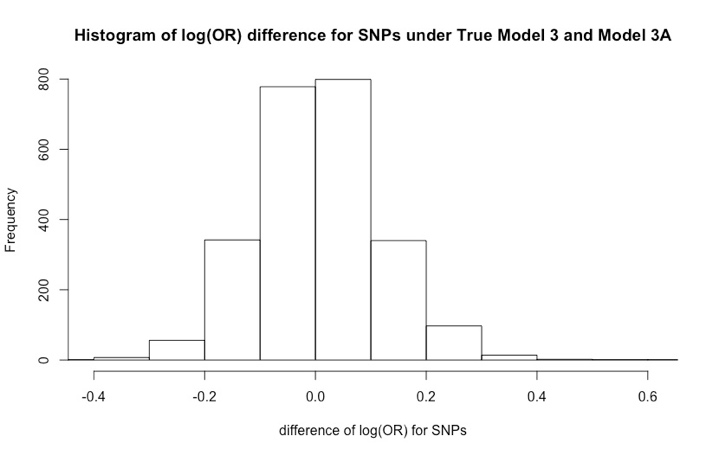

**Supplementary Figure 8**: Histograms of the difference in log(OR) estimates between model 3 that includes age, male, education, ApoE $\varepsilon$4 status, MMSE and the ratio between hippocampus and whole brain volumes and each of the 2,438 SNPs; and model 3A omits the ratio between hippocampus volume and whole brain volume.

We used the Kolmogorov-Smirnov test (two.sided) to compare the distributions of log(OR) estimates for each variable in full model 3 and reduced model 3A.

| Variables | Age | Male | Education | ApoE4 | MMSE | SNPs |
| --- | --- | --- | --- | --- | --- | --- |
| p-value | <0.001 | <0.001 | <0.001 | <0.001 | <0.001 | 0.316 |

**Supplementary Table 12:** Two-sided p-values from Kolmagorov-Smirnov test comparing the distributions of log(OR) estimates for each variable obtained from the full model 3 vs. reduced model 3A. Model 3 that includes age, male, education, ApoE $\varepsilon$4 status, MMSE and the ratio between hippocampus and whole brain volumes and each of the 2,438 SNPs; and model 3A omits the ratio between hippocampus volume and whole brain volume.

|  | True Model 3 | Model 3A |
| --- | --- | --- |
| p-value | 0.17 | 0.4234 |

**Supplementary Table 13:** Two-sided p-values from Kolmagorov-Smirnov test comparing the distributions of p-values estimates for SNPs obtained from the full model 3 and for reduced model 3A to the Uniform(0,1). Model 3 that includes age, male, education, ApoE $\varepsilon$4 status, MMSE and the ratio between hippocampus and whole brain volumes and each of the 2,438 SNPs; and model 3A omits the ratio between hippocampus volume and whole brain volume.

C. **Analyses of ADGC dataset**

In what follows we describe the ADGC dataset. **Supplementary Tables 14A-E** describe age, sex, education, ApoE $\varepsilon$4 status that is available on a subset. **Supplementary Table 15** shows estimates for the full model 1 that includes age, male, education and ApoE $\varepsilon$4 status and the reduced models that omit age, sex, education, ApoE $\varepsilon$4 status. We note that omitting ApoE $\varepsilon$4 status changes the estimates by more than omitting the other variables.

**Supplementary Figure 9** presents histograms of log(OR) estimates for age, sex, ApoE $\varepsilon$4 status across the full models 2 that include age, male, education and ApoE $\varepsilon$4 status and each of the 157 SNPs. **Supplementary Figure 10** shows histograms of log(OR) estimates for SNPs across the full models 2. **Supplementary Figure 11** shows histogram of p-values for log(OR) estimates for SNPs across true models 2 that include age, male, education, ApoE $\varepsilon$4 status and each of the 157 SNPs. **Supplementary Figure 12** Histograms of log(OR) estimates for age, male and education across model that that include age, male, education and each of the 157 SNPs. **Supplementary Table 13** shows histogram of log(OR) estimates in a model with age, sex, education and each of the SNPs. And **Supplementary Table 14** shows p-values for log(OR) estimates in model 2.

**Supplementary Table 16** shows the summary statistics of the differences in log(OR) estimates between model 2 that includes age, sex, education, ApoE $\varepsilon$4 status and each of the 157 SNPs, while model 2A omits ApoE $\varepsilon$4 status. The difference between log(OR) estimates is on average 0.02 for age, 0.03 for sex, 0.005 for education, 0.01 for SNPs.

**Supplementary Table 17** shows p-values comparing the distribution of log(OR) estimates between models 2 and 2A, showing that it is significant for age, sex, education. P-values testing the difference in the distribution of p-values for SNPs vs. Uniform(0,1) shown in **Supplementary Table 18** demonstrate that the distribution of p-values does not deviate from Uniform(0,1).

**Supplementary Figure 15** presents histograms of the difference in the log(OR) estimates between full model 2 and reduced model 2A.

| Age | N | Mean | Std | Min | Median | Max |
| --- | --- | --- | --- | --- | --- | --- |
| Control | 332 | 75.19 | 8.27 | 58 | 75 | 98 |
| Case | 2055 | 70.78 | 8.82 | 45 | 71 | 106 |

**Supplementary Table 14A**: Age distribution among cases and controls in ADGC data

| Gender | Male | Female |
| --- | --- | --- |
| Control | 109 (32.8%) | 223 (67.2%) |
| Case | 1005 (48.9%) | 1050 (51.1%) |

**Supplementary Table 14B**: Gender distribution among cases and controls in ADGC data

| Education | N | Mean | Std | Min | Median | Max |
| --- | --- | --- | --- | --- | --- | --- |
| Control | 332 | 15.93 | 2.72 | 0 | 16 | 25 |
| Case | 2055 | 14.08 | 3.38 | 0 | 14 | 30 |

**Supplementary Table 14C**: Education distribution among cases and controls in ADGC data

| ApoE4 | 0 | 1 |
| --- | --- | --- |
| Control | 236 (71.1%) | 96 (28.9%) |
| Case | 728 (35.4%) | 1327 (64.6%) |

**Supplementary Table 14D**: ApoE e4 status distribution among cases and controls in ADGC data

|  | Age | male | education | ApoE4 |
| --- | --- | --- | --- | --- |
| P-value | <0.001 | <0.001 | <0.001 | <0.001 |

**Supplementary Table 14E**: P-values for comparing the characteristics between cases and controls in ADGC data

|  | age | male | education | ApoE4 |
| --- | --- | --- | --- | --- |
| **True Model 1: age, male, education, ApoE4** | | | | |
| Log(OR) | -0.041 | 0.989 | -0.236 | 1.392 |
| SE | 0.007 | 0.144 | 0.023 | 0.137 |
| p-value | <0.001 | <0.001 | <0.001 | <0.001 |
| **Model 1A: omits variable age** | | | | |
| Log(OR) |  | 1.038 | -0.235 | 1.515 |
| SE |  | 0.141 | 0.022 | 0.134 |
| p-value |  | <0.001 | <0.001 | <0.001 |
| **Model 1B: omits variable male** | | | | |
| Log(OR) | -0.045 |  | -0.188 | 1.411 |
| SE | 0.007 |  | 0.02 | 0.135 |
| p-value | <0.001 |  | <0.001 | <0.001 |
| **Model 1C: omits variable education** | | | | |
| Log(OR) | -0.040 | 0.547 |  | 1.355 |
| SE | 0.007 | 0.130 |  | 0.133 |
| p-value | <0.001 | <0.001 |  | <0.001 |
| **Model 1D: omits variable ApoE4** | | | | |
| Log(OR)S | -0.056 | 1.014 | -0.231 |  |
| SE | 0.007 | 0.139 | 0.022 |  |
| p-value | <0.001 | <0.001 | <0.001 |  |

**Supplementary Table 15**: Estimates of log(OR), their standard errors (SE) and p-values for various settings of the assumed full (true) model and various reduced models in ADGC data.

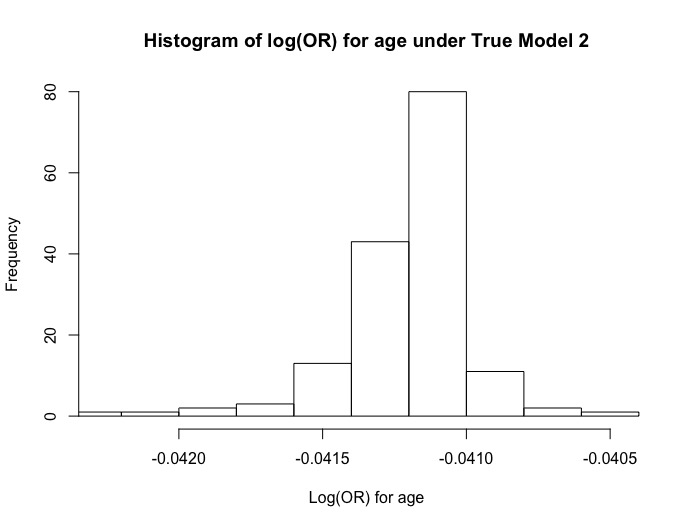

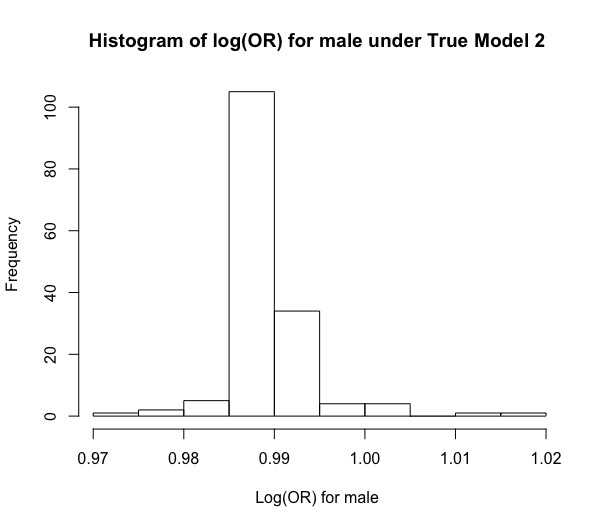

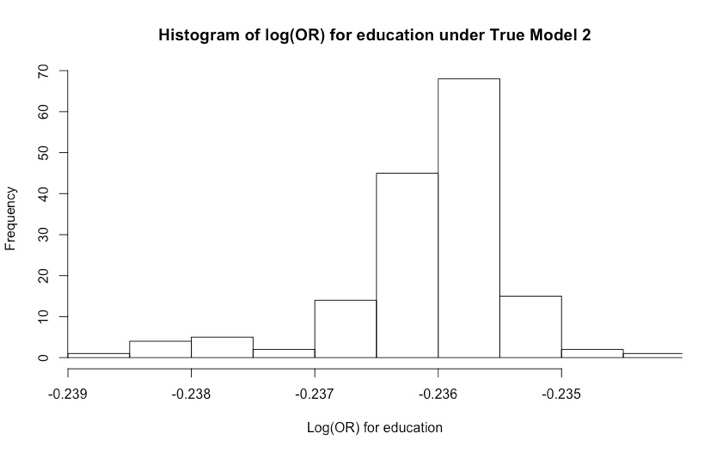

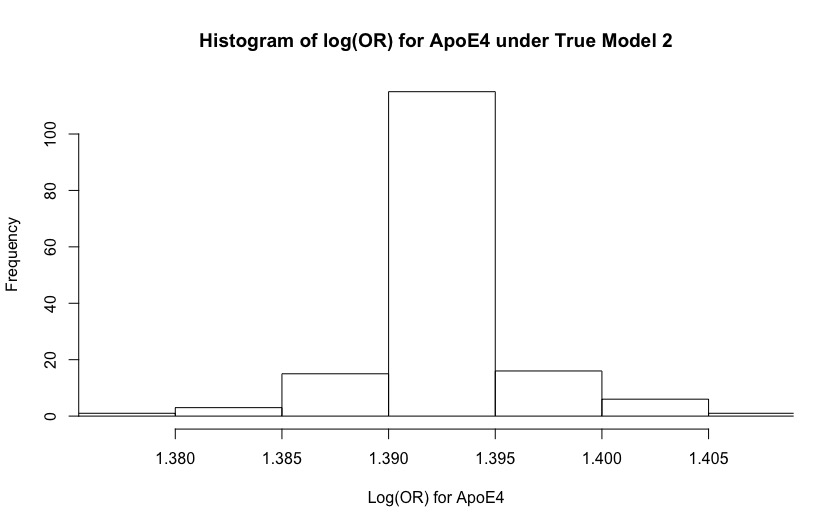

**Supplementary Figure 9**: Histograms of log(OR) estimates for age, male, education and ApoE $\varepsilon$4 status across true models 2 that include age, male, education and ApoE $\varepsilon$4 status and each of the 157 SNPs.

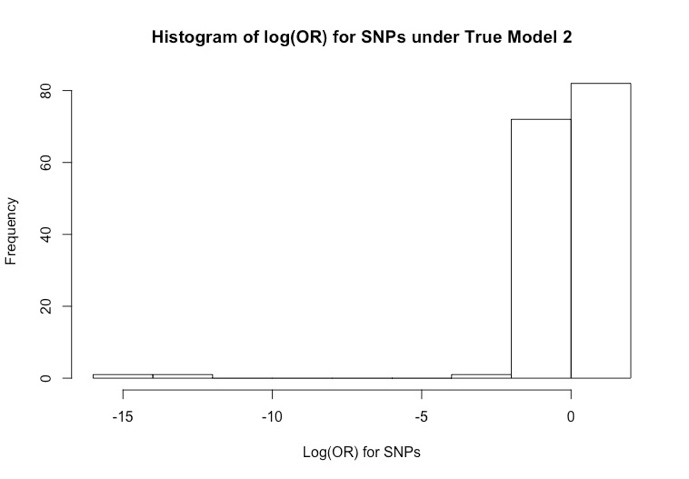

**Supplementary Figure 10:** Histograms of log(OR) estimates for SNPs across true models 2 that include age, male, education, ApoE $\varepsilon$4 status and each of the 157 SNPs.

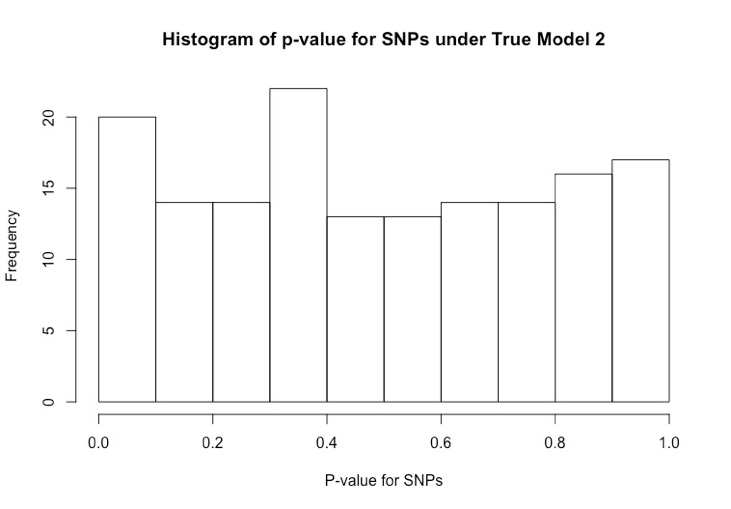

**Supplementary Figure 11:** Histograms of p-values for log(OR) estimates for SNPs across true models 2 that include age, male, education, ApoE $\varepsilon$4 status and each of the 157 SNPs.

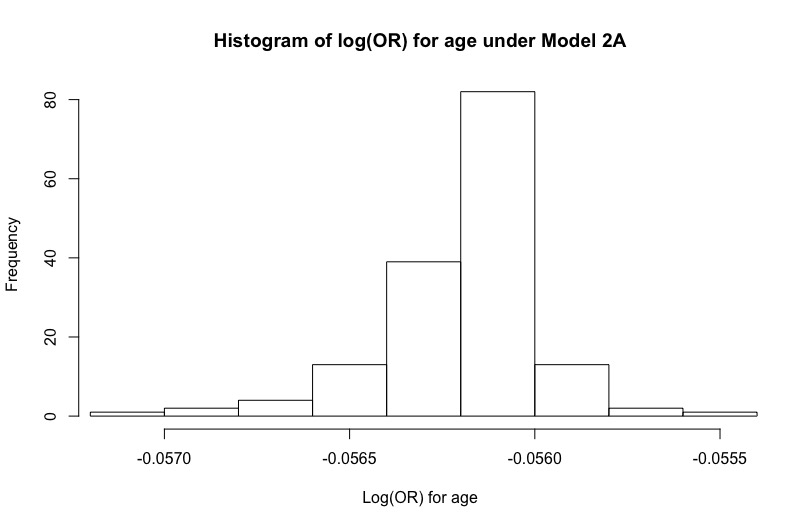

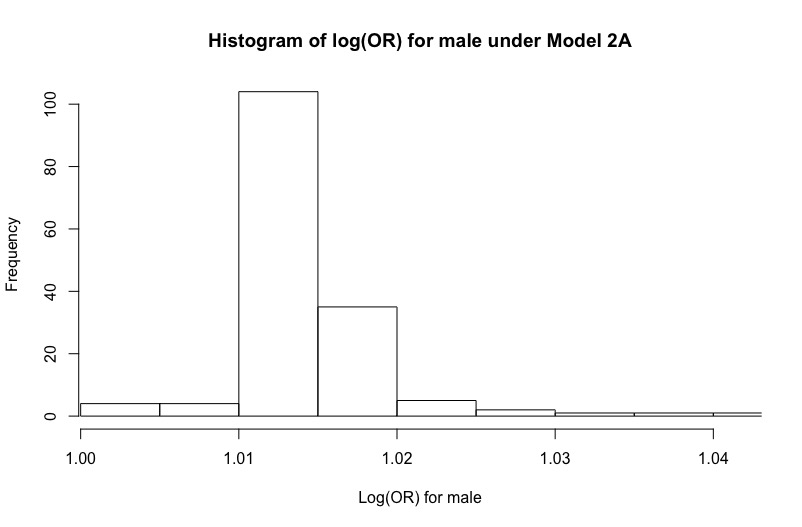

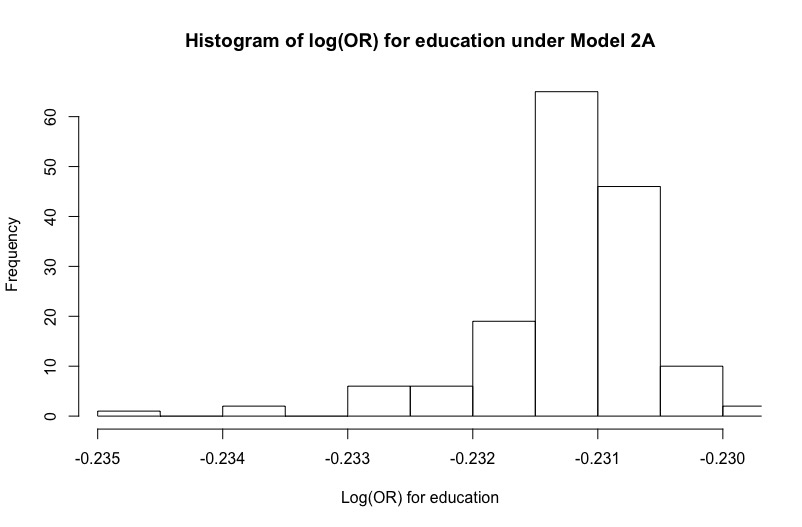

**Supplementary Figure 12**: Histograms of log(OR) estimates for age, male and education across model that that include age, male, education and each of the 157 SNPs.

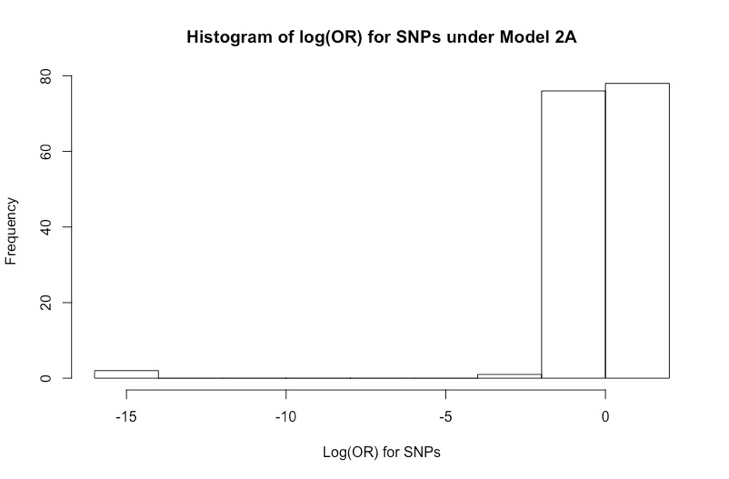

**Supplementary Figure 13:** Histograms of log(OR) estimates for SNPs across model that that include age, male, education and each of the 157 SNPs.

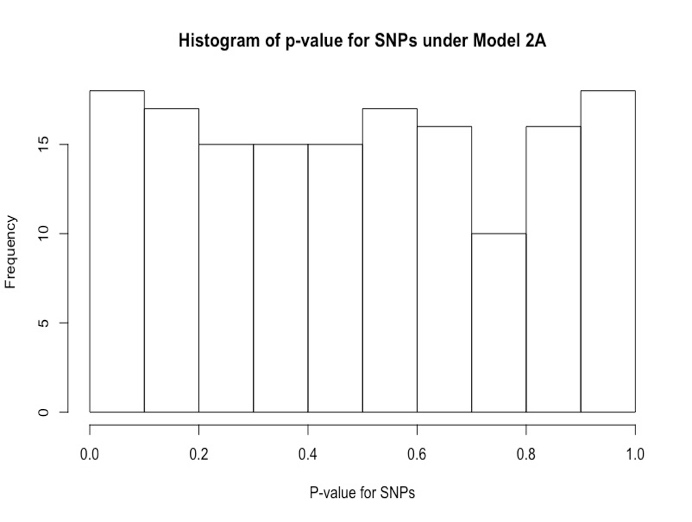

**Supplementary Figure 14:** Histograms of p-values for log(OR) estimates for SNPs across model that that include age, male, education and each of the 157 SNPs.

| Variables | Min | 25^th^ percentile | Median | Mean | 75^th^ percentile | Max | p-value |
| --- | --- | --- | --- | --- | --- | --- | --- |
| Age | 0.0147 | 0.0150 | 0.0150 | 0.0150 | 0.0150 | 0.0152 | <0.001 |
| Male | -0.0326 | -0.0258 | -0.0250 | -0.0250 | -0.0242 | -0.0190 | <0.001 |
| Education | -0.0060 | -0.0050 | -0.0049 | -0.0049 | -0.0048 | -0.0029 | <0.001 |
| SNPs | -0.1728 | -0.0174 | 0.0075 | 0.0134 | 0.0371 | 0.5770 | 0.9411 |

**Supplementary Table 16**: Summary statistics of the differences in log(OR) estimates obtained in full model 2 and reduced model 2A. Model 2 includes age, male, education, ApoE $\varepsilon$4 status and each of the 157 SNPs. Model 2A omits ApoE $\varepsilon$4 status.

| Variables | Age | Male | Education | SNPs |
| --- | --- | --- | --- | --- |
| p-value | <0.001 | <0.001 | <0.001 | 0.9977 |

**Supplementary Table 17:** P-values from Kolmogorov-Smirnov test to compare the distributions of log(OR) estimates for each variable in full model 2 vs. reduced model 2A. Model 2 includes age, male, education, ApoE $\varepsilon$4 status and each of the 157 SNPs. Model 2A omits ApoE $\varepsilon$4 status.

|  | True Model 2 | Model 2A |
| --- | --- | --- |
| p-value | 0.8072 | 0.7965 |

**Supplementary Table 18:** P-value for comparing the distribution of log(OR) estimates for SNPs within the full model 2 and reduced model 2A.

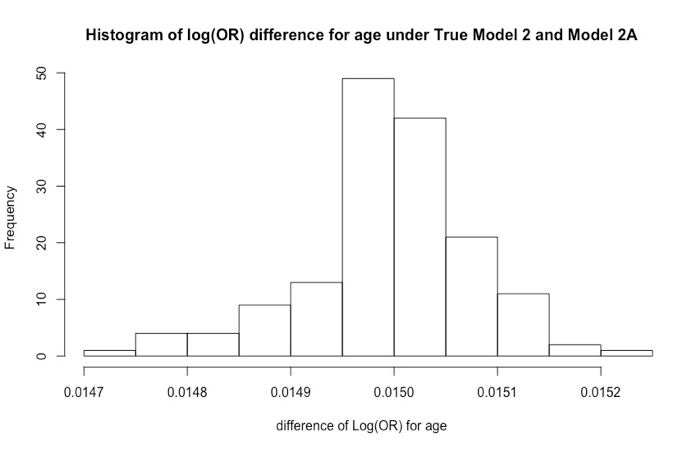

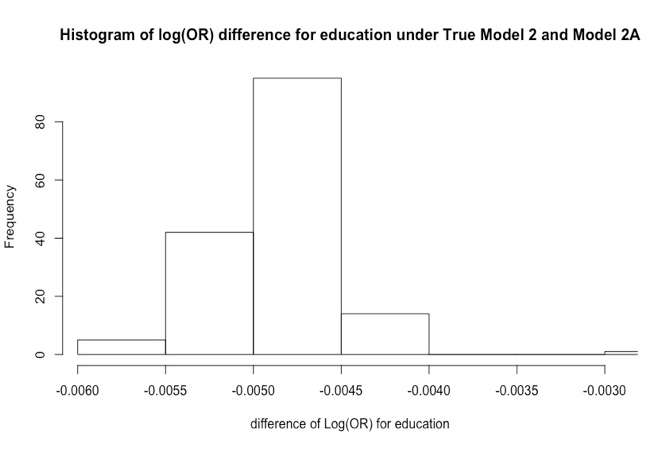

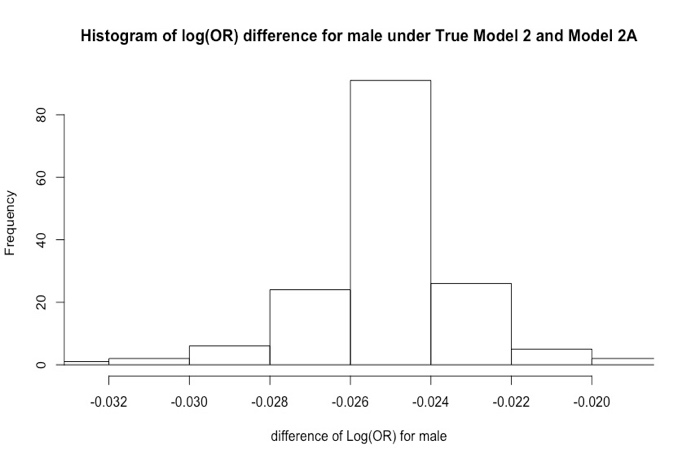

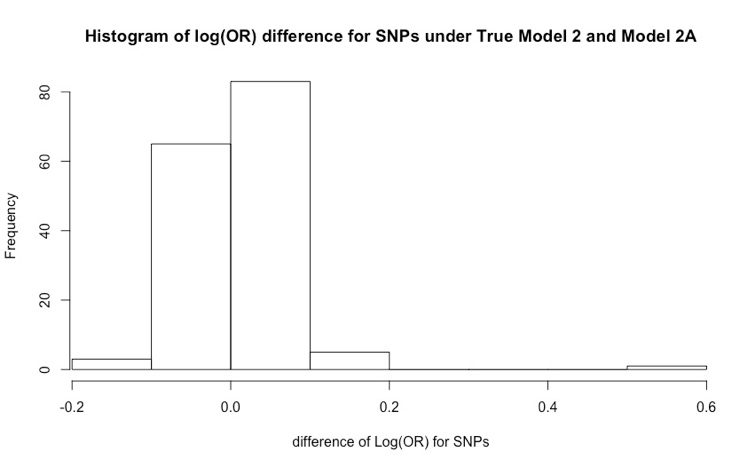

**Supplementary Figure 15:** Histogram of differences in log(OR) estimates between the full model 2 and reduced model 2A.
